# Novel genetic determinants of telomere length from a trans-ethnic analysis of 109,122 whole genome sequences in TOPMed

**DOI:** 10.1101/749010

**Authors:** Margaret A Taub, Matthew P Conomos, Rebecca Keener, Kruthika R Iyer, Joshua S Weinstock, Lisa R Yanek, John Lane, Tyne W Miller-Fleming, Jennifer A Brody, Caitlin P McHugh, Deepti Jain, Stephanie Gogarten, Cecelia A Laurie, Ali Keramati, Marios Arvanitis, Albert V Smith, Benjamin Heavner, Lucas Barwick, Lewis C Becker, Joshua C Bis, John Blangero, Eugene R Bleecker, Esteban G Burchard, Juan C Celedon, Yen Pei C Chang, Brian Custer, Dawood Darbar, Lisa de las Fuentes, Dawn L DeMeo, Barry I Freedman, Melanie E Garrett, Mark T Gladwin, Susan R Heckbert, Bertha A Hidalgo, Marguerite R Irvin, Talat Islam, W Craig Johnson, Stefan Kaab, Lenore Launer, Jiwon Lee, Simin Liu, Arden Moscati, Kari E North, Patricia A Peyser, Nicholas Rafaels, Laura M Raffield, Christine Seidman, Daniel E Weeks, Fayun Wen, Marsha M Wheeler, L. Keoki Williams, Ivana V Yang, Wei Zhao, Stella Aslibekyan, Paul L Auer, Donald W Bowden, Brian E Cade, Zhanghua Chen, Michael H Cho, L Adrienne Cupples, Joanne E Curran, Michelle Daya, Ranjan Deka, Celeste Eng, Tasha Fingerlin, Xiuqing Guo, Lifang Hou, Shih-Jen Hwang, Jill M Johnsen, Eimear E Kenny, Albert M Levin, Chunyu Liu, Ryan L Minster, Take Naseri, Mehdi Nouraie, Muagututi‘a Sefuiva Reupena, Ester C Sabino, Jennifer A Smith, Nicholas L Smith, Jessica Lasky Su, Taylor James G, Marilyn J Telen, Hemant K Tiwari, Russell P Tracy, Marquitta J White, Yingze Zhang, Kerri L Wiggins, Scott T Weiss, Ramachandran S Vasan, Kent D Taylor, Moritz F Sinner, Edwin K Silverman, M. Benjamin Shoemaker, Wayne H-H Sheu, Frank Sciurba, David Schwartz, Jerome I Rotter, Daniel Roden, Susan Redline, Benjamin A Raby, Bruce M Psaty, Juan M Peralta, Nicholette D Palmer, Sergei Nekhai, Courtney G Montgomery, Braxton D Mitchell, Deborah A Meyers, Stephen T McGarvey, Fernando D Martinez on behalf of the NHLBI CARE Network, Angel CY Mak, Ruth JF Loos, Rajesh Kumar, Charles Kooperberg, Barbara A Konkle, Shannon Kelly, Sharon LR Kardia, Robert Kaplan, Jiang He, Hongsheng Gui, Frank D Gilliland, Bruce Gelb, Myriam Fornage, Patrick T Ellinor, Mariza de Andrade, Adolfo Correa, Yii-Der Ida Chen, Eric Boerwinkle, Kathleen C Barnes, Allison E Ashley-Koch, Donna K Arnett, Christine Albert, NHLBI Trans-Omics for Precision Medicine (TOPMed) Consortium, TOPMed Hematology and Hemostasis Working Group, TOPMed Structural Variation Working Group, Cathy C Laurie, Goncalo Abecasis, Deborah A Nickerson, James G Wilson, Stephen S Rich, Daniel Levy, Ingo Ruczinski, Abraham Aviv, Thomas W Blackwell, Timothy Thornton, Jeff O’Connell, Nancy J Cox, James A Perry, Mary Armanios, Alexis Battle, Nathan Pankratz, Alexander P Reiner, Rasika A Mathias

## Abstract

Telomeres shorten in replicating somatic cells, and telomere length (TL) is associated with age-related diseases ^1,2^. To date, 17 genome-wide association studies (GWAS) have identified 25 loci for leukocyte TL ^3–19^, but were limited to European and Asian ancestry individuals and relied on laboratory assays of TL. In this study from the NHLBI Trans-Omics for Precision Medicine (TOPMed) program, we used whole genome sequencing (WGS) of whole blood for variant genotype calling and the bioinformatic estimation of TL in n=109,122 trans-ethnic (European, African, Asian and Hispanic/Latino) individuals. We identified 59 sentinel variants (p-value <5×10^−9^) from 36 loci (20 novel, 13 replicated in external datasets). There was little evidence of effect heterogeneity across populations, and 10 loci had >1 independent signal. Fine-mapping at *OBFC1* indicated the independent signals colocalized with cell-type specific eQTLs for *OBFC1* (*STN1*). We further identified two novel genes, *DCLRE1B* (*SNM1B*) and *PARN*, using a multi-variant gene-based approach.

## RESULTS

The decreasing costs of high throughput sequencing have enabled WGS data generation at an unprecedented scale, and TOPMed data offer the opportunity to address sample size, population diversity, rare variant evaluation, and fine mapping limitations of prior TL GWAS. We selected TelSeq ^20^ to bioinformatically determine TL due to its computational efficiency and high correlation with Southern blot ^21^ and flowFISH ^22^ measurements (***Materials and Methods. Figures S1a-c***). We developed a novel principal components-based approach to remove technical artifacts arising from the sequencing process that affected TL estimation, which improved accuracy (***Materials and Methods, Figures S1d-e***). Pooled trans-ethnic association analysis was performed with n=109,122 subjects (including 51,654 of European ancestry, 29,260 of African ancestry, 18,019 Hispanic/Latinos, 5,683 of Asian ancestry, and 4,506 of other, mixed, or uncertain ancestries, as determined by HARE ^23^, ***Materials and Methods***); 44% were male and age ranged from <1 to 98 years old (***Table S1***).

Genome-wide tests for association were performed across 163M variants. Using a series of single variant tests for association (primary to identify loci, iterative conditional by chromosome to identify additional independent variants, and joint tests including all independent variants to summarize effect sizes; see ***Materials and Methods***), we identified 59 independently associated variants mapping to 36 loci, each reaching a p-value <5×10^−9^ (***Figure 1, Table 1, Table S2***); 16 known and 20 novel, as further described below.

**Figure 1:**
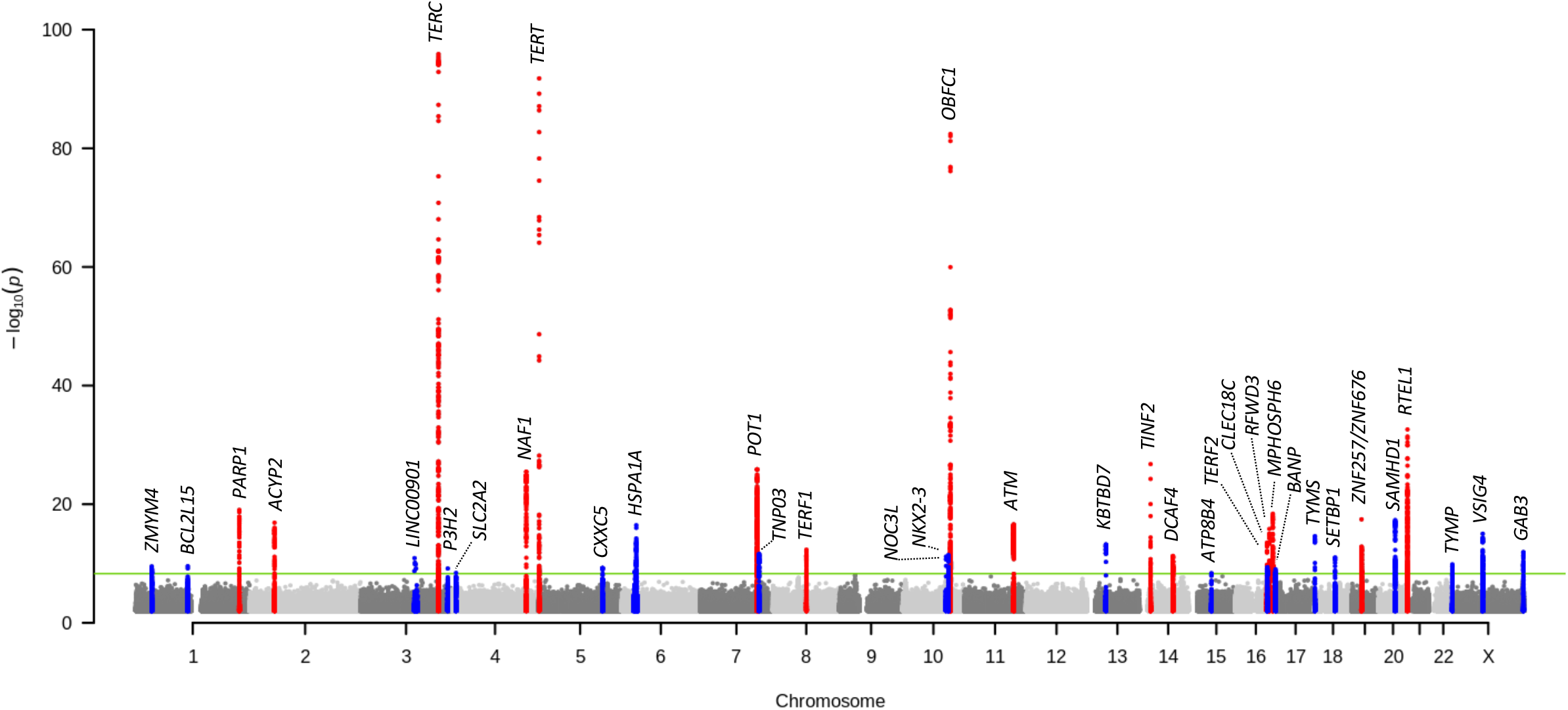
Genome-wide Manhattan plot. Trans-ethnic genome-wide tests for association using 163M sequence identified variants on n=109,122 samples with sequence generated telomere length from TOPMed. All loci had a peak p<5×10^−9^ in the pooled trans-ethnic analysis. Prior known loci are indicated in red, and novel loci are indicated in blue.

**Table 1:**
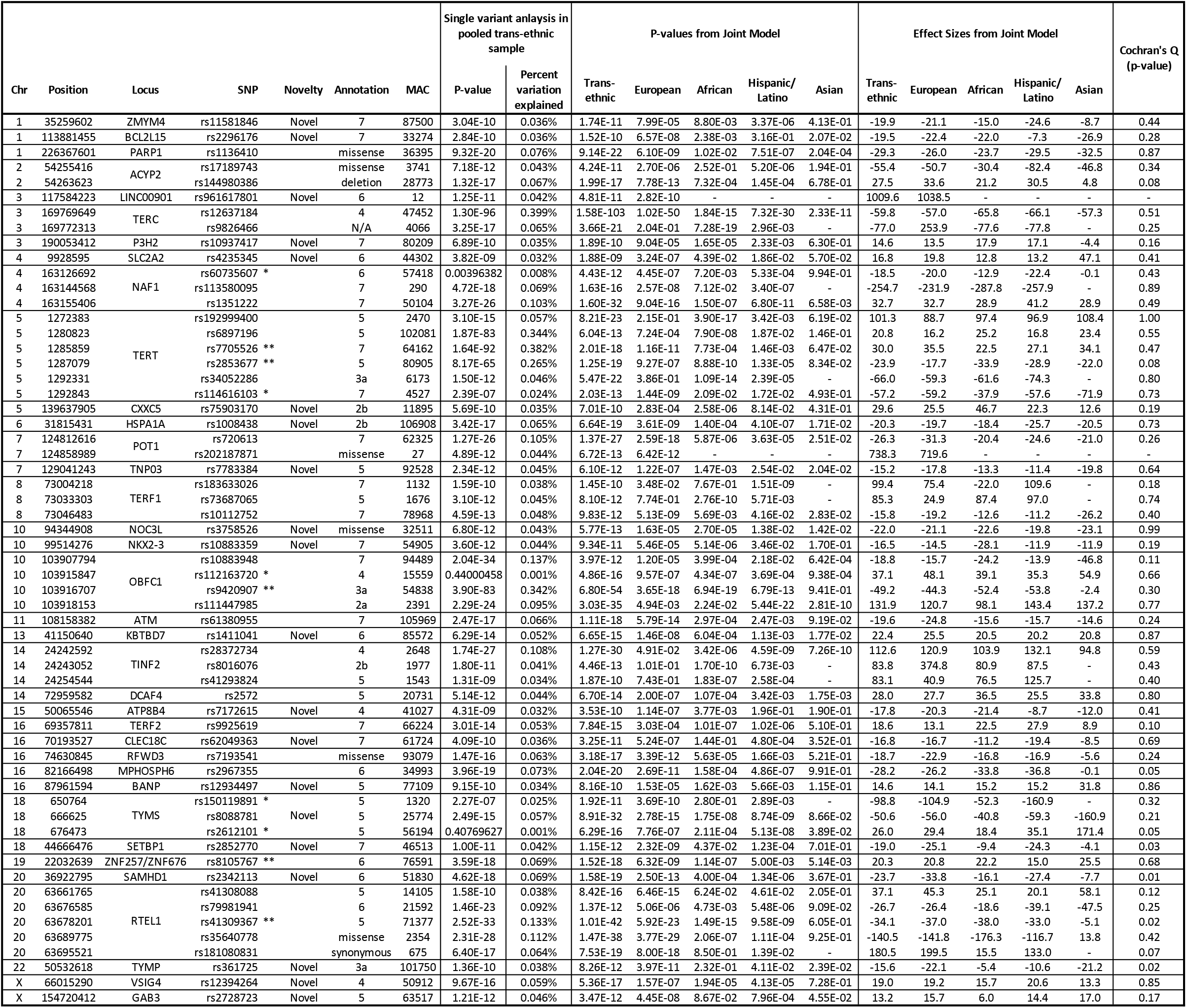
59 independently associated variants mapping to 36 loci from the whole genome sequencing of n=109,122 TOPMed individuals. Loci are labeled as novel if none of the sentinel variants in the locus was in LD (r^2^ < 0.7) with any previously documented GWAS signal for telomere length. There are 5 variants marked with an * where the primary analysis did not meet our threshold of p<5×10^−9^, however they reached significance after conditioning on significant variants mapping to the chromosome (detailed in ***Table S2***). Variants marked with ** are direct matches to prior reported sentinel variants. Percent of trait variation explained by each variant is provided from single-variant association tests. P-values and effect sizes (in base pairs) are reported from a joint model including all variants. P-values for effect heterogeneity across population groups were generated using Cochran’s Q statistic. MAC is the minor allele count from the full combined sample. For all exonic variants, detailed annotation is provided, while for all non-coding variants the RegulomeDB score is given. See also ***Tables S2, S4***.

Of 25 previously known loci, we identified 16 (*PARP1, ACYP2, TERC, NAF1, TERT, POT1, TERF1, OBFC1, ATM, TINF2, DCAF4, TERF2, RFWD3, MPHOSPH6, ZNF208/ZNF257/ZNF676,* and *RTEL1*) with a variant at a p-value <5×10^−9^ ***(Table 1, Table S3***). Directionally consistent and nominal evidence for replication was noted for *CTC1* (rs3027234, p-value = 7.97×10^−5^) and *SENP7* (rs55749605, p-value = 0.023). A signal previously attributed to *PRRC2A* is located less than 200kb from our novel signal for *HSPA1A* but may be distinct given low linkage disequilibrium (r^2^=0.26). We found no evidence of replication (all variants with p-value >0.05) for the remaining previously reported TL loci (*CXCR4, PXK, MOB1B, DKK2/PAPSS1, CARMIL1* and *CSNK2A2*, ***Table S3***). Our comprehensive conditional analyses revealed that there was more than one independent sentinel variant at nine of the sixteen previously reported loci (***Table 1, Figure 2a***). The resolution possible with our trans-ethnic WGS data identified a sentinel variant different from the one previously reported by tagging-based GWAS for 11 of the 16 known loci. At known loci *RTEL1*, *RFWD3*, *POT1*, *ACYP2*, and *PARP1,* our WGS-based sentinels included a coding missense variant in genes *RTEL1*, *RFWD3*, *POT1*, *TSPYL6*, and *PARP1*, respectively. For the remaining known TL loci, many of the non-coding sentinel variants are annotated as having regulatory evidence (RegulomeDB score < 7, ***Table 1***), as illustrated further for *OBFC1* below.

**Figure 2:**
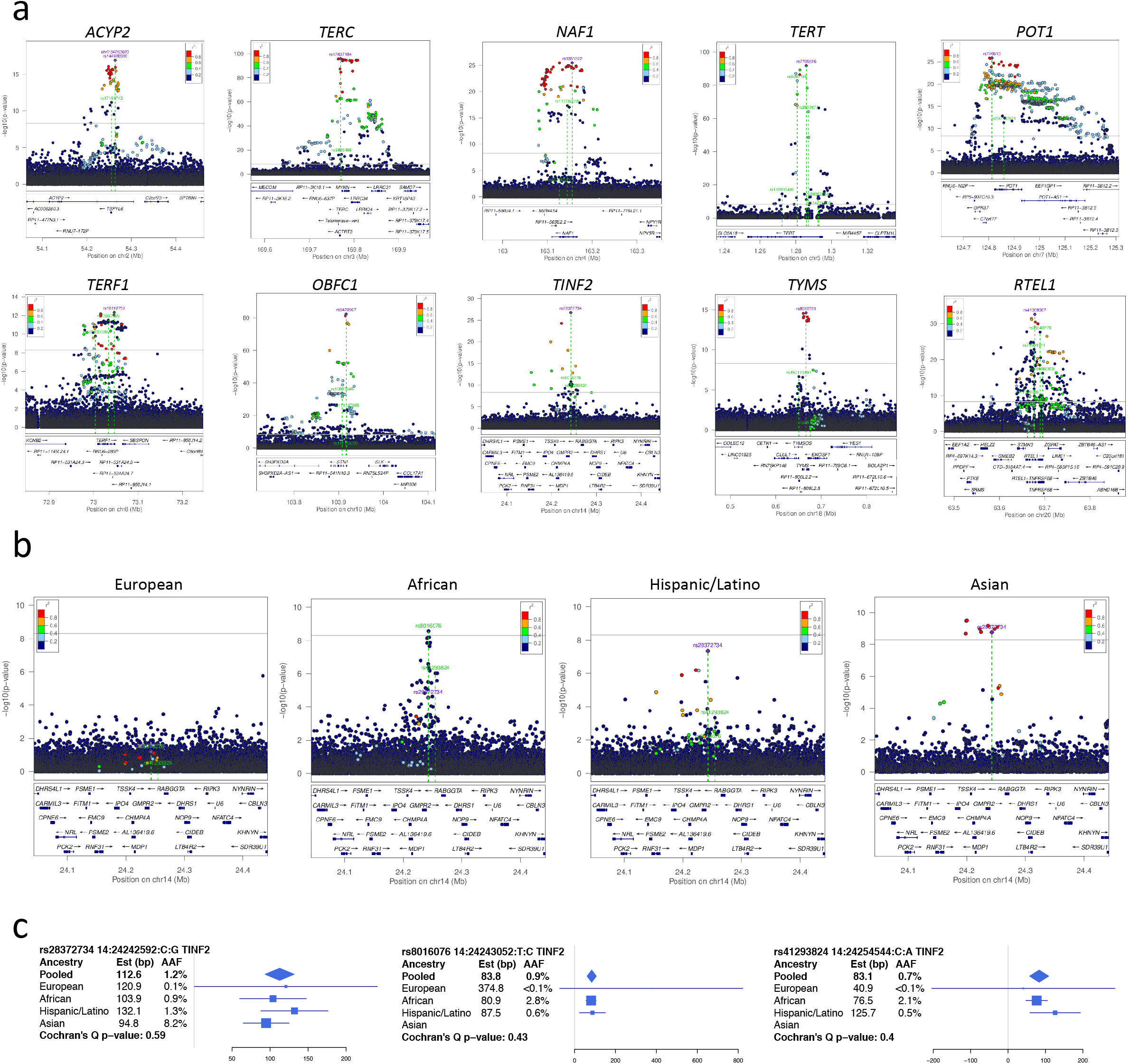
LocusZoom plots for multi-hit loci and *TINF2*. **[a]** LocusZoom plots for all loci with >1 sentinel variant. Linkage disequilibrium (LD) was calculated from the set of samples used in the analysis with respect to the peak variant in the pooled trans-ethnic primary analysis, thereby reflecting LD patterns specific to the TOPMed samples. For each figure, the peak sentinel variant from the pooled trans-ethnic analysis is indexed and labeled in purple, and all independent variants identified through the iterative conditional approach are labeled in green and highlighted with green dotted lines. **[b]** LocusZoom plots for four population groups for the *TINF2* locus. **[c]** Forest plots displaying effect sizes and standard errors, as well as minor allele frequencies, by population group for the three sentinel variants in *TINF2*. See also *Table S2*.

Our 20 novel loci (***Table 1***) had a total of 22 independent sentinel variants, and we tested for replication at the 19 variants available in two prior published GWAS with non-overlapping subjects ^18,19^ (***Figure 3a***). Variants at ten of these loci (*BCL2L15, CXXC5, HSPA1A, NOC3L, NKX2-3, ATP8B4, CLEC18C, TYMS, SAMHD1,* and *TYMP*) had a Bonferroni-corrected p<0.05/19=0.0026, and an additional three had variants with p<0.05 (*TNP03, KBTBD7,* and *BANP*), as did a second variant at *TYMS.* The variant at *SAMHD1* was previously reported at an FDR < 0.05 (p-value = 1.41×10^−7^)^19^ but here has genome-wide significance (p-value = 1.58×10^−19^). While qPCR and TelSeq quantify TL in different units (see ***Materials and Methods)***, there is high consistency in the effects at variants shared between our study and these prior studies (***Figure 3b-c***). Pearson correlations of effect sizes for all 19 shared variants were 0.83 (p-value = 7.0×10^−5^) for our study compared to Dorajoo et al. (n=23,096 Singaporean Chinese) and 0.73 (p-value = 3.7×10^−4^) for our study compared to Li et al. (n=78,592 European). The correlations were stronger (0.93 and 0.84, respectively) when restricted to variants with at least nominal significance in the prior studies (***Figure 3d-e***). The proteins encoded by two of these novel genes have strong biological connections to TL: CXXC5, which physically interacts with ATM and transcriptionally regulates p53 levels ^24^, two proteins implicated in telomere length regulation; and BANP (aka SMAR1) which forms a complex with p53 and functions as a tumor suppressor ^25^.

**Figure 3:**
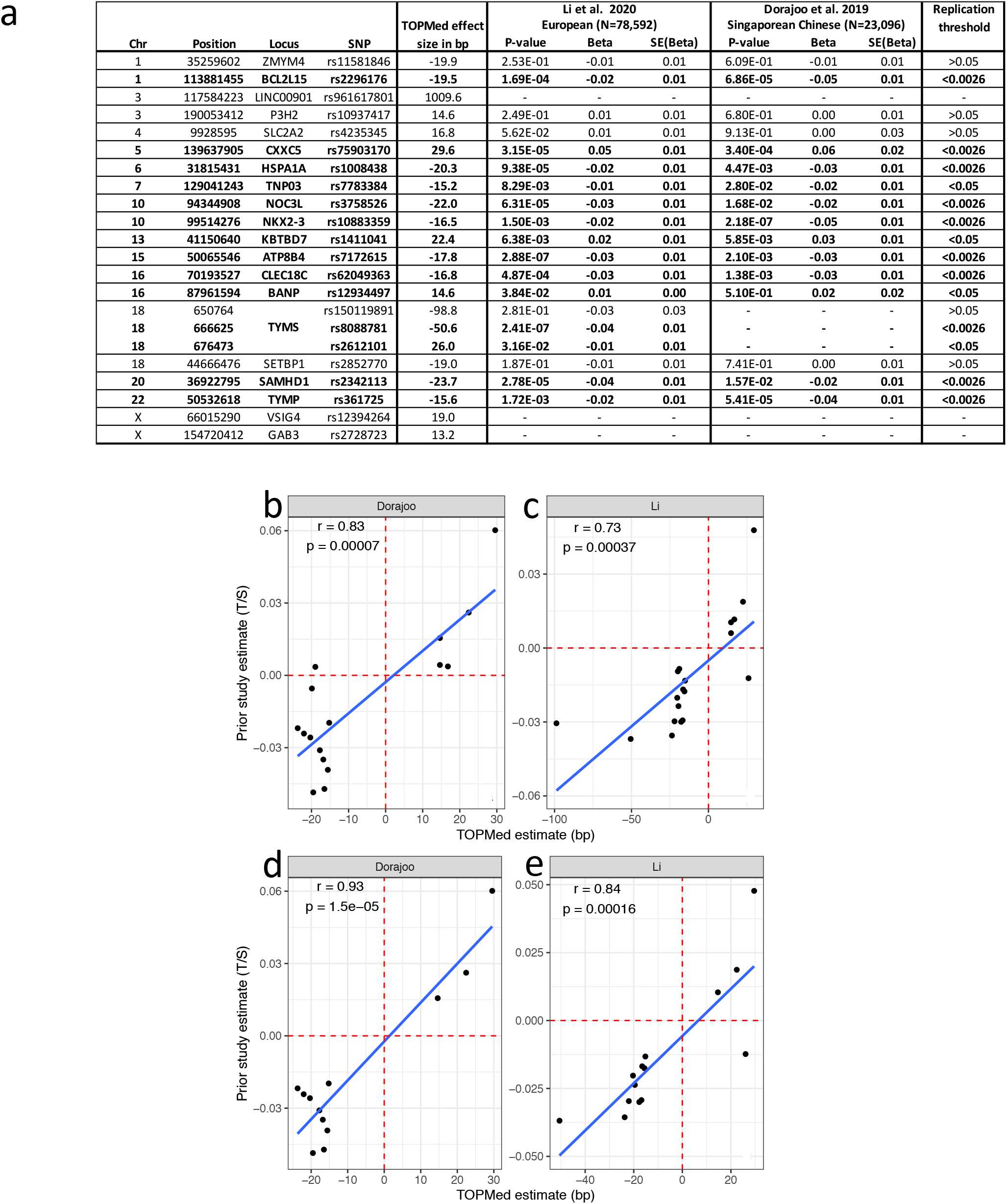
Replication by two prior studies. **[a]** Replication results for the 20 novel TOPMed loci, including 22 variants, pulled from two non-overlapping, prior published GWAS on telomere length using qPCR data. Threshold indicates the stronger level of significance between the two replication studies (Bonferroni significant: p <0.0026; nominally significant: p <0.05; non-significant: p >0.05). **[b-e]** Correlation between the estimated effect sizes for all novel loci (**b-c**) and replicated novel loci (**d-e**, p<0.05 in each dataset) between the present TOPMed pooled trans-ethnic analysis and two prior published GWAS studies.

Each of the 59 sentinel variants individually accounted for a small percentage of phenotypic variation (***Table 1***), consistent with prior GWAS of TL, but cumulatively accounted for 4.35% of TL variance, compared to 2-3% from prior GWAS ^3^. The 37 variants mapping to 16 known loci explained 3.38% of TL variability, with an additional 0.96% explained by the 22 variants mapping to our 20 novel loci; a sizable gain in explained variability for TL in this trans-ethnic sample. Prior GWAS report allelic effects ranging from ~ 49-120 base pairs ^3,4,11,13^. In the TOPMed data, effect sizes for common variants (minor allele frequency, MAF ≥5%) ranged from 2-59 base pairs per allele. Rare and low frequency variants (MAF <5%) showed larger effects (40-1,063 base pairs per allele).

Stratified association analyses were performed in population groups with at least 5,000 samples to evaluate effect heterogeneity of the 59 variants (***Table S4***). Reduced sample sizes, coupled with variation in allele frequency, often limited our power to detect population-specific associations at GWAS thresholds in individual strata (***Table S4***); no additional loci were identified. A major advantage of our analysis was the ability to rely on the individual-level WGS data for the iterative conditional approach to identify the final set of independent sentinel variants at each trans-ethnic-identified locus. Our sentinel variants, identified without relying on tagging through linkage to measured marker variants like prior GWAS, reveal little evidence for heterogeneity across populations (***Table 1***). All Cochran’s Q^26^ p-values (***Table 1***) were above a Bonferroni correction threshold (p-value>0.001), and the five with nominal significance (0.001<p-value<0.05) appear to be primarily driven by differences in the (smallest) Asian stratum. An interesting illustration of a locus with strong allele frequency differences between groups is *TINF2*; the evidence at the peak variant (rs28372734) in the trans-ethnic analysis was driven by the smaller Hispanic/Latino and Asian groups (group-specific p-values 4.6×10^−9^ and 7.3×10^−10^, respectively), and the secondary peak (rs8016076) was driven by the African group (group-specific p-value 1.7×10^−10^, ***Table 1, Figure 2b***). No association is noted in the European group, where these variants are nearly monomorphic (***Figure 2c***).

Gene-based tests in the pooled trans-ethnic sample identified eight protein coding genes with deleterious rare and low frequency (MAF <1%, including singletons) variants associated with TL (p-value <1.8×10^−6^, see ***Materials and Methods, Figure S2***). Six of these genes support a role for rare variants in previously identified GWAS loci (*POT1, TERT, RTEL1, CTC1, SAMHD1,* and *ATM).* The two novel genes have strong biological plausibility: both *DCLRE1B* and *PARN* have been implicated in short telomere syndrome (STS) patients ^27–29^. DCLRE1B protein localizes to the telomere via interaction with the protein of another previously implicated GWAS gene, *TERF2*, and contributes to telomere protection from DNA repair pathways ^30,31^. Notably, two *PARN* loss-of-function variants included in our gene-based test were previously identified in STS patients ^27^. Both rs878853260 and rs876661305 produce frame-shift mutations; rs876661305 produces an early termination codon, truncating most of the nuclease domain ^32^. For each of these eight genes, a leave-one-out approach iterating over each variant included in the aggregate test showed there were no detectible main driver variants and indicated that these gene-based association signals arise from cumulative signal across multiple rare deleterious variants (***Figure S2)***, with the possible exception of *ATM*. When conditioned on the 59 sentinel variants, all genes, except *POT1,* maintained or increased statistical significance (***Figure S2).*** For *POT1*, while the removal of the single variant identified in ***Table 1*** (rs202187871) and conditioning on all 59 sentinels resulted in a decrease in significance from 1.52×10^−24^ to 5.53×10^−18^, it nonetheless remained strongly significant, meeting Bonferroni thresholds.

The identification of multiple independent sentinel variants for several loci offers the unique opportunity to evaluate the potential for distinct regulatory mechanisms (***Figure 2a***, ***Figure S3***). OBFC1 is part of a complex that binds single-stranded telomeric DNA ^33^ and is expressed across multiple tissues in GTEx ^34^ and in whole blood studies meta-analyzed in eQTLGen ^35^. All four signals at the *OBFC1* locus are in the promoter and early introns of *OBFC1* (***Figure 4a-b***). Evidence for eQTL colocalization was detected at the primary, tertiary, and quaternary signals in various tissues (***Materials and Methods***). While all three signals colocalized with *OBFC1* eQTLs, the strongest colocalization evidence in each case was in a distinct tissue: sun exposed skin from the lower leg (posterior probability of shared signal, PPH4 = 98.0%) for the primary, skeletal muscle (PPH4 = 84.4%) for the tertiary, and whole blood (GTEx PPH4 = 75.5%, eQTLGen PPH4 = 75.5%) for the quaternary signal (***Figure 4c-e, Figure S4e, Table S5***). Data from the Roadmap Epigenomics Consortium^36^ indicate that all four signals are consistent with promoter or enhancer regions across blood cells and skeletal muscle tissue (***Figure 4b***). We were unable to perform colocalization analysis on the secondary signal with data from either GTEx or eQTLGen as it is driven by rare variants only in the Hispanic/Latino and Asian individuals (rs111447985, ***Table S4***).

**Figure 4:**
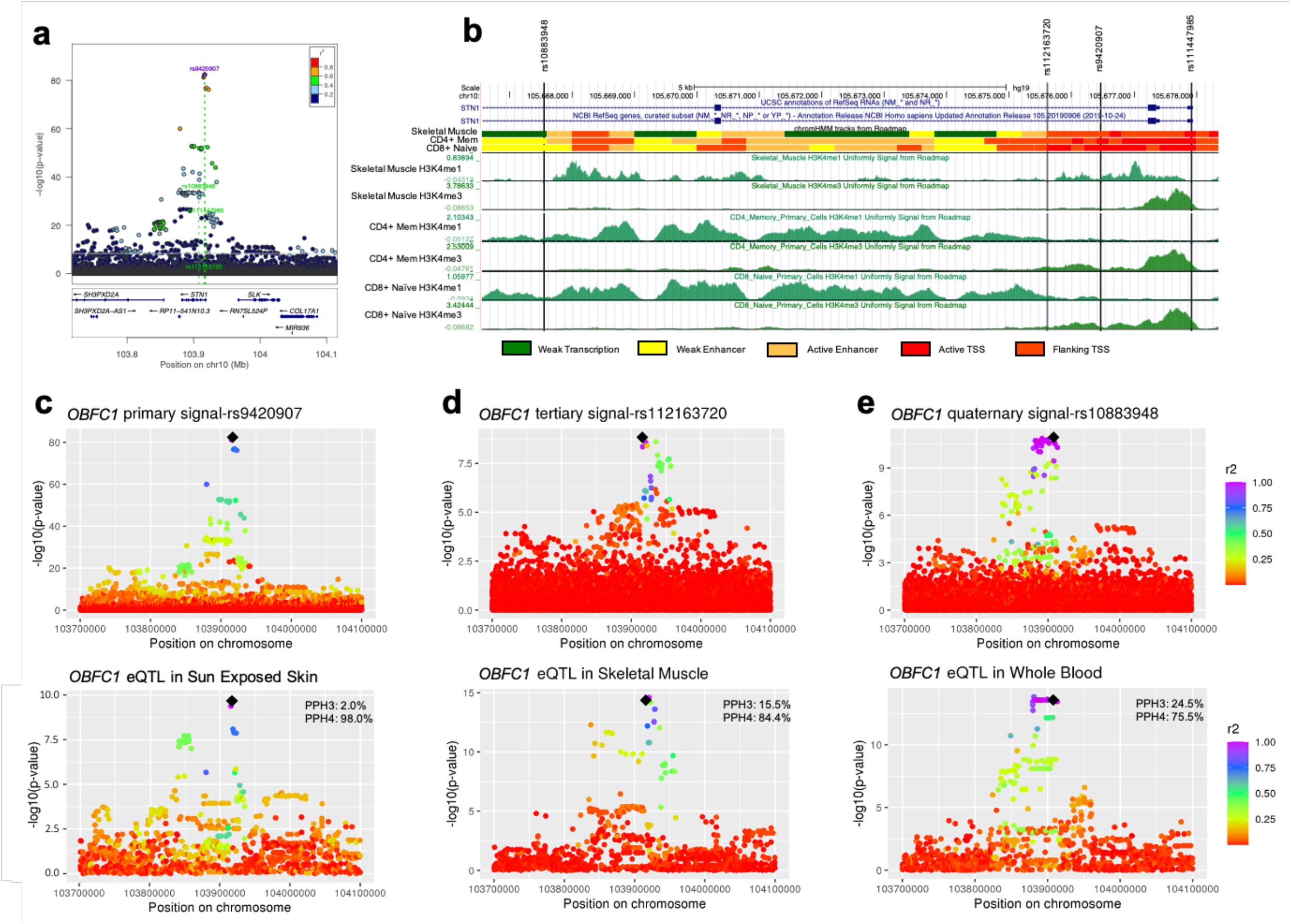
Fine-mapping of multiple *OBFC1* signals. **[a]** LocusZoom plot of the *OBFC1* locus where green dotted lines indicate each independent signal, as in ***Figure 2***. **[b]** Roadmap Epigenomics Consortium data in hg19 coordinates for skeletal muscle tissue, Primary T CD4^+^ memory cells from peripheral blood, and Primary T CD8^+^ naïve cells from peripheral blood (Roadmap samples E108, E037, and E047 respectively; data was not available for sun exposed skin). ChromHMM state model is shown for the 18-state auxiliary model. The state model suggests the primary (rs9420907), secondary (rs111447985), and tertiary (rs112163720) signals are in the promoter region, while the quaternary signal (rs10883948) is in an enhancer region in all Roadmap blood cell types but is transcriptional for peripheral blood monocytes and CD19^+^ B-cells. **[c-e]** GWAS and eQTL results for the primary, tertiary and quaternary signals. Top panels are the GWAS summary statistics from the primary, and iterative conditional analyses which were used to perform colocalization analysis (secondary signal was rare, and not available for colocalization). Bottom panels are eQTLs for *OBFC1* in the indicated tissue from GTEx. The GTEx eQTLs for these tissues do not colocalize with one another (PPH4 < 4.4×10^−7^) and each signal did not significantly colocalize in the other tissues. LD was calculated from the pooled trans-ethnic samples with respect to the sentinel (black diamond). See also ***Figures S3, S4, Table S5***.

Using individual level data within the Vanderbilt University biobank BioVU, we performed a PheWAS (***Table S6)*** using 49 available sentinel variants individually. European and African specific effect sizes from the joint analysis from ***Table 1*** were also combined to create separate polygenic trait scores (PTS) for each population group to conduct PheWAS. PTS values were significantly higher in BioVU African Americans (AAs, mean=-217bp, sd=96bp) compared to European Americans (EAs, mean=-279bp, sd=96bp, p-value<0.05, Welch’s two-sample t-test, ***Figure 5a***), offering evidence that previously observed differences in TL by ancestry (longer TL in individuals of African ancestry^1^) may be explained in part by TL genetics. The largest cumulative effect of the sentinel variants, as evidenced from the PTS, is for the category of neoplasms in the EAs, with higher PTS associated with increased risk to the individual phenotypes (11 of 14 significant results, ***Figure 5b, Table S6)***; associations were only nominal in the BioVU AAs, likely due to lower power from the smaller sample size. Single variant PheWAS (***Table S6)*** in the BioVU EAs are largely replicated within the UK Biobank (UKBB, ***Table S7)***, showing strong associations with neoplasms, and in general, demonstrating the alleles that increased TL also increased risk for these cancer related phenotypes. Additionally, both the UKBB and BioVU data revealed a strong association between the novel *HSPA1A* locus (rs1008438) and type I diabetes related endocrine/metabolism phenotypes; here the allele decreasing TL increased risk for these phenotypes. This agrees with prior associations between shorter TL and increased risk of type 1 diabetes ^37^, and between the protein product of *HSPA1A* (Hsp72) and diabetic ketoacidosis ^38^.

**Figure 5:**
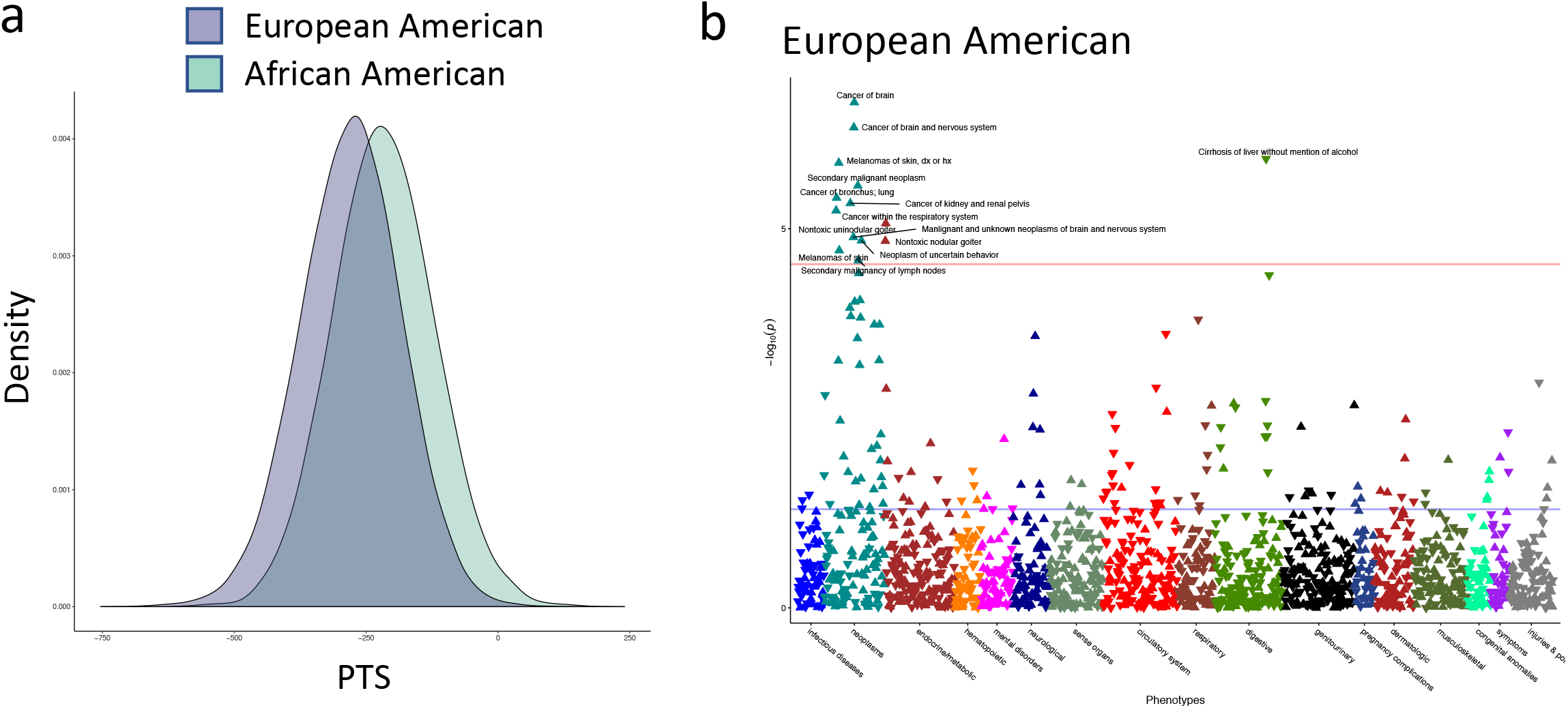
PheWAS results from Vanderbilt BioVU. Polygenic trait score (PTS) analysis across 49 available sentinel variants in BioVU. **[a]** Smoothed distributions of PTS values for European Americans (n=70,439) and African Americans (n=15,174) from BioVU biobank. **[b]** Overview of the PheWAS in the BioVU European Americans. ▲ = higher PTS is associated with the phenotype. ▼ = higher PTS is protective against the phenotype. See also ***Tables S6, S7***.

Leveraging WGS available through the NHLBI TOPMed program, we have illustrated the value of a large, trans-ethnic WGS study to generate a harmonized phenotype of broad interest (i.e. bioinformatically called TL), to confirm known TL GWAS loci, and to identify an additional set of novel loci that map to genes with strong biological plausibility for TL association. The well-powered study enabled identification of rare deleterious variants at known and novel loci with estimated effects larger than those of common variants. Utilizing WGS allowed us the unique opportunity to hone in on causal variants using fine-mapping approaches, and begin to identify tissue-specific genetic effects. We were also able to establish that for most population groups, effects are highly consistent at sentinel variants, despite differences in association strength at loci like *TINF2* and *OBFC1*, where allele frequencies varied among populations. The ability to implement this phenotype assessment of TL in a large, trans-ethnic dataset with pre-existing WGS creates opportunities beyond the genetics of TL. It will expand our ability to evaluate the role of TL and genes determining TL in health and human disease as illustrated by the PheWAS in large biobanks where we document identifiable effects that differ between sentinel variants and the cumulative score across all loci, and start to dissect the genetic basis to TL differences across populations.

## Acknowledgements

Chen Li, Claudia Langernberg, and Veryan Codd for providing summary statistics from their TL GWAS for replication analysis.

Whole genome sequencing (WGS) for the Trans-Omics in Precision Medicine (TOPMed) program was supported by the National Heart, Lung and Blood Institute (NHLBI). Specific funding sources for each study and genomic center are given in the Supplementary Information. Centralized read mapping and genotype calling, along with variant quality metrics and filtering were provided by the TOPMed Informatics Research Center (3R01HL-117626-02S1; contract HHSN268201800002I). Phenotype harmonization, data management, sample-identity QC, and general study coordination, were provided by the TOPMed Data Coordinating Center (3R01HL-120393-02S1; contract HHSN268201800001I). We gratefully acknowledge the studies and participants who provided biological samples and data for TOPMed. The full study specific acknowledgments as well as individual acknowledgements are detailed in the Supplementary Information.

The BioVU projects at Vanderbilt University Medical Center are supported by numerous sources: institutional funding, private agencies, and federal grants. These include the NIH funded Shared Instrumentation Grant S10OD017985 and S10RR025141; CTSA grants UL1TR002243, UL1TR000445, and UL1RR024975 from the National Center for Advancing Translational Sciences. Its contents are solely the responsibility of the authors and do not necessarily represent official views of the National Center for Advancing Translational Sciences or the National Institutes of Health. Genomic data are also supported by investigator-led projects that include U01HG004798, R01NS032830, RC2GM092618, P50GM115305, U01HG006378, U19HL065962, R01HD074711; and additional funding sources listed at https://victr.vumc.org/biovu-funding/.

Support for this work was provided by the National Institutes of Health, National Heart, Lung, and Blood Institute, through the BioData Catalyst program (award 1OT3HL142479-01, 1OT3HL142478-01, 1OT3HL142481-01, 1OT3HL142480-01, 1OT3HL147154). Any opinions expressed in this document are those of the authors and do not necessarily reflect the views of NHLBI, individual BioData Catalyst Consortium members, or affiliated organizations and institutions.

The views expressed in this manuscript are those of the authors and do not necessarily represent the views of the National Heart, Lung, and Blood Institute; the National Institutes of Health; or the U.S. Department of Health and Human Services.

## Author contributions

M.A.T., R.A.M. conceived of and led the study. M.A.T., M.P.C., R.Keener, K.R.I., L.R.Y., C.P.M., D.J., S.G., C.A.L., A.K., M.Arvanitis, J.C.B., E.G.B., J.C.C., Y.C.C., L.M.R., M.H.C., J.E.C., M.D., B.M.P., C.C.L., D.L., I.R., T.W.B., J.A.P., M.Armanios, A.B., A.P.R., R.A.M. drafted the manuscript. M.A.T., M.P.C., R.Keener, J.S.W., J.A.B., D.J., A.K., C.C.L., G.A., D.A.N., J.G.W., S.S.R., D.L., I.R., A.A., T.W.B., T.T., J.O., N.J.C., J.A.P., M.Armanios, A.B., N.P., A.P.R., R.A.M. contributed substantive analytical guidance. M.A.T., M.P.C., R.Keener, K.R.I., J.S.W., L.R.Y., J.Lane, T.W.M., J.A.B., C.P.M., D.J., S.G., C.A.L., A.K., M.Arvanitis, A.V.S., B.H., T.T., N.J.C., M.Armanios, A.B., N.P., A.P.R., R.A.M. performed and led analysis. L.R.Y., L.B., L.C.B., J.C.B., J.B., E.R.B., E.G.B., J.C.C., Y.C.C., B.C., D.D., L.d., D.L.D., B.I.F., M.E.G., M.T.G., S.R.H., B.A.H., M.R.I., T.I., W.C.J., S.Kaab, L.L., J.Lee, S.L., A.M., K.E.N., P.A.P., N.R., L.M.R., C.S., D.E.W., M.M.W., L.W., I.V.Y., W.Z., S.A., P.L.A., D.W.B., B.E.C., Z.C., M.H.C., L.A.C., J.E.C., M.D., R.D., C.E., T.F., X.G., L.H., S.H., J.M.J., E.E.K., A.M.L., C.L., R.L.M., T.N., M.N., M.S.R., E.C.S., J.A.S., N.L.S., J.L.S., M.J.T., H.K.T., R.P.T., M.J.W., Y.Z., K.L.W., S.T.W., R.S.V., K.D.T., M.F.S., E.K.S., M.S., W.H.S., F.S., D.S., J.I.R., D.R., S.R., B.A.R., B.M.P., J.M.P., N.D.P., S.N., C.G.M., B.D.M., D.A.M., S.T.M., F.D.M., A.C.M., R.J.L., R.Kumar, C.K., B.A.K., S.Kelly, S.L.K., R.Kaplan, J.H., H.G., F.D.G., B.G., M.F., P.T.E., M.d., A.C., Y.I.C., E.B., K.C.B., A.E.A., D.K.A., C.A., N.J.C., M.Armanios were involved in the guidance, collection and analysis for one or more of the studies which contributed data to this manuscript. All authors read and approved the final draft.

## Author Information

## Data Deposition Statement

TOPMed genomic data and pre-existing parent study phenotypic data are made available to the scientific community in study-specific accessions in the database of Genotypes and Phenotypes (dbGaP) (https://www.ncbi.nlm.nih.gov/gap/?term=TOPMed). Telomere length calls were derived from the raw sequence data as described in the Online Methods, and the phenotype covariates of age, sex, and race/ethnicity are available through the study-specific dbGAP accession IDs as listed in the Supplementary Information.

## Competing Interests

The authors declare the following competing interests:

J.C.C. has received research materials from GlaxoSmithKline and Merck (inhaled steroids) and Pharmavite (vitamin D and placebo capsules) to provide medications free of cost to participants in NIH-funded studies, unrelated to the current work.

B.I.F. is a consultant for AstraZeneca Pharmaceuticals and RenalytixAI

L.W. is on the advisory board for GlaxoSmithKline and receives grant funding from NIAID, NHLBI, and NIDDK, NIH

I.V.Y. is a consultant for ElevenP15

S.A. receives equity and salary from 23andMe, Inc.

M.H.C. receives grant support from GlaxoSmithKline

S.T.W. receives royalties from UpToDate

E.K.S. received grant support from GlaxoSmithKline and Bayer in the past three years.

B.M.P. serves on the Steering Committee of the Yale Open Data Access Project funded by Johnson & Johnson.

F.D.M. is supported by grants from NIH/NHLBI (HL139054,HL091889,HL132523,HL130045,HL098112,HL056177), the NIH/NIEHS (ES006614), the NIH/NIAID (AI126614), and the NIH/ Office of Director (OD023282). Vifor Pharmaceuticals provided medicine and additional funding to support recruitment for HL130045. Dr. Martinez is a council member for the Council for the Developing Child.

P.T.E. is supported by a grant from Bayer AG to the Broad Institute focused on the genetics and therapeutics of cardiovascular diseases. Dr. Ellinor has also served on advisory boards or consulted for Bayer AG, Quest Diagnostics, and Novartis.

K.C.B. receives royalties from UpToDate

G.A. is an employee of Regeneron Pharmaceuticals and owns stock and stock options for Regeneron Pharmaceuticals.

## Correspondence and requests for materials

should be addressed to *Rasika A Mathias, ScD, 410-550-2487*

## ALL TABLE LEGENDS

**Table S1, Related to TOPMed study populations, Materials and Methods:** Sample demographics summarized by each set of analysis performed: pooled trans-ethnic and four non-overlapping population groups (defined with HARE, using reported race/ethnicity and genetically inferred ancestry in combination; “Other/Uncertain” includes individuals with maximum stratum probability from HARE < 0.7, as well as Brazilians and Samoans who were excluded from the HARE analysis).

**Table S2, Related to Table 1 and Figure 2:** Results of the iterative conditional analysis using individual level data. Results are presented by chromosome providing a detailed overview of the conditional step at which each variant was identified as the peak signal. (Note that some variants had p-value>5×10^−9^ in the primary analysis, but p-value<5×10^−9^ in a conditional step). Within each chromosome, variants are ordered not on position, but in the sequence of identification through the conditional analysis showing the iterative process used: Primary and Rounds 1-6 of conditioning. Chromosomes varied in the number of analyses needed until no additional variants were included (maximum steps = 7 on chromosome 5).

**Table S3, Related to Assessing novelty of identified loci and variants, Materials and Methods:** TOPMed results from the primary pooled trans-ethnic analysis for all prior telomere length GWAS sentinel variants with reported p-value <5×10^−8^ in prior published studies.

**Table S4, Related to Table 1:** Association results for the single variant association analysis for each of the 59 sentinel variants from ***Table 1***. Single variant results are shown for the pooled trans-ethnic analysis, and each population group. Variants with a minor allele count <5 were not included in the analysis and are listed as “-”.

**Table S5, Related to Figure 4:** Iterative conditional analysis was repeated on chromosome 10 focusing exclusively on the *OBFC1* locus, defined as a 2Mb window around the original top sentinel, rs9420907. The sentinel for each signal was consistent with the iterative conditional analysis performed on the entirety of chromosome 10 (***Table S2***). The summary statistics from each iterative conditional analysis were used to perform colocalization analysis on the non-primary signals. Colocalization analysis was performed using coloc with all significant gene-tissue pairs in GTEx and with all genes in a 2Mb window around rs9420907 in eQTLGen. All results with PPH4 > 0.7 for each signal are reported.

**Table S6, Related to Figure 5:** Results of the PheWAS performed on 49 available sentinel variants and a polygenic trait score across these 49 sentinel variants using BioVU self-identified African Americans (AA, n=15,174) and BioVU self-identified European Americans (EA, n=70,439). The cumulative TL risk score for the BioVU AA and EA samples was derived from the African and European specific effects sizes from ***Table 1***, respectively. Results were evaluated at a Bonferroni threshold corrected for the number of informative phecodes for each variant.

**Table S7, Related to Figure 5:** Results of the PheWAS conducted using SAIGE, a method employing generalized linear mixed models and allowing for imbalance between case and control counts, adjusting for genetic relatedness, sex, birth year and the first 4 principal components of ancestry, on approximately 400,000 UK Biobank (UKBB) participants. Results were downloaded from files provided by the University of Michigan PheWeb server (http://pheweb.sph.umich.edu/SAIGE-UKB/about) on 47 available sentinel variants. Results were evaluated at a Bonferroni threshold corrected for the number of phecodes available for each variant (N=1,403).

## ALL FIGURE LEGENDS

**Figure S1, Related to Estimating telomere length for whole-genome sequencing (WGS) samples and Batch adjustment to correct for technical confounders, Materials and Methods:** For 2,389 samples from the Jackson Heart Study (JHS), **[a]** scatter plot with Pearson correlation between TelSeq and Computel length estimates; [**b]** Comparison of computational times for TelSeq and Computel; **[c]** scatter plots with Pearson correlations between TelSeq (left) and Computel (right) and Southern blot TL estimates; **[d]** scatter plot with Pearson correlation between TelSeq and Southern blot TL estimates after adjustment for the final set of 200 batch principal components (bPCs) used in our full analysis. Colors indicate on which plate samples were shipped to the sequencing center, in Panels A, C and D; and **[e]** scatter plot with Pearson correlation between bPC-adjusted TelSeq and flowFISH data on 19 samples from the GeneSTAR study.

**Figure S2, Related to Gene-based coding variant tests - Tests for association, Materials and Methods**: Eight genes were identified as passing the Bonferroni threshold based on number of genes tested (p-value < 0.05/27,558 = 1.8×10^−6^). For each gene, a leave-one-out analysis was performed iterating the SMMAT test and leaving one variant out at a time. The plots show the change in SMMAT p-value for each variant (orange line with marker) relative to the variant’s allele frequency (blue bar), the overall gene-based test including all variants (dotted red line) and the single variant results for all variants with an MAC≥5 that were included in single variant tests for association (brown and green diamonds). For each gene, the number of rare and deleterious variants included in SMMAT is indicated. For any variant with a MAC≥5, a single variant test was also performed as part of the primary analysis. The count of these variants is indicated. In addition the SMMAT p-value for these genes when conditioning on the 59 sentinel variants is also given.

**Figure S3, Related to Figure 4: [a]** LocusZoom plots for four population groups for the *OBFC1* locus. Linkage disequilibrium (LD) was calculated from the set of samples used in the analysis with respect to the peak variant in the pooled trans-ethnic primary analysis, thereby reflecting LD patterns specific to the TOPMed samples. For each figure, the peak sentinel variant from the pooled trans-ethnic analysis is indexed and labeled in purple, and all independent variants identified through the iterative conditional approach are labeled in green and highlighted with green dotted lines. **[b]** Forest plots displaying effect sizes and standard errors, as well as minor allele frequencies, by population group for the four sentinel variants in *OBFC1*.

**Figure S4, Related to Figure 4: [a-d]** Credible set analysis and colocalization analysis in eQTLGen. Manhattan plots for each *OBFC1* signal are shown where p-values were taken from the appropriate conditional analysis output and LD was calculated with respect to the sentinel variant. Credible set variants are indicated with black diamonds; the sentinel variant is indicated as a black diamond with red outline. **[e]** Manhattan plot for colocalization analysis of the *OBFC1* quaternary signal with an *OBFC1* eQTL in eQTLGen. PPH3 and PPH4 for colocalization are indicated in the top right corner of the eQTL plot. LD was calculated with respect to the sentinel, indicated with a black diamond on each graph, and with pooled trans-ethnic analysis samples.

## MATERIALS AND METHODS

### TOPMed study populations

To perform this trans-ethnic genome-wide association study of telomere length, we leveraged the whole genome sequence samples available through the NHLBI Trans Omics for Precision Medicine (TOPMed) program. The program currently consists of more than 80 participating studies ^1^,with a range of study designs as described in Taliun et al ^2^ (*Nature, submitted, 2019*). Participants are mainly U.S. residents with diverse ancestry, race, and ethnicity (European, African, Hispanic/Latino, Asian, and Other). Smaller representation comes from non-US populations including Samoan, Brazilian, and Asian studies. Details on the specific samples included for telomere length analysis are outlined below, summarized in ***Table S1***, and described by TOPMed ^1^.

### TOPMed whole genome sequencing (WGS)

WGS was performed to an average depth of 38X using DNA isolated from blood, PCR-free library construction, and Illumina HiSeq X technology. Details for variant calling and quality control are described in Taliun et al. ^2^ (*Nature, submitted, 2019*). Briefly, variant discovery and genotype calling was performed jointly, across all the available TOPMed Freeze 8 studies, using the GotCloud ^3^ pipeline resulting in a single multi-study genotype call set.

### Estimating telomere length for whole-genome sequencing (WGS) samples

A variety of computational tools exist that leverage WGS data to generate an estimate of telomere length ^4^. Here, we performed a thorough comparison of two leading methods for estimating telomere length from WGS data to choose the preferred scalable method for performing the estimation on all available samples from TOPMed. The first method, TelSeq ^5^, calculates an estimate of individual telomere length using counts of sequencing reads containing a fixed number of repeats of the telomeric nucleotide motif TTAGGG. Given that 98% of our data was sequenced using read lengths of 151 or 152 (as confirmed from the SEQ field in the analyzed CRAM files), we chose to use a repeat number of 12. These read counts are then normalized according to the number of reads in the individual WGS data set with between 48% and 52% GC content to adjust for potential technical artifacts related to GC content. The second method, Computel ^6^ uses an alignment-based method to realign all sequenced reads from an individual to a “telomeric reference sequence”. Reads aligning to this reference sequence are considered to be telomeric and are included in the estimate of telomere length. Because Computel performs a complete realignment, additional computational steps are involved compared to those needed for TelSeq.

To compare the results and scalability from these two methods, we first directly compared estimates obtained from TelSeq and Computel on 2,389 samples from the Jackson Heart Study (JHS) and found them to be highly correlated with one another (Pearson correlation r=0.98, ***Figure S1a***). We also compared computational time to generate the telomere length estimates on these samples and show that Computel is around ten times more time-consuming (***Figure S1b***). This is in part due to the fact that Computel requires CRAM-formatted files (as the WGS data are currently stored) to first be converted back to Fastq format (while TelSeq requires a CRAM to BAM conversion), but also due to the computationally expensive step of realignment to the telomeric reference genome that the Computel algorithm employs.

Telseq generates an estimate of TL in bp similar to laboratory assays such as Southern blot ^7^ and flowFISH ^8^; in contrast qPCR approaches are represented as T/S ratios^9,10^. As a further comparison to orthogonally measured telomere length values, we used data on the same 2,389 samples from JHS with Southern blot^7^ telomere length estimates ^11^. For these samples, the Southern blot assay was performed on the same source DNA sample that was used to generate the WGS in TOPMed. The Pearson correlation values between the TelSeq and Computel estimates and the Southern blot estimates did not differ (r=0.58 and 0.56 for TelSeq and Computel, respectively, ***Figure S1c***). Based on our observation that both Computel and TelSeq showed similar correlation to the Southern blot estimates and high correlation with each other, and that TelSeq was an order of magnitude more computationally efficient, we chose to use TelSeq to perform telomere length estimation on our data. Final telomere length estimation was performed on a set of 128,901 samples whose CRAM-files were available for analysis at the TOPMed IRC at the time of analysis.

### Batch adjustment to correct for technical confounders

To account for technical sources of variability in our telomere length estimates, both within a study (see, for example, colors in ***Figures S1a*** and ***S1b*** which indicate grouping by shared 96-well plate for shipment to the sequencing center) and across studies, we developed a method to estimate components of technical variability in our samples. We estimated these covariates using the sequencing data itself, similar to methods developed for other multivariate genomics data types (SVA or PEER factors ^12,13^), using aligned sequencing reads and relying on the fact that genomic coverage patterns of aligned reads can reflect technical variation.

We computed average sequencing depth for every 1,000 bp genomic region (“bin”) genome-wide using mosdepth ^14^. We removed bins known to be problematic: those containing repetitive DNA sequence with difficulty mapping (mappability<1.0 using 50bp k-mers in GEMTools v1.759 ^15^) or that overlap the list of known problematic SVs ^16^ or overlap known CNVs in the Database of Genomic Variants. To avoid overcorrecting for sex, bins were limited to autosomes. After normalizing the approximately 150,000 remaining bin counts within sample, we performed Randomized Singular Value Decomposition ^17^ (rSVD), a scalable alternative to principal components analysis, to generate batch principal components (bPCs). We included increasing numbers of bPCs in a linear regression model predicting TelSeq TL, and computed the correlation of the resulting residuals with external data measurements, including Southern blot measurements for JHS (n=2,389) and the Women’s Health Initiative (WHI; n=596) and age at blood draw (JHS n=3,294; WHI n=10,708). Based on the observed correlation, the final decision was to include the first 200 bPCs across all samples. Using the n=2,389 JHS samples described above, we compared TL estimates before and after batch correction. The percent of variance in TL explained by sequencing plate reduced from 21.9% (baseline) to 10.5% (200 bPCs), and the variance explained by age increased from 8.0% (baseline) to 10.3% (200 bPCs), evidence that the signal-to-noise ratio was improved. Overall, the correlation between the bPC corrected TL and Southern blot data improved from r=0.58 to 0.68 (***Figure S1d***) in the JHS data and from r=0.54 to 0.72 for the WHI data. Further, we compared TelSeq estimates of 19 samples within a single sequencing batch from the GeneSTAR study to the clinical gold standard of flowFISH ^8^ (***Figure S1e***) and observed a correlation of 0.80 in both granulocytes and lymphocytes. Therefore, our data show that we are able to reduce the sequencing artifacts stemming from batch variability to attain correlation of TelSeq to Southern blot similar to the correlation of TelSeq to flowFISH.

### Samples included in genetic analysis

All samples with telomere length estimated from the WGS data from TOPMed Freeze 8 were considered for inclusion, provided they had consent that allowed for genetic analysis of telomere length. Only samples with sequencing read lengths of 151 or 152 base pairs and having age at blood draw data available were included. For the set of samples that were part of a duplicate pair/group (either part of the intended duplicates designed by TOPMed, or a duplicate identified across the studies through sample QC) only one sample from each duplicated pair/group was retained. The final counts and demographic summary statistics for subjects grouped by TOPMed study for all 54 studies included in our analysis are shown in ***Table S1.***

While reported race/ethnicity data are available in TOPMed, these data have limitations for analysis that include individuals with missing information or non-specific responses (e.g., ‘other’ or ‘multiple’) and high variability in genetically inferred measures of ancestry among individuals with the same reported race/ethnicity. To overcome these limitations, we used a computational method called HARE (harmonized ancestry and race/ethnicity), a newly developed machine learning approach for jointly leveraging reported and genetic data in the definition of population strata for GWAS ^18^. HARE uses provided race/ethnicity labels and genetic ancestry principal component (PC) values to compute probability estimates for each individual’s membership in each race/ethnicity stratum. For our HARE analysis, we used provided race (Asian, Black, White) or Hispanic ethnicity group (Central American, Costa Rican, Cuban, Dominican, Mexican, Puerto Rican, South American) as input labels to define population strata, and we used 11 PCs computed with PC-AiR using 638,486 LD-pruned (r^2^ < 0.1) autosomal variants with minor allele frequency > 1% to represent genetic ancestry. Genetic outliers for population strata were identified as individuals for whom their maximum stratum probability was more than 5 times greater than their reported stratum probability. Stratum values for genetic outliers and individuals with missing or non-specific race/ethnicity were imputed as the stratum for which they had the highest membership probability.

Our primary analysis allowed for heterogeneous residual variance (see **Primary single variant tests for association** for details) among groups defined jointly by study and HARE-based population stratum assignment, with minor study-specific modifications to account for small strata. We required at least 30 individuals within a study-HARE grouping and collapsed individuals into merged HARE groups within a study as necessary to retain everyone for analysis. For our population-specific analyses, we used HARE assignment to stratify individuals into the following population groups: African (corresponding to the Black HARE stratum), Asian (Asian), European (White), and Hispanic/Latino (Central American, Costa Rican, Cuban, Dominican, Mexican, Puerto Rican, and South American). To better preserve genetic ancestry similarity among individuals in population-specific stratified analyses, we restricted to individuals for whom their HARE population stratum membership probability was at least 0.7; the population stratum counts in ***Table S1*** reflect the counts in the stratified analyses, where individuals not meeting this criterion are labeled as “Other/Uncertain”.

Samoan individuals from the Samoan Adiposity Study and Brazilian individuals from the Reds-III Brazil study were excluded from the HARE analyses due to their unique ancestry in the TOPMed dataset; these studies were treated as their own population groups for analyses.

### Primary single variant tests for association

Genome-wide tests for association were performed using the R Bioconductor package GENESIS ^19^. The primary analysis included all available trans-ethnic TOPMed samples (n=109,122). A secondary analysis was performed for all population groups with n>5,000, which included European (n=51,654), African (n=29,260), Hispanic/Latino (n=18,019) and Asian (n=5,683) groups as defined above using HARE. Prior to genetic modeling, we generated residuals from a linear regression model on all 109,122 samples with 200 batch principal components (bPCs), as described above; for clarity we call these residuals *TL_bPC_* below. For the pooled trans-ethnic analysis, we used a fully-adjusted two-stage model, as described in the next two bullets^20^. For each population-specific analysis, the same approach was used, limited to samples within that population group.

▪ **Stage 1:** We fit a linear mixed model (LMM) on n=109,122 samples, using *TL_bPC_* as the outcome; adjusting for age, sex, study, sequencing center, and 11 PC-AiR ^21^ PCs of ancestry as fixed effect covariates; including a random effect with covariance matrix proportional to a sparse empirical kinship matrix computed with PC-Relate^22^ to account for genetic relatedness among samples; and allowing for heteroskedasticity of residual variance across study-HARE groups as defined above. The marginal residuals from this Stage 1 model were then inverse-normalized and rescaled by their original standard deviation. This rescaling restores values to the original trait scale, providing more meaningful effect size estimates from subsequent association tests ^23^.
▪ **Stage 2:** We fit a second LMM on all n=109,122 samples, using the inverse-normalized and rescaled residuals from Stage 1 as the outcome; all other aspects of the model including fixed effects adjustment, random effects, and residual variance structure were identical to the model in Stage 1. This two-stage covariate adjustment has been shown to be most effective at controlling for false-positives and increasing statistical power in this setting^20^. The output of this Stage 2 model was then used to perform both single variant and gene-based tests for association.

### Single variant tests for association

We used the output of the two-stage LMM to perform score tests of association for each variant with minor allele count (MAC) ≥ 5 that passed TOPMed Informatics Research Center (IRC) at the University of Michigan quality filters ^2^ and which had <10% of samples with read depth <10. Genotype effect size estimates and percent of variability explained (PVE) were approximated from the score test results ^24^.

### Assessing significance, performing conditional analysis to identify independent variants, and defining genetic loci

A p-value cutoff of 5×10^−9^ was used to determine genome-wide significance in the primary trans-ethnic analysis. We identified our set of independent significant variants (as reported in ***Table 1***) through an iterative conditioning process within each chromosome. For a given chromosome, if at least one variant from the primary analysis crossed the genome-wide significance cutoff, this peak variant was included as an additional fixed-effect covariate in a new two-stage LMM (see Stages 1 and 2 described above), and score test results were examined to see if any remaining variants crossed the 5×10^−9^ threshold. If so, we performed a second round of conditioning, including both the original peak variant and the new conditional peak variant as fixed-effect covariates in another two-stage LMM; and so on, adding conditional peak variants for additional rounds (***Table S2***). For each chromosome, the conditioning procedure was completed when no additional variants crossed the genome-wide threshold (p < 5×10^−9^) on that chromosome. At each step, all variants passing the p < 5×10^−9^ threshold were examined in BRAVO ^25^ to assess quality, and 334 variants were filtered out due to variant call quality issues. In the case where a current peak variant was flagged for quality, the next most significant variant, provided its p-value was below the 5×10^−9^ cutoff, was considered the peak variant instead. Variants were grouped into loci based on physical distance and an examination of linkage disequilibrium (LD) patterns, and locus names were determined using a combination of previous literature, known telomere biology, and physical location.

### Cumulative Percent of Variability Explained (PVE)

Through the iterative conditional approach, we identified a total of 59 variants (***Table 1***) that met our genome-wide significance threshold of p < 5×10^−9^. The cumulative PVE values for this full set of 59 variants (4.35%), the set of 37 variants mapping to known loci (3.38%), and the set of 22 variants mapping to novel loci (0.96%, see **Assessing novelty of identified loci and variants** below for definition of novel variants) were each estimated jointly using approximations from multi-parameter score tests. This joint PVE approximation is similar to the single variant PVE approximation described above, except that the set of variants is tested jointly, accounting for covariance among the genotypes. This approach avoids inadvertently double counting any partially shared signal among the set of identified variants.

### Joint tests for association and testing for heterogeneity across population groups

We then performed joint association analyses for the full multi-ethnic sample (n=109,122), as well as each of the four population groups with n>5000, to determine effect sizes and p-values when all 59 variants were considered together. Using the inverse-normalized and rescaled residuals from the primary analysis Stage 1 LMM as the outcome, we fit a new Stage 2 LMM that was the same as described above, except with the additional inclusion of the genotypes for these 59 variants as additive genetic fixed effects. Given this joint modeling framework, the variant effect size estimates are all adjusted for one another. These estimates were used as input for calculation of a polygenic trait score used for the PheWAS described below. Finally, we tested for heterogeneity of effect sizes from these analyses among the population groups by adapting Cochran’s Q statistic and its p-value ^26^, commonly used to test for effect heterogeneity in meta-analysis (***Table 1***). For each variant, the effect size estimates and standard errors from each population group analysis were used to calculate Q, and a Bonferroni threshold of 0.001 (0.05/59) was used to assess significance.

### Assessing novelty of identified loci and variants

For each of the 59 variants identified, we examined the linkage disequilibrium (LD) with previously reported sentinel variants from 17 published GWAS. Only sentinel variants with p < 5×10^−8^ in their published study were considered, which included a total of 56 variants (***Table S3***). If one of our variants had LD≥0.7 with a published variant, it was labeled as a known variant/part of a known locus; otherwise it was labeled as novel in ***Table 1***. Within a locus, we then compared each independent variant to the prior GWAS reported sentinel variant. If they were identical, the variant was labeled as a known sentinel variant in ***Table 1***. Additionally, locus names for the final set of independent variants were selected based on (i) prior GWAS study definition for known loci, and (ii) the specific gene annotation for each variant mapping directly to a gene for novel loci.

### Replication of novel results with published GWAS

To determine whether our novel results are supported by findings from prior studies, we considered the two largest most recent studies of telomere genetics in European ^27^ (Li et al., n=78,592) and Asian ^28^ (Dorajoo et al., n=26,875) ancestry individuals. These studies both used telomere length as measured by qPCR. For all novel variants in ***Table 1***, we pulled the effect size estimates, standard errors, and p-values, where available (***Figure 3a***). These results were available in at least one of the two studies for 19 of our 22 novel variants, so we considered a p-value cutoff of 0.05/19 = 0.0026 to be replicated, after multiple testing correction. We also labeled variants where at least one study reported p<0.05 as suggestive.

### Gene-based coding variant tests - Variant annotation

For its use in gene-based tests for association, annotation based variant filtering and GENCODE v28 gene model-based ^29^ aggregation was performed using the TOPMed freeze 8 WGSA Google BigQuery-based variant annotation database on the BioData Catalyst powered by Seven Bridges platform (http://doi.org/10.5281/zenodo.3822858). The annotation database was built using variant annotations for TOPMed freeze 8 variants gathered by Whole Genome Sequence Annotator (WGSA) version v0.8^30^ and formatted by WGSAParsr version 6.3.8 (https://github.com/UW-GAC/wgsaparsr). Variants were annotated as exonic, splicing, transcript ablation/amplification, ncRNA, UTR5, UTR3, intronic, upstream, downstream, or intergenic using Ensembl Variant effect predictor (VEP) ^31^. Exonic variants were further annotated as frameshift insertion, frameshift deletion, frameshift block substitution, stop-gain, stop-loss, start-loss, non-frameshift insertion, non-frameshift deletion, non-frameshift block substitution, nonsynonymous variant, synonymous variant, or unknown. Additional scores used included REVEL ^32^, MCAP ^33^ or CADD ^34^ effect prediction algorithms.

### Gene-based coding variant tests - Tests for association

Gene-based association testing was performed on the pooled trans-ethnic dataset (n=109,122). To improve the power to identify rare variant associations in coding regions, we aggregated deleterious rare coding variants in all protein-coding genes and then tested for association with telomere length. To enrich for likely functional variants, only variants with a “deleterious” consequence for its corresponding gene or genes ^35^, were included. For each protein-coding gene, a set of rare coding variants (MAF < 0.01, including singletons where MAC=1, restricted to variants which passed IRC quality filters ^2^ and which had <10% of samples with read depth <10) was constructed, which was composed of all stop-gain, stop-loss, start-loss, transcript ablation, transcript amplification, splice acceptor variants, splice donor variants and frameshift variants, as well as the exonic missense variants that fulfilled one of these criteria: 1) REVEL score > 0.5, 2) predicted M_CAP value was “Damaging”, or 3) CADD PHRED-scaled score > 30. We applied the variant Set Mixed Model Association Test (SMMAT) ^36^ as implemented in GENESIS, using the genesis_tests app on the Analysis Commons ^37^, with MAF based variant weights given by a beta-distribution with parameters of 1 and 25, as proposed by Wu et al. ^38^, and using the same two-stage LMM output as used in the primary single variant analysis. Only genes with a cumulative MAC ≥ 5 over all variants were evaluated, leaving a total of 27,558 genes, and significance was evaluated after a Bonferroni correction for multiple testing (p < 0.05 / 27,558 = 1.815×10^−6^) (***Figure S2***).

Next, we sought to determine the influence of each rare deleterious variant in each significant gene on the association signal. We iterated through the variants, removing one variant at a time (leave-one-out approach) ^39^, and repeated the SMMAT analysis. If a variant made a large contribution to the original association signal, one would expect the signal to be significantly weakened with the removal of the variant from the set ***(Figure S2***).

Finally, we further tested for independence of the gene-based and single variant signals by performing a conditional SMMAT analysis that included the 59 genome-wide significant variants from our primary analysis as fixed-effect covariates in the two-stage LMM. These 59 variants were also removed from the aggregated set of rare variants for a gene if they had been previously included (e.g. rs202187871 in *POT1*). All other analysis parameters were the same as described above (***Figure S2***).

### Colocalization analysis of *OBFC1* signals using GTEx ^40^ and eQTLGen^41^

Iterative conditional analysis was repeated for chromosome 10 focusing on a 2Mb window centered on the primary signal near *OBFC1* (rs10883948). The original pooled GWAS results (n = 109,122) were used for colocalization analysis with the primary signal while the appropriate round of conditional analysis was used for each subsequent signal (e.g., the output of the second round of conditional analysis was used for colocalization analysis with the tertiary signal). Credible set analysis was performed using CAVIAR on primary signal data and the output of each conditional analysis each with a single assumed causal variant ^42^. For each independent *OBFC1* signal, the credible set contained the top sentinel variant (***Figure S4a-d***).

Colocalization analysis was performed using coloc, a Bayesian posterior probability method that estimates the probability of shared signal across testing modalities at each variant ^43^. We report the posterior probability that the two signals are independent (PPH3) and the posterior probability that the two signals overlap (PPH4). The sentinel variants from each signal were assayed as expression quantitative trait loci (eQTLs) in both GTEx^**40**^ and eQTLGen ^41^ datasets. For each sentinel, significant gene-tissue pairs for that sentinel were identified from GTEx v8 (FDR < 0.05) and assayed for colocalization comparing the beta and standard error of the beta from our GWAS results and the eQTL results. For colocalization analysis in the eQTLGen dataset, all eGenes within a 2Mb window of the sentinel were identified and assayed for colocalization comparing the MAF, p-value, and number of observations. MAF was estimated for eQTLGen data using the TOPMed MAF. Colocalization analysis was not possible for the *OBFC1* secondary signal as that variant is absent in both datasets and a representative proxy variant was not available. Roadmap ^44^ data was accessed July, 2020 using the hg19 (February, 2009 release) UCSC genome browser^45^ track data hubs

### Phenome-wide association tests (PheWAS)

Using individual level data within the Vanderbilt University biobank BioVU, PheWAS ^48^ (tests for association between genotype and phenotype) were performed using the 49 (of 59) sentinel variants available in the multi-ethnic genotyping array (MEGA) chip results imputed to the Haplotype Reference Consortium ^49^. Single variant tests using SNP dosage values were performed for all available phecodes (number of cases at least 20), including the covariates age, sex, genotype batch and the first ten ancestry principal components. Analysis was performed separately in BioVU self-identified African Americans (AA, n=15,174) and BioVU self-identified European Americans (EA, n=70,439). In addition, European and African specific effect sizes from the joint analysis from ***Table 1*** were combined to create separate polygenic trait scores (PTS) for each population group which were then tested for association with available phecodes, again including the covariates age, sex, genotype batch and the first ten ancestry principal components. Results were evaluated at a Bonferroni threshold corrected for the number of informative phecodes for each variant (range n=1,114-1,361) or the PTS (n=1,704) (***Table S6***). Analysis was performed using the PheWAS R package ^50^.

We queried United Kingdom Biobank (UKBB) GWAS results using the University of Michigan PheWeb web interface (http://pheweb.sph.umich.edu/SAIGE-UKB/). The UKBB PheWeb interface contains results from a SAIGE ^51^ genetic analysis of 1,403 ICD-based traits of 408,961 UKBB participants of European ancestry. PheWeb is a publicly accessible database that allows querying genome-wide association results for 28 million imputed genetic variants. 47 out of our 59 sentinel variants were present in PheWeb. We report all hits passing a Bonferroni correction for the number of tests performed for each variant (0.05/1403 = 3.6×10^−5^, ***Table S7***).

## Notes

### Summary of Updates

Analysis was redone on a substantially expanded data set using a streamlined analysis approach

