## Supplementary Information for "Novel genetic determinants of telomere length from a trans-ethnic analysis of 109,122 whole genome sequences in TOPMed"

**Table S1, Related to TOPMed study populations, Materials and Methods:** Sample demographics summarized by each set of analysis performed: pooled trans-ethnic and four non-overlapping population groups (defined with HARE, using reported race/ethnicity and genetically inferred ancestry in combination; "Other/Uncertain" includes individuals with maximum stratum probability from HARE < 0.7, as well as Brazilians and Samoans who were excluded from the HARE analysis).

| | | | | HARE Population Group | | | | | Sequencing Center | | | | | |
|---|---|---|---|---|---|---|---|---|---|---|---|---|---|---|
| Study | Total Count | Male Count (Pct) | Mean Age (SD, Range) | European | African | Hispanic/Latino | Asian | Other/ Uncertain | Baylor | Broad | Psomagen | NYGC | UW | WASHU |
| AFLMU | 350 | 266 (0.76) | 48 (11.9, 19-79) | 349 | - | - | - | 1 | - | 350 | - | - | - | - |
| Amish | 1109 | 557 (0.5) | 52 (16.8, 20-91) | 1109 | - | - | - | - | - | 1109 | - | - | - | - |
| ARIC | 3829 | 1833 (0.48) | 58 (5.9, 44-79) | 3578 | 245 | 5 | - | 1 | 3825 | 4 | - | - | - | - |
| BioMe | 9455 | 3804 (0.4) | 56 (14.9, 19-90) | 3180 | 2497 | 3212 | 428 | 138 | 6828 | - | - | - | - | 2627 |
| CAMP | 1662 | 860 (0.52) | 31 (15, 9-66) | 1270 | 223 | 131 | 21 | 17 | - | - | - | - | 1662 | - |
| CARDIA | 3396 | 1480 (0.44) | 45 (7, 27-61) | 1802 | 1578 | 7 | 3 | 6 | 3396 | - | - | - | - | - |
| CARE_BADGER | 35 | 23 (0.66) | 11 (3.5, 6-18) | 9 | 2 | 24 | - | - | - | - | - | - | 35 | - |
| CARE_CLIC | 12 | 7 (0.58) | 11 (3.9, 6-17) | 2 | - | 10 | - | - | - | - | - | - | 12 | - |
| CARE_PACT | 22 | 11 (0.5) | 9 (2.4, 6-14) | 6 | 2 | 12 | - | 2 | - | - | - | - | 22 | - |
| CARE_TREXA | 65 | 33 (0.51) | 11 (3.3, 6-18) | 16 | 1 | 46 | - | 2 | - | - | - | - | 65 | - |
| CFS | 1281 | 576 (0.45) | 39 (19.6, 2-92) | 612 | 664 | 4 | 1 | - | - | - | - | - | 1281 | - |
| ChildrensHS_GAP | 7 | 5 (0.71) | 19 (0.6, 18-20) | 2 | - | 5 | - | - | - | - | - | - | 7 | - |
| ChildrensHS_IGERA | 152 | 52 (0.34) | 26 (2, 23-31) | 50 | 3 | 94 | - | 5 | - | - | - | - | 152 | - |
| ChildrensHS_MetaAir | 52 | 26 (0.5) | 19 (0.7, 18-21) | 11 | - | 38 | - | 3 | - | - | - | - | 52 | - |
| CHIRAH | 263 | 96 (0.37) | 19 (10.9, 8-40) | - | 263 | - | - | - | - | - | - | - | 263 | - |
| CHS | 3510 | 1466 (0.42) | 74 (5.7, 64-98) | 2775 | 712 | 14 | - | 9 | 3510 | - | - | - | - | - |
| COPDGene | 10054 | 5327 (0.53) | 59 (9.1, 39-85) | 6785 | 3235 | 22 | 1 | 11 | - | 8241 | - | - | 1813 | - |
| CRA | 2982 | 1591 (0.53) | 29 (15.2, 0-93) | - | - | 2982 | - | - | - | - | - | - | 2982 | - |
| DHS | 372 | 159 (0.43) | 60 (8.7, 36-86) | - | 372 | - | - | - | - | 372 | - | - | - | - |
| ECLIPSE | 2308 | 1483 (0.64) | 62 (7.9, 40-76) | 2224 | 41 | 11 | 9 | 23 | - | - | - | - | - | 2308 |
| EOCOPD | 74 | 21 (0.28) | 49 (5.8, 26-59) | 73 | - | - | - | 1 | - | - | - | - | 74 | - |
| FHS | 4118 | 1887 (0.46) | 61 (15.6, 20-98) | 4102 | - | 1 | 1 | 14 | - | 4118 | - | - | - | - |
| GALAI | 893 | 417 (0.47) | 19 (9.6, 8-55) | 3 | - | 888 | - | 2 | - | - | - | - | 893 | - |
| GALAII | 3456 | 1706 (0.49) | 13 (3.7, 4-60) | 11 | 12 | 3410 | - | 23 | - | - | - | 986 | 2470 | - |
| GeneSTAR | 1485 | 606 (0.41) | 44 (12.6, 20-79) | 833 | 651 | 1 | - | - | - | 102 | 1383 | - | - | - |
| GENOA | 1188 | 362 (0.3) | 62 (10, 25-95) | - | 1188 | - | - | - | - | 101 | - | - | 1087 | - |
| GenSalt | 1823 | 961 (0.53) | 39 (9.6, 15-62) | - | - | - | 1823 | - | 1823 | - | - | - | - | - |
| GOLDN | 947 | 445 (0.47) | 48 (16.4, 18-87) | 946 | - | 1 | - | - | - | - | - | - | 947 | - |
| HCHS_SOL | 3854 | 1564 (0.41) | 47 (14, 18-75) | 40 | 59 | 3638 | - | 117 | 3854 | - | - | - | - | - |
| HVH | 691 | 452 (0.65) | 64 (12, 22-92) | 653 | 26 | 5 | 7 | - | 616 | 75 | - | - | - | - |
| HyperGEN | 1863 | 684 (0.37) | 47 (12.8, 18-85) | 1 | 1861 | - | 1 | - | - | - | - | - | 1863 | - |
| IPF | 309 | 185 (0.6) | 65 (10.2, 25-91) | 298 | 2 | 7 | - | 2 | - | - | - | - | - | 309 |
| JHS | 3324 | 1237 (0.37) | 56 (12.9, 21-92) | - | 3321 | 1 | 1 | 1 | - | - | - | - | 3324 | - |
| LTRC | 1340 | 695 (0.52) | 63 (10.7, 21-88) | 1226 | 83 | 25 | 4 | 2 | - | 1340 | - | - | - | - |
| Mayo_VTE | 1279 | 603 (0.47) | 57 (16.8, 18-96) | 1261 | 6 | 9 | 1 | 2 | 1279 | - | - | - | - | - |
| MESA | 5310 | 2519 (0.47) | 61 (9.7, 39-91) | 1861 | 1872 | 947 | 605 | 25 | - | 5310 | - | - | - | - |
| MLOF | 5094 | 4566 (0.9) | 26 (19.2, 0-92) | 3590 | 514 | 698 | 215 | 77 | 2924 | - | - | 2170 | - | - |
| OMG_SCD | 616 | 278 (0.45) | 33 (12.3, 18-84) | - | 614 | 1 | - | 1 | 616 | - | - | - | - | - |
| PCGC_CHD | 1905 | 1020 (0.54) | 28 (17.5, 0-72) | 1371 | 101 | 272 | 114 | 47 | - | 1905 | - | - | - | - |
| PharmHU | 173 | 87 (0.5) | 8 (5.7, 0-21) | - | 154 | 16 | 2 | 1 | 173 | - | - | - | - | - |
| PIMA | 50 | 17 (0.34) | 10 (3.2, 6-17) | 1 | 2 | 47 | - | - | - | - | - | - | 50 | - |
| PUSH_SCD | 364 | 184 (0.51) | 12 (5.2, 3-20) | 2 | 356 | 5 | - | 1 | 364 | - | - | - | - | - |
| REDS-III_Brazil | 2610 | 1227 (0.47) | 20 (13.9, 0-77) | - | - | - | - | 2610 | 2610 | - | - | - | - | - |
| SAFS | 996 | 425 (0.43) | 37 (15.1, 16-84) | 19 | 3 | 971 | - | 3 | - | 996 | - | - | - | - |
| SAGE | 1838 | 839 (0.46) | 17 (6.7, 7-41) | 1 | 1831 | 2 | 1 | 3 | - | - | - | 443 | 1395 | - |
| SAPPHIRE_asthma | 4688 | 1649 (0.35) | 34 (14, 12-64) | 220 | 4284 | 112 | 52 | 20 | - | - | - | - | 4688 | - |
| SARP | 1748 | 670 (0.38) | 36 (17.4, 6-84) | 1059 | 594 | 44 | 42 | 9 | - | - | - | 1748 | - | - |
| Samoan | 1284 | 510 (0.4) | 45 (11.3, 25-65) | - | - | - | - | 1284 | - | - | - | 906 | 378 | - |
| THRV | 2144 | 1056 (0.49) | 52 (10, 17-86) | - | - | - | 2144 | - | 2144 | - | - | - | - | - |
| VAFAR | 117 | 76 (0.65) | 59 (8.5, 26-75) | 117 | - | - | - | - | - | 117 | - | - | - | - |
| VU_AF | 1106 | 807 (0.73) | 53 (11.2, 16-81) | 1044 | 48 | 8 | 4 | 2 | - | 1106 | - | - | - | - |
| walk_PHaSST | 415 | 193 (0.47) | 37 (13.5, 12-84) | 1 | 397 | 16 | - | 1 | 415 | - | - | - | - | - |
| WGHS | 115 | 0 (0) | 49 (3.4, 45-59) | 114 | - | - | - | 1 | - | 115 | - | - | - | - |
| WHI | 10989 | 1 (0) | 69 (6.9, 50-87) | 9027 | 1443 | 277 | 203 | 39 | - | 10989 | - | - | - | - |
| **Total** | **109122** | **47604 (0.44)** | **49 (20.4, 0-98)** | **51654** | **29260** | **18019** | **5683** | **4506** | **34377** | **36350** | **1383** | **6253** | **25515** | **5244** |

**Table S2, Related to Table 1 and Figure 2:** Results of the iterative conditional analysis using individual level data. Results are presented by chromosome providing a detailed overview of the conditional step at which each variant was identified as the peak signal. (Note that some variants had p-value>5x10$^{-9}$ in the primary analysis, but p-value<5x10$^{-9}$ in a conditional step). Within each chromosome, variants are ordered not on position, but in the sequence of identification through the conditional analysis showing the iterative process used: Primary and Rounds 1-6 of conditioning. Chromosomes varied in the number of analyses needed until no additional variants were included (maximum steps = 7 on chromosome 5).

| | | | | | LD with Primary | | Primary | | Conditional Round 1 | | Conditional Round 2 | | Conditional Round 3 | | Conditional Round 4 | | Conditional Round 5 | | Conditional Round 6 | |
|---|---|---|---|---|---|---|---|---|---|---|---|---|---|---|---|---|---|---|---|---|
| chr | pos | Locus | rsNum | Round | r2 | D' | Est | p-value | Est | p-value | Est | p-value | Est | p-value | Est | p-value | Est | p-value | Est | p-value |
| 1 | 226367601 | PARP1 | rs1136410 | Primary | | | **-28.50** | **9.32E-20** | | | | | | | | | | | | |
| 1 | 35259602 | ZMYM4 | rs11581846 | Cond_1 | 0.001 | 0.06 | -19.09 | 3.04E-10 | **-19.14** | **2.74E-10** | | | | | | | | | | |
| 1 | 113881455 | BCL2L15 | rs2296176 | Cond_2 | 0.000 | 0.02 | -19.70 | 2.84E-10 | -19.58 | 3.58E-10 | **-19.56** | **3.69E-10** | | | | | | | | |
| 2 | 54263623 | ACYP2 | rs144980386 | Primary | | | **28.23** | **1.32E-17** | | | | | | | | | | | | |
| 2 | 54255416 | ACYP2 | rs17189743 | Cond_1 | 0.003 | 1.00 | -58.93 | 7.18E-12 | **-55.56** | **1.06E-10** | | | | | | | | | | |
| 3 | 169769649 | TERC | rs12637184 | Primary | | | **-59.07** | **1.30E-96** | | | | | | | | | | | | |
| 3 | 169772313 | TERC | rs9826466 | Cond_1 | 0.005 | 1.00 | -70.30 | 3.25E-17 | **-75.18** | **1.70E-19** | | | | | | | | | | |
| 3 | 117584223 | LINC00901 | rs961617801 | Cond_2 | 0.000 | 1.00 | 1063.26 | 1.25E-11 | 1050.17 | 2.02E-11 | **1049.68** | **2.05E-11** | | | | | | | | |
| 3 | 190053412 | P3H2 | rs10937417 | Cond_3 | 0.008 | 0.22 | 14.49 | 6.89E-10 | 14.13 | 1.66E-09 | 14.23 | 1.26E-09 | **14.19** | **1.39E-09** | | | | | | |
| 4 | 163155406 | NAF1 | rs1351222 | Primary | | | **28.15** | **3.27E-26** | | | | | | | | | | | | |
| 4 | 163144568 | NAF1 | rs113580095 | Cond_1 | 0.004 | 1.00 | -273.46 | 4.72E-18 | **-251.28** | **1.96E-15** | | | | | | | | | | |
| 4 | 163126692 | NAF1 | rs60735607 | Cond_2 | 0.106 | 1.00 | -7.44 | 0.00396382 | -18.51 | 1.28E-11 | **-18.43** | **1.55E-11** | | | | | | | | |
| 4 | 9928595 | SLC2A2 | rs4235345 | Cond_3 | 0.000 | 0.01 | 16.86 | 3.82E-09 | 16.79 | 4.30E-09 | 16.77 | 4.47E-09 | **16.92** | **3.22E-09** | | | | | | |
| 5 | 1285859 | TERT | rs7705526 | Primary | | | **50.87** | **1.64E-92** | | | | | | | | | | | | |
| 5 | 1292331 | TERT | rs34052286 | Cond_1 | 0.005 | 0.27 | -48.46 | 1.50E-12 | **-70.34** | **2.75E-24** | | | | | | | | | | |
| 5 | 1287079 | TERT | rs2853677 | Cond_2 | 0.257 | 0.60 | -39.84 | 8.17E-65 | -21.47 | 1.31E-15 | **-22.91** | **1.54E-17** | | | | | | | | |
| 5 | 1272383 | TERT | rs192999400 | Cond_3 | 0.004 | 0.97 | 82.11 | 3.10E-15 | 91.36 | 1.72E-18 | 87.95 | 2.94E-17 | **89.49** | **8.00E-18** | | | | | | |
| 5 | 1292843 | TERT | rs114616103 | Cond_4 | 0.000 | 0.06 | -40.67 | 2.39E-07 | -42.83 | 5.06E-08 | -44.51 | 1.47E-08 | -47.07 | 2.14E-09 | **-57.82** | **3.25E-13** | | | | |
| 5 | 1280823 | TERT | rs6897196 | Cond_5 | 0.316 | 0.82 | 44.48 | 1.87E-83 | 25.00 | 1.15E-17 | 21.30 | 4.98E-13 | 20.77 | 1.83E-12 | 20.81 | 1.61E-12 | **21.49** | **3.07E-13** | | |
| 5 | 139637905 | CXXC5 | rs75903170 | Cond_6 | 0.001 | 0.09 | 30.40 | 5.69E-10 | 30.19 | 6.88E-10 | 29.87 | 1.03E-09 | 29.64 | 1.36E-09 | 29.60 | 1.41E-09 | 29.48 | 1.63E-09 | **29.42** | **1.76E-09** |
| 6 | 31815431 | HSPA1A | rs1008438 | Primary | | | **-19.75** | **3.42E-17** | | | | | | | | | | | | |
| 7 | 124812616 | POT1 | rs720613 | Primary | | | **-26.40** | **1.27E-26** | | | | | | | | | | | | |
| 7 | 129041243 | TNP03 | rs7783384 | Cond_1 | 0.000 | 0.01 | -15.89 | 2.34E-12 | **-15.89** | **2.23E-12** | | | | | | | | | | |
| 7 | 124858989 | POT1 | rs202187871 | Cond_2 | 0.000 | 0.96 | 727.22 | 4.89E-12 | 726.39 | 4.98E-12 | **727.04** | **4.70E-12** | | | | | | | | |
| 8 | 73046483 | TERF1 | rs10112752 | Primary | | | **-17.10** | **4.59E-13** | | | | | | | | | | | | |
| 8 | 73033303 | TERF1 | rs73687065 | Cond_1 | 0.004 | 1.00 | 88.85 | 3.10E-12 | **83.89** | **4.90E-11** | | | | | | | | | | |
| 8 | 73004218 | TERF1 | rs183633026 | Cond_2 | 0.003 | 0.99 | 101.90 | 1.59E-10 | 97.09 | 1.13E-09 | **97.77** | **8.66E-10** | | | | | | | | |
| 10 | 103916707 | OBFC1 | rs9420907 | Primary | | | **-54.72** | **3.90E-83** | | | | | | | | | | | | |
| 10 | 103918153 | OBFC1 | rs111447985 | Cond_1 | 0.004 | 1.00 | 109.76 | 2.29E-24 | **116.56** | **2.72E-27** | | | | | | | | | | |
| 10 | 94344908 | NOC3L | rs3758526 | Cond_2 | 0.001 | 0.05 | -21.48 | 6.80E-12 | -22.01 | 1.88E-12 | **-21.79** | **3.06E-12** | | | | | | | | |
| 10 | 99514276 | NKX2-3 | rs10883359 | Cond_3 | 0.021 | 0.43 | -18.14 | 3.60E-12 | -16.71 | 1.46E-10 | -17.12 | 4.98E-11 | **-17.29** | **3.23E-11** | | | | | | |
| 10 | 103915847 | OBFC1 | rs112163720 | Cond_4 | 0.026 | 1.00 | 3.35 | 0.44000458 | 24.06 | 6.29E-08 | 26.92 | 1.47E-09 | 27.21 | 9.69E-10 | **26.89** | **1.50E-09** | | | | |
| 10 | 103907794 | OBFC1 | rs10883948 | Cond_5 | 0.246 | 0.98 | -27.96 | 2.04E-34 | -8.30 | 0.0015196 | -12.20 | 3.81E-06 | -12.10 | 4.55E-06 | -12.29 | 3.20E-06 | **-18.90** | **8.80E-12** | | |
| 11 | 108158382 | ATM | rs61380955 | Primary | | | -19.29 | **2.47E-17** | | | | | | | | | | | | |
| 13 | 41150640 | KBTBD7 | rs1411041 | Primary | | | 22.08 | **6.29E-14** | | | | | | | | | | | | |
| 14 | 24242592 | TINF2 | rs28372734 | Primary | | | 108.81 | **1.74E-27** | | | | | | | | | | | | |
| 14 | 72959582 | DCAF4 | rs2572 | Cond_1 | 0.004 | 0.18 | 26.37 | 5.14E-12 | **26.29** | **5.67E-12** | | | | | | | | | | |
| 14 | 24243052 | TINF2 | rs8016076 | Cond_2 | 0.000 | 1.00 | 79.49 | 1.80E-11 | 80.64 | 9.06E-12 | **80.60** | **9.23E-12** | | | | | | | | |
| 14 | 24254544 | TINF2 | rs41293824 | Cond_3 | 0.000 | 1.00 | 80.93 | 1.31E-09 | 82.24 | 6.96E-10 | 82.48 | 6.19E-10 | **84.54** | **2.26E-10** | | | | | | |
| 15 | 50065546 | ATP8B4 | rs7172615 | Primary | | | -17.00 | **4.31E-09** | | | | | | | | | | | | |
| 16 | 82166498 | MPHOSPH6 | rs2967355 | Primary | | | -27.86 | **3.96E-19** | | | | | | | | | | | | |
| 16 | 74630845 | RFWD3 | rs7193541 | Cond_1 | 0.001 | 0.07 | -18.69 | 1.47E-16 | **-18.71** | **1.34E-16** | | | | | | | | | | |
| 16 | 69357811 | TERF2 | rs9925619 | Cond_2 | 0.002 | 0.14 | 18.64 | 3.01E-14 | 18.61 | 3.24E-14 | **18.92** | **1.21E-14** | | | | | | | | |
| 16 | 70193527 | CLEC18C | rs62049363 | Cond_3 | 0.010 | 0.14 | -16.23 | 4.09E-10 | -16.38 | 2.73E-10 | -16.62 | 1.48E-10 | **-16.65** | **1.36E-10** | | | | | | |
| 16 | 87961594 | BANP | rs12934497 | Cond_4 | 0.004 | 0.11 | 14.93 | 9.15E-10 | 14.86 | 1.06E-09 | 14.75 | 1.42E-09 | 14.88 | 9.96E-10 | **14.83** | **1.12E-09** | | | | |
| 18 | 666625 | TYMS | rs8088781 | Primary | | | **-27.73** | **2.49E-15** | | | | | | | | | | | | |
| 18 | 676473 | TYMS | rs2612101 | Cond_1 | 0.357 | 0.96 | 2.15 | 0.40769627 | **22.60** | **3.37E-12** | | | | | | | | | | |
| 18 | 44666476 | SETBP1 | rs2852770 | Cond_2 | 0.000 | 0.01 | -18.59 | 1.00E-11 | -18.56 | 1.06E-11 | **-18.58** | **9.99E-12** | | | | | | | | |
| 18 | 650764 | TYMS | rs150119891 | Cond_3 | 0.000 | 0.60 | -76.76 | 2.27E-07 | -80.32 | 6.13E-08 | -100.93 | 1.98E-11 | **-100.93** | **1.96E-11** | | | | | | |
| 19 | 22032639 | ZNF257_ZNF676 | rs8105767 | Primary | | | 20.57 | 3.59E-18 | | | | | | | | | | | | |
| 20 | 63678201 | RTEL1 | rs41309367 | Primary | | | **-29.14** | **2.52E-33** | | | | | | | | | | | | |
| 20 | 63689775 | RTEL1 | rs35640778 | Cond_1 | 0.022 | 0.99 | -120.85 | 2.31E-28 | **-144.36** | **6.39E-39** | | | | | | | | | | |
| 20 | 36922795 | SAMHD1 | rs2342113 | Cond_2 | 0.003 | 0.06 | -23.27 | 4.62E-18 | -23.26 | 4.60E-18 | **-23.25** | **4.52E-18** | | | | | | | | |
| 20 | 63695521 | RTEL1 | rs181080831 | Cond_3 | 0.000 | 0.47 | 174.01 | 6.40E-17 | 177.46 | 1.49E-17 | 176.23 | 2.36E-17 | **177.20** | **1.55E-17** | | | | | | |
| 20 | 63661765 | RTEL1 | rs41308088 | Cond_4 | 0.033 | 0.99 | 29.50 | 1.58E-10 | 40.34 | 6.80E-18 | 39.84 | 1.64E-17 | 39.69 | 2.09E-17 | **40.36** | **5.88E-18** | | | | |
| 20 | 63676585 | RTEL1 | rs79981941 | Cond_5 | 0.044 | 0.91 | -37.56 | 1.46E-23 | -29.82 | 6.54E-15 | -29.91 | 5.22E-15 | -29.97 | 4.45E-15 | -29.46 | 1.26E-14 | **-25.64** | **2.82E-11** | | |
| 22 | 50532618 | TYMP | rs361725 | Primary | | | **-14.99** | **1.36E-10** | | | | | | | | | | | | |
| X | 66015290 | VSIG4 | rs12394264 | Primary | | | **18.62** | **9.67E-16** | | | | | | | | | | | | |
| X | 154720412 | GAB3 | rs2728723 | Cond_1 | 0.009 | 0.11 | 13.80 | 1.21E-12 | **13.76** | **1.38E-12** | | | | | | | | | | |

**Table S3, Related to Assessing novelty of identified loci and variants, Materials and Methods:** TOPMed results from the primary pooled trans-ethnic analysis for all prior telomere length GWAS sentinel variants with reported p-value $<5x10^{-8}$ in prior published studies.

| | | | | | Previously reported results | | | | | | | Single variant anlaysis in pooled trans-ethnic sample | | |
|---|---|---|---|---|---|---|---|---|---|---|---|---|---|---|
| Chromosome | Position (hg38) | rsID | Ref | Alt | Locus Name | First Author | Year | PMCID | P-Value | Direction | Near Table 1 Variant | P-value | Estimate (bp) | AAF |
| 1 | 226374920 | rs3219104 | A | C | PARP1 | Dorajoo | 2019 | PMC6554354 | 2.23E-16 | + | Yes | 1.27E-18 | 25.6 | 80.9% |
| 1 | 226374920 | rs3219104 | A | C | PARP1 | Li | 2020 | PMC7058826 | 9.31E-11 | + | Yes | 1.27E-18 | 25.6 | 80.9% |
| 2 | 54248729 | rs11125529 | C | A | ACYP2 | Codd | 2013 | PMC4006270 | 4.48E-08 | + | Yes | 8.82E-16 | 25.5 | 14.6% |
| 2 | 136258421 | rs4452212 | G | A | CXCR4 | Levy | 2010 | PMC2889047 | 2.94E-08 | N/A | No | 2.19E-01 | -3.4 | 32.7% |
| 3 | 58390292 | rs6772228 | T | A | PXK | Pooley | 2013 | PMC3836481 | 3.91E-10 | N/A | No | 6.38E-01 | -3.2 | 3.0% |
| 3 | 101513249 | rs55749605 | C | A | SENP7 | Li | 2020 | PMC7058826 | 2.38E-08 | - | No | 2.26E-02 | -5.6 | 69.7% |
| 3 | 169759718 | rs12638862 | A | G | TERC | Delgado | 2017 | PMC5749304 | 2.20E-08 | - | Yes | 2.24E-65 | -44.5 | 25.8% |
| 3 | 169763483 | rs12696304 | C | G | TERC | Prescott | 2011 | PMC3091863 | 1.50E-14 | N/A | Yes | 4.18E-44 | -33.2 | 39.9% |
| 3 | 169764547 | rs2293607 | T | C | TERC | Dorajoo | 2019 | PMC6554354 | 7.57E-39 | - | Yes | 3.19E-96 | -58.9 | 21.7% |
| 3 | 169774313 | rs10936599 | C | T | TERC | Codd | 2013 | PMC4006270 | 2.54E-31 | - | Yes | 8.15E-95 | -58.5 | 21.8% |
| 3 | 169779797 | rs1317082 | A | G | TERC | Mangino | 2012 | PMC3510758 | 1.14E-08 | N/A | Yes | 1.61E-95 | -58.8 | 21.8% |
| 3 | 169779797 | rs1317082 | A | G | TERC | Pooley | 2013 | PMC3836481 | 1.33E-19 | N/A | Yes | 1.61E-95 | -58.8 | 21.8% |
| 3 | 169796797 | rs10936600 | A | T | LRRC34 (TERC) | Li | 2020 | PMC7058826 | 6.42E-51 | - | Yes | 3.99E-86 | -54.5 | 22.6% |
| 3 | 169850328 | rs16847897 | G | C | TERC | Prescott | 2011 | PMC3091863 | 1.60E-13 | N/A | Yes | 1.54E-38 | -31.5 | 31.9% |
| 3 | 169862183 | rs1920116 | G | A | TERC | Walsh | 2014 | PMC4074274 | 8.30E-09 | - | Yes | 2.49E-46 | -37.5 | 26.3% |
| 4 | 70908630 | rs13137667 | T | C | MOB1B | Li | 2020 | PMC7058826 | 2.37E-08 | + | No | 5.81E-02 | 8.1 | 92.3% |
| 4 | 107383042 | rs7680468 | G | T | DKK2_PAPSS1 | Lee | 2014 | PMC3894567 | 4.70E-08 | N/A | No | 7.26E-01 | 2.3 | 2.8% |
| 4 | 163086668 | rs7675998 | A | G | NAF1 | Codd | 2013 | PMC4006270 | 4.35E-16 | + | Yes | 5.05E-22 | 26.3 | 78.4% |
| 4 | 163127047 | rs4691895 | G | C | NAF1 | Li | 2020 | PMC7058826 | 1.47E-21 | + | Yes | 3.48E-26 | 28.5 | 77.8% |
| 4 | 163180330 | rs10857352 | A | G | NAF1 | Dorajoo | 2019 | PMC6554354 | 4.85E-09 | + | Yes | 1.48E-21 | 21.7 | 45.7% |
| 5 | 1282204 | rs7726159 | C | A | TERT | Pooley | 2013 | PMC3836481 | 4.67E-17 | N/A | Yes | 4.04E-87 | 49.1 | 29.8% |
| 5 | 1285859 | rs7705526 | C | A | TERT | Dorajoo | 2019 | PMC6554354 | 2.61E-38 | + | Yes | 1.64E-92 | 50.9 | 29.4% |
| 5 | 1285859 | rs7705526 | C | A | TERT | Li | 2020 | PMC7058826 | 4.82E-45 | + | Yes | 1.64E-92 | 50.9 | 29.4% |
| 5 | 1286401 | rs2736100 | C | A | TERT | Codd | 2013 | PMC4006270 | 4.38E-19 | - | Yes | 5.27E-79 | -42.3 | 52.3% |
| 5 | 1286401 | rs2736100 | C | A | TERT | Walsh | 2014 | PMC4074274 | 1.40E-15 | - | Yes | 5.27E-79 | -42.3 | 52.3% |
| 5 | 1287079 | rs2853677 | G | A | TERT | Li | 2020 | PMC7058826 | 3.12E-31 | - | Yes | 8.17E-65 | -39.8 | 62.9% |
| 6 | 25480100 | rs34991172 | T | G | CARMIL1 | Li | 2020 | PMC7058826 | 6.03E-09 | - | No | 2.51E-01 | -7.3 | 3.4% |
| 6 | 31619784 | rs2736176 | G | C | PRRC2A | Li | 2020 | PMC7058826 | 3.41E-10 | + | Yes | 2.10E-09 | 16.3 | 24.1% |
| 7 | 124914213 | rs59294613 | C | A | POT1 | Li | 2020 | PMC7058826 | 1.12E-13 | - | Yes | 5.59E-25 | -25.7 | 28.0% |
| 7 | 124959695 | rs7776744 | A | G | POT1 | Dorajoo | 2019 | PMC6554354 | 2.51E-10 | - | Yes | 4.28E-13 | -16.3 | 53.8% |
| 8 | 73008648 | rs28365964 | T | C | TERF1 | Dorajoo | 2019 | PMC6554354 | 6.96E-15 | + | Yes | 1.46E-02 | 76.9 | 0.1% |
| 10 | 103900068 | rs2487999 | T | C | OBFC1 | Pooley | 2013 | PMC3836481 | 4.22E-14 | N/A | Yes | 3.14E-53 | -49.6 | 84.4% |
| 10 | 103916188 | rs9419958 | T | C | STN1 (OBFC1) | Li | 2020 | PMC7058826 | 4.77E-19 | - | Yes | 9.05E-83 | -54.6 | 74.9% |
| 10 | 103916188 | rs9419958 | T | C | OBFC1 | Mangino | 2012 | PMC3510758 | 9.13E-11 | N/A | Yes | 9.05E-83 | -54.6 | 74.9% |
| 10 | 103916707 | rs9420907 | C | A | OBFC1 | Codd | 2013 | PMC4006270 | 6.90E-11 | - | Yes | 3.90E-83 | -54.7 | 74.9% |
| 10 | 103918139 | rs4387287 | A | C | OBFC1 | Levy | 2010 | PMC2889047 | 3.87E-09 | N/A | Yes | 2.35E-77 | -49.7 | 67.8% |
| 10 | 103920828 | rs12415148 | T | C | OBFC1 | Dorajoo | 2019 | PMC6554354 | 2.78E-25 | + | Yes | 1.04E-23 | 107.5 | 1.1% |
| 11 | 108234866 | rs228595 | G | A | ATM | Li | 2020 | PMC7058826 | 1.39E-08 | - | Yes | 2.12E-13 | -17.7 | 36.2% |
| 11 | 108377161 | rs227080 | A | G | ATM | Dorajoo | 2019 | PMC6554354 | 1.87E-10 | - | Yes | 1.27E-14 | -17.5 | 53.7% |
| 14 | 24252121 | rs41293836 | C | T | TINF2 | Dorajoo | 2019 | PMC6554354 | 2.47E-42 | + | Yes | 6.49E-08 | 47.6 | 1.6% |
| 14 | 72938044 | rs2302588 | G | C | DCAF4 | Li | 2020 | PMC7058826 | 1.64E-08 | + | Yes | 5.76E-10 | 23.4 | 9.7% |
| 14 | 72948525 | rs2535913 | G | A | DCAF4 | Mangino | 2015 | PMC4345921 | 6.38E-10 | N/A | Yes | 1.77E-09 | -15.8 | 24.7% |
| 16 | 58175370 | rs74019828 | G | A | CSNK2A2 | Saxena | 2014 | PMC4106467 | 4.52E-08 | - | No | 3.62E-01 | -4.2 | 6.3% |
| 16 | 69373083 | rs3785074 | A | G | TERF2 | Li | 2020 | PMC7058826 | 4.50E-10 | + | Yes | 4.73E-14 | 18.4 | 30.8% |
| 16 | 74646176 | rs62053580 | A | G | RFWD3 | Li | 2020 | PMC7058826 | 3.96E-08 | - | Yes | 3.67E-11 | -22.1 | 13.2% |
| 16 | 82166375 | rs7194734 | C | T | MPHOSPH6 | Li | 2020 | PMC7058826 | 6.72E-10 | - | Yes | 3.31E-13 | -20.8 | 80.9% |
| 17 | 8232774 | rs3027234 | C | T | CTC1 | Mangino | 2012 | PMC3510758 | 2.29E-08 | N/A | No | 7.97E-05 | -12.6 | 15.2% |
| 19 | 22032639 | rs8105767 | A | G | ZNF208 | Codd | 2013 | PMC4006270 | 1.11E-09 | + | Yes | 3.59E-18 | 20.6 | 35.1% |
| 19 | 22032639 | rs8105767 | A | G | ZNF208 | Li | 2020 | PMC7058826 | 5.21E-13 | + | Yes | 3.59E-18 | 20.6 | 35.1% |
| 19 | 22176638 | rs412658 | C | T | ZNF676 | Mangino | 2012 | PMC3510758 | 9.75E-09 | N/A | Yes | 5.15E-07 | 11.6 | 42.7% |
| 20 | 63638397 | rs75691080 | C | T | RTEL1/STMN3 | Li | 2020 | PMC7058826 | 5.75E-14 | - | Yes | 6.92E-17 | -33.5 | 8.5% |
| 20 | 63660246 | rs34978822 | C | G | RTEL1 | Li | 2020 | PMC7058826 | 7.04E-10 | - | Yes | 1.78E-28 | -122.5 | 1.1% |
| 20 | 63678201 | rs41309367 | C | T | RTEL1 | Dorajoo | 2019 | PMC6554354 | 1.16E-08 | - | Yes | 2.52E-33 | -29.1 | 67.3% |
| 20 | 63678486 | rs6010620 | A | G | RTEL1 | Walsh | 2014 | PMC4074274 | 4.70E-19 | - | Yes | 3.13E-10 | -18.5 | 79.1% |
| 20 | 63790269 | rs755017 | A | G | RTEL1 | Codd | 2013 | PMC4006270 | 6.71E-09 | + | Yes | 3.58E-05 | 11.9 | 19.6% |
| 20 | 63805045 | rs73624724 | T | C | RTEL1/ZBTB46 | Li | 2020 | PMC7058826 | 6.08E-12 | + | Yes | 1.65E-04 | 10.9 | 19.3% |

**Table S4, Related to Table 1:** Association results for the single variant association analysis for each of the 59 sentinel variants from ***Table 1*** . Single variant results are shown for the pooled trans-ethnic analysis, and each population group. Variants with a minor allele count <5 were not included in the analysis and are listed as "-".

| | | | | Est (in bp) from the single variant analysis by stratum | | | | | P-value from the single variant analysis by stratum | | | | | Alternate allele frequency | | | | | Percent variation explained | | | | |
|---|---|---|---|---|---|---|---|---|---|---|---|---|---|---|---|---|---|---|---|---|---|---|---|
| Chr | Position | Locus | rsNum | Trans-ethnic | European | African | Hispanic /Latino | Asian | Trans-ethnic | European | African | Hispanic /Latino | Asian | Trans-ethnic | European | African | Hispanic /Latino | Asian | Trans-ethnic | European | African | Hispanic /Latino | Asian |
| 1 | 35259602 | ZMYM4 | rs11581846 | -19.1 | -21.6 | -11.8 | -25.4 | -6.1 | 3.04E-10 | 7.72E-05 | 4.31E-02 | 3.18E-06 | 5.73E-01 | 59.9% | 89.6% | 19.4% | 57.8% | 23.4% | 0.036% | 0.030% | 0.014% | 0.121% | 0.006% |
| 1 | 113881455 | BCL2L15 | rs2296176 | -19.7 | -22.1 | -22.9 | -9.5 | -23.9 | 2.84E-10 | 1.82E-07 | 1.93E-03 | 2.06E-01 | 4.38E-02 | 15.2% | 19.1% | 9.2% | 13.9% | 16.0% | 0.036% | 0.053% | 0.033% | 0.009% | 0.072% |
| 1 | 226367601 | PARP1 | rs1136410 | -28.5 | -27.0 | -19.5 | -26.3 | -32.2 | 9.32E-20 | 3.52E-09 | 3.79E-02 | 1.84E-05 | 3.11E-04 | 16.7% | 15.7% | 5.5% | 27.6% | 41.7% | 0.076% | 0.068% | 0.015% | 0.102% | 0.230% |
| 2 | 54255416 | ACYP2 | rs17189743 | -58.9 | -57.5 | -28.7 | -82.6 | -37.8 | 7.18E-12 | 1.84E-07 | 2.89E-01 | 9.38E-06 | 3.04E-01 | 1.7% | 2.3% | 0.6% | 1.9% | 1.4% | 0.043% | 0.053% | 0.004% | 0.109% | 0.019% |
| 2 | 54263623 | ACYP2 | rs144980386 | 28.2 | 33.7 | 21.5 | 30.2 | 5.9 | 1.32E-17 | 1.92E-12 | 7.80E-04 | 2.50E-04 | 6.15E-01 | 13.2% | 14.1% | 12.8% | 11.2% | 16.9% | 0.067% | 0.096% | 0.039% | 0.075% | 0.004% |
| 3 | 117584223 | LINC00901 | rs961617801 | 1063.3 | 1082.0 | - | - | - | 1.25E-11 | 1.33E-10 | - | - | - | 0.01% | 0.01% | - | - | - | 0.042% | 0.080% | - | - | - |
| 3 | 169769649 | TERC | rs12637184 | -59.1 | -56.6 | -62.4 | -65.8 | -55.1 | 1.30E-96 | 6.32E-48 | 1.35E-13 | 4.36E-28 | 2.94E-10 | 21.7% | 24.1% | 7.1% | 29.5% | 52.9% | 0.399% | 0.410% | 0.188% | 0.671% | 0.702% |
| 3 | 169772313 | TERC | rs9826466 | -70.3 | 307.0 | -71.6 | -66.2 | - | 3.25E-17 | 1.33E-01 | 9.16E-16 | 1.38E-02 | - | 1.9% | 0.01% | 6.1% | 0.9% | - | 0.065% | 0.004% | 0.221% | 0.034% | - |
| 3 | 190053412 | P3H2 | rs10937417 | 14.5 | 12.8 | 18.1 | 17.2 | -3.7 | 6.89E-10 | 2.97E-04 | 2.03E-05 | 2.85E-03 | 6.91E-01 | 36.8% | 33.3% | 49.1% | 29.4% | 31.2% | 0.035% | 0.025% | 0.062% | 0.050% | 0.003% |
| 4 | 9928595 | SLC2A2 | rs4235345 | 16.9 | 19.0 | 12.4 | 15.1 | 48.5 | 3.82E-09 | 1.64E-06 | 5.66E-02 | 8.70E-03 | 5.44E-02 | 20.3% | 23.0% | 12.5% | 31.5% | 3.5% | 0.032% | 0.045% | 0.012% | 0.038% | 0.065% |
| 4 | 163126692 | NAF1 | rs60735607 | -7.4 | -9.0 | -2.2 | -10.9 | 6.6 | 3.96E-03 | 1.97E-02 | 6.23E-01 | 8.81E-02 | 7.32E-01 | 26.3% | 24.8% | 36.3% | 21.1% | 5.5% | 0.008% | 0.011% | 0.001% | 0.016% | 0.002% |
| 4 | 163144568 | NAF1 | rs113580095 | -273.5 | -254.3 | -331.8 | -278.3 | - | 4.72E-18 | 2.07E-09 | 4.09E-02 | 7.66E-08 | - | 0.1% | 0.2% | 0.02% | 0.3% | - | 0.069% | 0.070% | 0.014% | 0.161% | - |
| 4 | 163155406 | NAF1 | rs1351222 | 28.2 | 28.5 | 23.5 | 36.7 | 30.8 | 3.27E-26 | 4.56E-13 | 4.04E-06 | 3.11E-09 | 4.18E-03 | 77.0% | 76.4% | 77.3% | 77.5% | 79.0% | 0.103% | 0.102% | 0.073% | 0.195% | 0.145% |
| 5 | 1272383 | TERT | rs192999400 | 82.1 | 81.7 | 81.0 | 75.7 | 95.6 | 3.10E-15 | 2.64E-01 | 8.22E-13 | 2.41E-02 | 1.05E-01 | 1.1% | 0.0% | 3.6% | 0.6% | 0.5% | 0.057% | 0.002% | 0.175% | 0.028% | 0.046% |
| 5 | 1280823 | TERT | rs6897196 | 44.5 | 43.3 | 38.7 | 44.9 | 65.0 | 1.87E-83 | 2.47E-37 | 2.80E-18 | 1.27E-16 | 2.45E-13 | 46.8% | 40.1% | 63.0% | 38.6% | 39.4% | 0.344% | 0.316% | 0.260% | 0.381% | 0.948% |
| 5 | 1285859 | TERT | rs7705526 | 50.9 | 52.8 | 35.0 | 53.0 | 67.0 | 1.64E-92 | 1.29E-50 | 1.65E-10 | 5.22E-20 | 3.65E-14 | 29.4% | 33.5% | 18.8% | 28.5% | 40.8% | 0.382% | 0.434% | 0.140% | 0.467% | 1.014% |
| 5 | 1287079 | TERT | rs2853677 | -39.8 | -33.3 | -38.7 | -47.6 | -61.8 | 8.17E-65 | 2.76E-23 | 8.35E-16 | 1.59E-17 | 3.10E-12 | 62.9% | 56.9% | 72.7% | 67.0% | 60.6% | 0.265% | 0.191% | 0.222% | 0.404% | 0.860% |
| 5 | 1292331 | TERT | rs34052286 | -48.5 | -60.1 | -45.5 | -51.2 | - | 1.50E-12 | 3.90E-01 | 2.26E-09 | 4.33E-03 | - | 2.8% | 0.1% | 8.6% | 2.1% | - | 0.046% | 0.001% | 0.122% | 0.045% | - |
| 5 | 1292843 | TERT | rs114616103 | -40.7 | -47.9 | -17.3 | -44.3 | -28.1 | 2.39E-07 | 1.30E-06 | 2.79E-01 | 7.08E-02 | 7.93E-01 | 2.1% | 2.9% | 1.8% | 1.1% | 0.2% | 0.024% | 0.045% | 0.004% | 0.018% | 0.001% |
| 5 | 139637905 | CXXC5 | rs75903170 | 30.4 | 23.5 | 48.6 | 27.4 | 13.9 | 5.69E-10 | 1.02E-03 | 1.56E-06 | 3.68E-02 | 3.92E-01 | 5.5% | 5.6% | 4.6% | 4.0% | 7.7% | 0.035% | 0.021% | 0.079% | 0.024% | 0.013% |
| 6 | 31815431 | HSPA1A | rs1008438 | -19.8 | -18.6 | -18.5 | -25.6 | -17.8 | 3.42E-17 | 5.22E-08 | 1.68E-04 | 8.89E-07 | 4.23E-02 | 51.0% | 38.0% | 73.9% | 52.2% | 44.4% | 0.065% | 0.057% | 0.048% | 0.134% | 0.073% |
| 7 | 124812616 | POT1 | rs720613 | -26.4 | -31.9 | -19.4 | -26.8 | -19.3 | 1.27E-26 | 3.17E-18 | 2.54E-05 | 1.17E-05 | 4.24E-02 | 28.6% | 29.1% | 30.9% | 24.0% | 28.6% | 0.105% | 0.147% | 0.061% | 0.107% | 0.073% |
| 7 | 124858989 | POT1 | rs202187871 | 727.2 | 710.0 | - | - | - | 4.89E-12 | 3.61E-11 | - | - | - | 0.01% | 0.03% | - | - | - | 0.044% | 0.085% | - | - | - |
| 7 | 129041243 | TNP03 | rs7783384 | -15.9 | -18.3 | -14.0 | -13.3 | -19.7 | 2.34E-12 | 1.03E-07 | 9.75E-04 | 1.07E-02 | 2.39E-02 | 57.6% | 62.6% | 53.5% | 54.0% | 50.9% | 0.045% | 0.055% | 0.037% | 0.036% | 0.090% |
| 8 | 73004218 | TERF1 | rs183633026 | 101.9 | 73.1 | -7.0 | 112.0 | - | 1.59E-10 | 4.49E-02 | 9.26E-01 | 1.85E-09 | - | 0.5% | 0.2% | 0.1% | 2.2% | - | 0.038% | 0.008% | 0.000% | 0.201% | - |
| 8 | 73033303 | TERF1 | rs73687065 | 88.9 | 29.8 | 93.0 | 79.4 | - | 3.10E-12 | 7.37E-01 | 3.73E-11 | 2.79E-02 | - | 0.8% | 0.03% | 2.3% | 0.5% | - | 0.045% | 0.000% | 0.150% | 0.027% | - |
| 8 | 73046483 | TERF1 | rs10112752 | -17.1 | -18.7 | -14.8 | -16.4 | -26.9 | 4.59E-13 | 2.35E-08 | 1.31E-03 | 3.77E-03 | 2.67E-02 | 36.2% | 43.7% | 30.8% | 32.0% | 15.6% | 0.048% | 0.060% | 0.035% | 0.047% | 0.087% |
| 10 | 94344908 | NOC3L | rs3758526 | -21.5 | -20.2 | -21.9 | -16.8 | -26.5 | 6.80E-12 | 5.83E-05 | 6.94E-05 | 4.16E-02 | 5.88E-03 | 14.9% | 12.5% | 18.2% | 11.1% | 28.1% | 0.043% | 0.031% | 0.054% | 0.023% | 0.134% |
| 10 | 99514276 | NKX2-3 | rs10883359 | -18.1 | -14.9 | -34.5 | -12.7 | -13.5 | 3.60E-12 | 5.15E-05 | 3.75E-08 | 2.73E-02 | 1.27E-01 | 25.2% | 28.6% | 13.3% | 29.5% | 41.3% | 0.044% | 0.032% | 0.104% | 0.027% | 0.041% |
| 10 | 103907794 | OBFC1 | rs10883948 | -28.0 | -25.8 | -49.0 | -14.8 | 3.5 | 2.04E-34 | 8.05E-15 | 8.43E-27 | 4.61E-03 | 7.09E-01 | 43.3% | 50.1% | 31.7% | 46.9% | 33.1% | 0.137% | 0.117% | 0.393% | 0.045% | 0.002% |
| 10 | 103915847 | OBFC1 | rs112163720 | 3.3 | 32.9 | -12.8 | 8.8 | 10.7 | 4.40E-01 | 8.78E-04 | 5.36E-02 | 3.60E-01 | 3.77E-01 | 7.1% | 2.9% | 11.6% | 8.2% | 14.9% | 0.001% | 0.021% | 0.013% | 0.005% | 0.014% |
| 10 | 103916707 | OBFC1 | rs9420907 | -54.7 | -51.3 | -59.3 | -57.1 | -0.1 | 3.90E-83 | 8.24E-27 | 4.41E-43 | 1.13E-16 | 9.98E-01 | 74.9% | 85.9% | 47.8% | 81.8% | 98.0% | 0.342% | 0.223% | 0.648% | 0.382% | 0.000% |
| 10 | 103918153 | OBFC1 | rs111447985 | 109.8 | 97.3 | 48.9 | 127.9 | 94.1 | 2.29E-24 | 2.65E-02 | 2.63E-01 | 9.14E-18 | 5.30E-07 | 1.1% | 0.1% | 0.2% | 3.5% | 5.9% | 0.095% | 0.010% | 0.004% | 0.410% | 0.445% |
| 11 | 108158382 | ATM | rs61380955 | -19.3 | -23.9 | -15.5 | -16.6 | -14.7 | 2.47E-17 | 1.45E-12 | 4.16E-04 | 1.88E-03 | 9.65E-02 | 51.4% | 57.5% | 38.1% | 59.6% | 46.7% | 0.066% | 0.097% | 0.043% | 0.054% | 0.049% |
| 13 | 41150640 | KBTBD7 | rs1411041 | 22.1 | 24.9 | 21.3 | 21.7 | 17.4 | 6.29E-14 | 6.16E-08 | 4.87E-04 | 6.57E-04 | 5.16E-02 | 60.8% | 84.5% | 17.1% | 75.7% | 39.3% | 0.052% | 0.057% | 0.042% | 0.065% | 0.067% |
| 14 | 24242592 | TINF2 | rs28372734 | 108.8 | 117.6 | 95.7 | 126.7 | 94.5 | 1.74E-27 | 6.07E-02 | 2.75E-05 | 4.51E-08 | 1.75E-09 | 1.2% | 0.1% | 0.9% | 1.3% | 8.2% | 0.108% | 0.007% | 0.060% | 0.166% | 0.641% |
| 14 | 24243052 | TINF2 | rs8016076 | 79.5 | 429.9 | 76.9 | 83.0 | - | 1.80E-11 | 6.51E-02 | 2.79E-09 | 1.25E-02 | - | 0.9% | 0.0% | 2.8% | 0.6% | - | 0.041% | 0.007% | 0.121% | 0.035% | - |
| 14 | 24254544 | TINF2 | rs41293824 | 80.9 | 60.7 | 73.4 | 124.4 | - | 1.31E-09 | 6.36E-01 | 9.37E-07 | 4.55E-04 | - | 0.7% | 0.0% | 2.1% | 0.5% | - | 0.034% | 0.000% | 0.082% | 0.068% | - |
| 14 | 72959582 | DCAF4 | rs2572 | 26.4 | 26.6 | 35.2 | 21.0 | 33.6 | 5.14E-12 | 1.04E-06 | 2.50E-04 | 1.89E-02 | 2.27E-03 | 9.5% | 10.3% | 5.2% | 8.9% | 19.5% | 0.044% | 0.046% | 0.046% | 0.031% | 0.165% |
| 15 | 50065546 | ATP8B4 | rs7172615 | -17.0 | -19.7 | -21.1 | -7.2 | -11.2 | 4.31E-09 | 4.64E-07 | 5.19E-03 | 3.00E-01 | 2.30E-01 | 18.8% | 23.3% | 8.8% | 16.8% | 31.7% | 0.032% | 0.049% | 0.027% | 0.006% | 0.025% |
| 16 | 69357811 | TERF2 | rs9925619 | 18.6 | 14.4 | 22.2 | 25.9 | 7.4 | 3.01E-14 | 1.10E-04 | 2.63E-07 | 1.04E-05 | 5.88E-01 | 30.3% | 27.5% | 40.9% | 26.6% | 12.0% | 0.053% | 0.029% | 0.091% | 0.108% | 0.005% |
| 16 | 70193527 | CLEC18C | rs62049363 | -16.2 | -16.3 | -11.3 | -17.9 | -10.7 | 4.09E-10 | 1.68E-06 | 1.49E-01 | 1.78E-03 | 2.48E-01 | 28.3% | 39.2% | 8.6% | 30.6% | 30.5% | 0.036% | 0.044% | 0.007% | 0.054% | 0.024% |
| 16 | 74630845 | RFWD3 | rs7193541 | -18.7 | -23.5 | -17.6 | -13.3 | -5.6 | 1.47E-16 | 2.57E-12 | 3.38E-05 | 1.60E-02 | 5.25E-01 | 42.6% | 41.6% | 50.1% | 34.1% | 41.5% | 0.063% | 0.095% | 0.059% | 0.032% | 0.007% |
| 16 | 82166498 | MPHOSPH6 | rs2967355 | -27.9 | -25.6 | -32.2 | -37.7 | 1.0 | 3.96E-19 | 2.00E-10 | 4.19E-04 | 5.10E-07 | 9.33E-01 | 84.0% | 78.0% | 94.1% | 85.8% | 83.1% | 0.073% | 0.078% | 0.043% | 0.140% | 0.000% |
| 16 | 87961594 | BANP | rs12934497 | 14.9 | 13.8 | 16.1 | 16.8 | 34.3 | 9.15E-10 | 3.57E-05 | 1.06E-03 | 2.97E-03 | 9.59E-02 | 35.3% | 46.5% | 25.4% | 31.8% | 4.8% | 0.034% | 0.033% | 0.037% | 0.049% | 0.049% |
| 18 | 650764 | TYMS | rs150119891 | -76.8 | -77.6 | -41.2 | -138.4 | - | 2.27E-07 | 2.12E-06 | 4.03E-01 | 1.27E-02 | - | 0.6% | 1.1% | 0.2% | 0.2% | - | 0.025% | 0.044% | 0.002% | 0.035% | - |
| 18 | 666625 | TYMS | rs8088781 | -27.7 | -28.3 | -27.2 | -28.0 | -4.9 | 2.49E-15 | 2.46E-09 | 3.51E-05 | 2.54E-03 | 9.21E-01 | 11.8% | 14.4% | 11.9% | 8.4% | 0.8% | 0.057% | 0.069% | 0.059% | 0.051% | 0.000% |
| 18 | 676473 | TYMS | rs2612101 | 2.2 | -8.5 | 5.5 | 18.5 | 45.9 | 4.08E-01 | 3.07E-02 | 2.20E-01 | 1.44E-03 | 2.88E-01 | 25.7% | 23.3% | 34.2% | 27.9% | 1.1% | 0.001% | 0.009% | 0.005% | 0.056% | 0.020% |
| 18 | 44666476 | SETBP1 | rs2852770 | -18.6 | -25.9 | -7.8 | -24.0 | -6.0 | 1.00E-11 | 1.60E-09 | 1.01E-01 | 2.25E-04 | 5.79E-01 | 21.3% | 18.6% | 27.0% | 20.3% | 20.0% | 0.042% | 0.071% | 0.009% | 0.076% | 0.005% |
| 19 | 22032639 | ZNF257_ZNF | rs8105767 | 20.6 | 21.6 | 23.3 | 12.7 | 26.7 | 3.59E-18 | 3.63E-09 | 4.85E-08 | 2.15E-02 | 4.03E-03 | 35.1% | 29.1% | 47.1% | 34.6% | 32.2% | 0.069% | 0.067% | 0.102% | 0.029% | 0.146% |
| 20 | 36922795 | SAMHD1 | rs2342113 | -23.3 | -33.8 | -15.3 | -26.1 | -7.9 | 4.62E-18 | 7.52E-13 | 9.76E-04 | 7.99E-06 | 3.66E-01 | 23.7% | 14.5% | 30.0% | 30.8% | 46.9% | 0.069% | 0.100% | 0.037% | 0.111% | 0.014% |
| 20 | 63661765 | RTEL1 | rs41308088 | 29.5 | 36.2 | 14.0 | 16.6 | 59.7 | 1.58E-10 | 2.75E-10 | 3.04E-01 | 1.00E-01 | 1.92E-01 | 6.5% | 9.2% | 2.5% | 7.3% | 1.1% | 0.038% | 0.077% | 0.004% | 0.015% | 0.030% |
| 20 | 63676585 | RTEL1 | rs79981941 | -37.6 | -40.3 | -28.4 | -47.7 | -51.3 | 1.46E-23 | 1.65E-12 | 1.43E-05 | 2.80E-08 | 6.40E-02 | 9.9% | 9.3% | 12.0% | 10.1% | 2.9% | 0.092% | 0.097% | 0.064% | 0.171% | 0.061% |
| 20 | 63678201 | RTEL1 | rs41309367 | -29.1 | -26.2 | -37.8 | -34.0 | -7.0 | 2.52E-33 | 2.55E-13 | 8.79E-16 | 1.67E-09 | 4.73E-01 | 67.3% | 68.8% | 70.7% | 69.1% | 30.1% | 0.133% | 0.104% | 0.221% | 0.202% | 0.009% |
| 20 | 63689775 | RTEL1 | rs35640778 | -120.9 | -121.4 | -143.7 | -95.6 | 5.2 | 2.31E-28 | 1.00E-21 | 3.07E-05 | 1.90E-03 | 9.72E-01 | 1.1% | 1.7% | 0.4% | 0.7% | 0.1% | 0.112% | 0.178% | 0.059% | 0.054% | 0.000% |
| 20 | 63695521 | RTEL1 | rs181080831 | 174.0 | 190.3 | 12.5 | 142.0 | - | 6.40E-17 | 9.02E-16 | 8.81E-01 | 1.07E-02 | - | 0.3% | 0.5% | 0.1% | 0.2% | - | 0.064% | 0.125% | 0.000% | 0.036% | - |
| 22 | 50532618 | TYMP | rs361725 | -15.0 | -21.2 | -5.0 | -11.3 | -20.1 | 1.36E-10 | 5.91E-10 | 2.86E-01 | 3.31E-02 | 3.55E-02 | 53.4% | 63.0% | 30.9% | 59.0% | 69.8% | 0.038% | 0.074% | 0.004% | 0.025% | 0.078% |
| X | 66015290 | VSIG4 | rs12394264 | 18.6 | 18.5 | 15.3 | 22.2 | 7.7 | 9.67E-16 | 7.46E-07 | 4.58E-05 | 1.76E-05 | 8.44E-01 | 29.8% | 16.0% | 62.3% | 23.8% | 0.9% | 0.059% | 0.047% | 0.057% | 0.102% | 0.001% |
| X | 154720412 | GAB3 | rs2728723 | 13.8 | 16.5 | 7.1 | 14.4 | 16.6 | 1.21E-12 | 1.86E-08 | 4.96E-02 | 1.10E-03 | 5.50E-02 | 62.8% | 67.9% | 52.2% | 60.0% | 79.8% | 0.046% | 0.061% | 0.013% | 0.059% | 0.065% |

**Table S5, Related to Figure 4:** Iterative conditional analysis was repeated on chromosome 10 focusing exclusively on the *OBFC1* locus, defined as a 2Mb window around the original top sentinel, rs9420907. The sentinel for each signal was consistent with the iterative conditional analysis performed on the entirety of chromosome 10 (***Table S2***). The summary statistics from each iterative conditional analysis were used to perform colocalization analysis on the non-primary signals. Colocalization analysis was performed using coloc with all significant gene-tissue pairs in GTEx and with all genes in a 2Mb window around rs9420907 in eQTLGen. All results with PPH4 > 0.7 for each signal are reported.

| chr | position | rsNum | Round | r2 | D' | Primary | | Conditional Round 1 | | Conditional Round 2 | | Conditional Round 3 | | coloc results with GTEx (PPH4 > 0.7) | | | | |
|---|---|---|---|---|---|---|---|---|---|---|---|---|---|---|---|---|---|---|
| | | | | | | Est | p-value | Est | p-value | Est | p-value | Est | p-value | Gene | Gene Symbol | Tissue | PPH3 | PPH4 |
| 10 | 103916707 | rs9420907 | Primary | | | **-54.70** | **3.90E-83** | | | | | | | ENSG00000260461.1 | RP11-541N10.3 | Adipose_Subcutaneous | 0.03 | 0.95 |
| | | | | | | | | | | | | | | ENSG00000107960.10 | OBFC1 | Adipose_Subcutaneous | 0.08 | 0.9 |
| | | | | | | | | | | | | | | ENSG00000260461.1 | RP11-541N10.3 | Colon_Sigmoid | 0.13 | 0.72 |
| | | | | | | | | | | | | | | ENSG00000107960.10 | OBFC1 | Colon_Transverse | 0.08 | 0.92 |
| | | | | | | | | | | | | | | ENSG00000107960.10 | OBFC1 | Esophagus_Mucosa | 0.03 | 0.97 |
| | | | | | | | | | | | | | | ENSG00000107960.10 | OBFC1 | Esophagus_Muscularis | 0.05 | 0.94 |
| | | | | | | | | | | | | | | ENSG00000107960.10 | OBFC1 | Colon_Sigmoid | 0.16 | 0.81 |
| | | | | | | | | | | | | | | ENSG00000107960.10 | OBFC1 | Skin_Not_Sun_Exposed_Suprapubic | 0.03 | 0.97 |
| | | | | | | | | | | | | | | ENSG00000107960.10 | OBFC1 | Skin_Sun_Exposed_Lower_leg | 0.02 | 0.98 |
| | | | | | | | | | | | | | | ENSG00000107960.10 | OBFC1 | Spleen | 0.1 | 0.79 |
| | | | | | | | | | | | | | | ENSG00000107960.10 | OBFC1 | Nerve_Tibial | 0.07 | 0.92 |
| 10 | 103918153 | rs111447985 | Secondary | 0.004 | 1.00 | 109.76 | 2.29E-24 | **116.56** | **2.72E-27** | | | | | N/A (rare variant, not available in GTEx or eQTLGen) | | | | |
| 10 | 103915847 | rs112163720 | Tertiary | 0.026 | 1 | 3.35 | 0.440005 | 24.10 | 6.29E-08 | **26.90** | **1.47E-09** | | | ENSG00000107960.10 | OBFC1 | Muscle_Skeletal | 0.16 | 0.84 |
| 10 | 103907794 | rs10883948 | Quaternary | 0.246 | 0.98 | -28.00 | 2.04E-34 | -8.30 | 1.52E-03 | -12.20 | 3.81E-06 | **-18.80** | **1.14E-11** | ENSG00000260461.1 | RP11-541N10.3 | Brain_Cerebellar_Hemisphere | 0.25 | 0.73 |
| | | | | | | | | | | | | | | ENSG00000260461.1 | RP11-541N10.3 | Brain_Cerebellum | 0.26 | 0.74 |
| | | | | | | | | | | | | | | ENSG00000260461.1 | RP11-541N10.3 | Brain_Cortex | 0.24 | 0.72 |
| | | | | | | | | | | | | | | ENSG00000260461.1 | RP11-541N10.3 | Brain_Frontal_Cortex_BA9 | 0.25 | 0.75 |
| | | | | | | | | | | | | | | ENSG00000260461.1 | RP11-541N10.3 | Esophagus_Mucosa | 0.23 | 0.77 |
| | | | | | | | | | | | | | | ENSG00000065613.13 | SLK | Muscle_Skeletal | 0.24 | 0.72 |
| | | | | | | | | | | | | | | ENSG00000065613.13 | SLK | Ovary | 0.25 | 0.75 |
| | | | | | | | | | | | | | | ENSG00000107960.10 | OBFC1 | Whole_Blood | 0.25 | 0.75 |

**Table S6, Related to Figure 5:** Results of the PheWAS performed on 49 available sentinel variants and a polygenic trait score across these 49 sentinel variants using BioVU self-identified African Americans (AA, n=15,174) and BioVU self-identified European Americans (EA, n=70,439). The cumulative TL risk score for the BioVU AA and EA samples was derived from the African and European specific effects sizes from ***Table 1*** , respectively. Results were evaluated at a Bonferroni threshold corrected for the number of informative phecodes for each variant.

| Chr | Pos | Locus | rsNum | Ref | Alt | Trans-ethnic TL Effect size in (bp) | PheWAS Results: Phecode | Category | Phenotype | OR | P-Value | Number of Samples | Ancestry |
|---|---|---|---|---|---|---|---|---|---|---|---|---|---|
| 49 SNP EA PTS | | | | | | | 165 | neoplasms | Cancer within the respiratory system | 1.104 | 5.76E-06 | 2465 / 66885 | EA |
| | | | | | | | 165.1 | neoplasms | Cancer of bronchus; lung | 1.107 | 3.92E-06 | 2444 / 66885 | |
| | | | | | | | 172.1 | neoplasms | Melanomas of skin, dx or hx | 1.140 | 1.35E-06 | 1461 / 59520 | |
| | | | | | | | 172.11 | neoplasms | Melanomas of skin | 1.135 | 1.93E-05 | 1209 / 59520 | |
| | | | | | | | 189.1 | neoplasms | Cancer of kidney and renal pelvis | 1.162 | 4.61E-06 | 980 / 68020 | |
| | | | | | | | 191 | neoplasms | Manlignant and unknown neoplasms of brain and nervous system | 1.166 | 1.29E-05 | 838 / 67856 | |
| | | | | | | | 191.1 | neoplasms | Cancer of brain and nervous system | 1.223 | 4.62E-07 | 657 / 67856 | |
| | | | | | | | 191.11 | neoplasms | Cancer of brain | 1.243 | 2.16E-07 | 592 / 67856 | |
| | | | | | | | 198 | neoplasms | Secondary malignant neoplasm | 1.072 | 2.70E-06 | 6574 / 52740 | |
| | | | | | | | 198.1 | neoplasms | Secondary malignancy of lymph nodes | 1.085 | 2.63E-05 | 3381 / 52740 | |
| | | | | | | | 199 | neoplasms | Neoplasm of uncertain behavior | 1.100 | 1.43E-05 | 2254 / 52740 | |
| | | | | | | | 241 | endocrine/metabolic | Nontoxic nodular goiter | 1.108 | 1.45E-05 | 1909 / 56362 | |
| | | | | | | | 241.1 | endocrine/metabolic | Nontoxic uninodular goiter | 1.161 | 8.50E-06 | 925 / 56362 | |
| | | | | | | | 571.51 | digestive | Cirrhosis of liver without mention of alcohol | 0.896 | 1.20E-06 | 2062 / 56630 | |
| 3 | 169769649 | TERC | rs12637184 | G | A | -59.79 | 782.6 | symptoms | Pallor and flushing | 6.329 | 2.68E-06 | 21 / 12880 | AA |
| 4 | 9928595 | SLC2A2 | rs4235345 | G | A | 16.78 | 274.1 | endocrine/metabolic | Gout | 0.649 | 9.03E-24 | 2095 / 66779 | EA |
| | | | | | | | 274 | endocrine/metabolic | Gout and other crystal arthropathies | 0.671 | 9.57E-22 | 2186 / 66779 | EA |
| | | | | | | | 274.11 | endocrine/metabolic | Gouty arthropathy | 0.607 | 4.06E-11 | 685 / 66779 | EA |
| 5 | 1272383 | TERT | rs192999400 | C | T | 101.32 | 573.3 | digestive | Hepatomegaly | 4.619 | 2.71E-06 | 59 / 12759 | AA |
| | | | | | | | 742.1 | musculoskeletal | Loose body in joint | 212.237 | 3.16E-06 | 42 / 59012 | EA |
| | | | | | | | 772 | symptoms | Symptoms of the muscles | 37.798 | 1.01E-05 | 357 / 61619 | EA |
| | | | | | | | 199.4 | neoplasms | Neurofibromatosis | 179.327 | 1.10E-05 | 71 / 52740 | EA |
| | | | | | | | 286.13 | hematopoietic | Congenital factor VIII disorder | 163.406 | 2.89E-05 | 80 / 55029 | EA |
| 5 | 1285859 | TERT | rs7705526 | C | A | 30.04 | 200.1 | neoplasms | Polycythemia vera | 1.720 | 4.88E-08 | 214 / 59260 | EA |
| | | | | | | | 200 | neoplasms | Myeloproliferative disease | 1.287 | 9.32E-08 | 1006 / 67218 | EA |
| | | | | | | | 191.11 | neoplasms | Cancer of brain | 1.349 | 9.53E-07 | 592 / 67856 | EA |
| | | | | | | | 191.1 | neoplasms | Cancer of brain and nervous system | 1.301 | 5.96E-06 | 657 / 67856 | EA |
| 5 | 1287079 | TERT | rs2853677 | G | A | -23.89 | 200.1 | neoplasms | Polycythemia vera | 0.576 | 2.06E-08 | 214 / 59260 | EA |
| 5 | 139637905 | CXXC5 | rs75903170 | T | A | 29.55 | 603.2 | genitourinary | Spermatocele | 3.164 | 1.31E-05 | 59 / 61894 | EA |
| 6 | 31815431 | HSPA1A | rs1008438 | A | C | -20.33 | 250.1 | endocrine/metabolic | Type 1 diabetes | 1.407 | 4.11E-28 | 2192 / 54196 | EA |
| | | | | | | | 250.11 | endocrine/metabolic | Type 1 diabetes with ketoacidosis | 2.052 | 5.76E-18 | 300 / 54196 | EA |
| | | | | | | | 557.1 | digestive | Celiac disease | 1.873 | 1.18E-17 | 382 / 48299 | EA |
| | | | | | | | 250.13 | endocrine/metabolic | Type 1 diabetes with ophthalmic manifestations | 1.883 | 2.52E-13 | 273 / 54196 | EA |
| | | | | | | | 250.12 | endocrine/metabolic | Type 1 diabetes with renal manifestations | 1.795 | 1.24E-12 | 300 / 54196 | EA |
| | | | | | | | 250.3 | endocrine/metabolic | Insulin pump user | 1.204 | 3.04E-12 | 3108 / 54196 | EA |
| | | | | | | | 250 | endocrine/metabolic | Diabetes mellitus | 1.101 | 1.88E-09 | 10334 / 54196 | EA |
| | | | | | | | 250.14 | endocrine/metabolic | Type 1 diabetes with neurological manifestations | 1.437 | 1.24E-08 | 505 / 54196 | EA |
| | | | | | | | 250.7 | endocrine/metabolic | Diabetic retinopathy | 1.294 | 2.65E-07 | 827 / 63608 | EA |
| | | | | | | | 709 | dermatologic | Diffuse diseases of connective tissue | 1.226 | 1.16E-06 | 1202 / 59384 | EA |
| | | | | | | | 250.2 | endocrine/metabolic | Type 2 diabetes | 1.079 | 4.01E-06 | 9876 / 54196 | EA |
| | | | | | | | 709.2 | dermatologic | Sicca syndrome | 1.371 | 1.02E-05 | 402 / 59384 | EA |
| | | | | | | | 244.5 | endocrine/metabolic | Congenital hypothyroidism | 1.833 | 1.99E-05 | 102 / 56362 | EA |
| | | | | | | | 250.23 | endocrine/metabolic | Type 2 diabetes with ophthalmic manifestations | 1.241 | 2.49E-05 | 799 / 54196 | EA |
| 8 | 73004218 | TERF1 | rs183633026 | A | G | 99.36 | 769 | symptoms | Nonallopathic lesions NEC | 14.514 | 1.06E-05 | 73 / 70207 | EA |
| 8 | 73033303 | TERF1 | rs73687065 | T | C | 85.32 | 523.3 | digestive | Periodontitis (acute or chronic) | 21.070 | 6.63E-06 | 214 / 64696 | EA |
| | | | | | | | 389.2 | sense organs | Conductive hearing loss | 3.463 | 1.85E-05 | 90 / 14180 | AA |
| 10 | 99514276 | NKX2-3 | rs10883359 | A | G | -16.52 | 555 | digestive | Inflammatory bowel disease and other gastroenteritis and colitis | 0.856 | 3.86E-06 | 2479 / 48299 | EA |
| | | | | | | | 555.1 | digestive | Regional enteritis | 0.849 | 7.20E-06 | 2103 / 48299 | EA |
| 10 | 103915847 | OBFC1 | rs112163720 | C | T | 37.13 | 634.3 | pregnancy complications | Ectopic pregnancy | 4.927 | 1.01E-05 | 37 / 67408 | EA |
| 14 | 24242592 | TINF2 | rs28372734 | C | G | 112.59 | 379.3 | sense organs | Aphakia and other disorders of lens | 31.987 | 7.73E-06 | 91 / 61188 | EA |
| 14 | 24243052 | TINF2 | rs8016076 | T | C | 83.78 | 385.5 | sense organs | Tympanosclerosis and middle ear disease related to otitis media | 423.472 | 2.52E-06 | 58 / 61381 | EA |
| | | | | | | | 530.3 | digestive | Stricture and stenosis of esophagus | 3.987 | 1.53E-05 | 80 / 10913 | AA |
| | | | | | | | 367.9 | sense organs | Blindness and low vision | 58.593 | 1.98E-05 | 452 / 64773 | EA |
| | | | | | | | 386.1 | sense organs | Meniere's disease | 170.739 | 2.81E-05 | 213 / 60084 | EA |
| 14 | 24254544 | TINF2 | rs41293824 | C | A | 83.12 | 747.2 | congenital anomalies | Congenital anomalies of peripheral vascular system | 51.724 | 1.18E-05 | 203 / 65822 | EA |
| | | | | | | | 340 | neurological | Migraine | 11.942 | 1.58E-05 | 2556 / 62491 | EA |
| 15 | 50065546 | ATP8B4 | rs7172615 | G | T | -17.75 | 200 | neoplasms | Myeloproliferative disease | 1.263 | 3.91E-06 | 1006 / 67218 | EA |
| 18 | 650764 | TYMS | rs150119891 | C | T | -98.77 | 270.2 | endocrine/metabolic | Disorders of amino-acid metabolism | 48.425 | 7.90E-06 | 20 / 14081 | AA |
| 20 | 63695521 | RTEL1 | rs181080831 | G | A | 180.51 | 735.22 | musculoskeletal | Claw toe (acquired) | 25.582 | 5.75E-07 | 28 / 61376 | EA |
| | | | | | | | 840.2 | injuries & poisonings | Rotator cuff (capsule) sprain | 74.457 | 1.26E-06 | 68 / 13031 | AA |

**Table S7, Related to Figure 5:** Results of the PheWAS conducted using SAIGE, a method employing generalized linear mixed models and allowing for imbalance between case and control counts, adjusting for genetic relatedness, sex, birth year and the first 4 principal components of ancestry, on approximately 400,000 UK Biobank (UKBB) participants. Results were downloaded from files provided by the University of Michigan PheWeb server (http://pheweb.sph.umich.edu/SAIGE-UKB/about) on 47 available sentinel variants. Results were evaluated at a Bonferroni threshold corrected for the number of phecodes available for each variant (N=1,403).

| | | | | | | | PheWAS Results | | | | | |
|---|---|---|---|---|---|---|---|---|---|---|---|---|
| **Chr** | **Pos** | **Locus** | **rsNum** | **Ref** | **Alt** | **Trans-ethnic TL Effect size in (bp)** | **Phecode** | **Category** | **Phenotype** | **OR** | **P-Value** | **Number of Samples** |
| 3 | 169487437 | TERC | rs12637184 | G | A | -59.79 | 216 | neoplasms | Benign neoplasm of skin | 0.89 | 6.9E-10 | 7722 / 400618 |
| | | | | | | | 218.1 | neoplasms | Uterine leiomyoma | 0.91 | 7.9E-08 | 10344 / 204369 |
| | | | | | | | 218 | neoplasms | Benign neoplasm of uterus | 0.92 | 7.6E-07 | 10609 / 204369 |
| | | | | | | | 208 | neoplasms | Benign neoplasm of colon | 0.94 | 2.7E-06 | 20204 / 386011 |
| | | | | | | | 622 | genitourinary | Polyp of female genital organs | 0.93 | 9.5E-06 | 10881 / 208551 |
| | | | | | | | 214.1 | neoplasms | Lipoma of skin and subcutaneous tissue | 0.90 | 1.1E-05 | 4611 / 401613 |
| | | | | | | | 214 | neoplasms | Lipoma | 0.91 | 1.8E-05 | 6271 / 401613 |
| 4 | 9930219 | SLC2A2 | rs4235345 | G | A | 16.78 | 274.1 | endocrine/metabolic | Gout | 0.67 | 3.0E-37 | 3195 / 405198 |
| | | | | | | | 274 | endocrine/metabolic | Gout and other crystal arthropathies | 0.71 | 4.4E-33 | 3763 / 405198 |
| 5 | 1285974 | TERT | rs7705526 | C | A | 30.04 | 200 | neoplasms | Myeloproliferative disease | 1.43 | 2.6E-13 | 995 / 404466 |
| | | | | | | | 218.1 | neoplasms | Uterine leiomyoma | 1.10 | 2.7E-09 | 10344 / 204369 |
| | | | | | | | 218 | neoplasms | Benign neoplasm of uterus | 1.10 | 3.2E-09 | 10609 / 204369 |
| | | | | | | | 191 | neoplasms | Manlignant and unknown neoplasms of brain and nervous system | 1.41 | 1.3E-08 | 655 / 407239 |
| | | | | | | | 172.2 | neoplasms | Other non-epithelial cancer of skin | 1.09 | 2.4E-08 | 11149 / 395071 |
| | | | | | | | 191.1 | neoplasms | Cancer of brain and nervous system | 1.45 | 2.4E-08 | 531 / 407239 |
| | | | | | | | 172 | neoplasms | Skin cancer | 1.08 | 3.2E-08 | 13752 / 395071 |
| | | | | | | | 191.11 | neoplasms | Cancer of brain | 1.47 | 3.6E-08 | 486 / 407239 |
| | | | | | | | 220 | neoplasms | Benign neoplasm of ovary | 1.24 | 5.3E-08 | 1481 / 192900 |
| | | | | | | | 702.2 | dermatologic | Seborrheic keratosis | 0.86 | 1.4E-07 | 3092 / 403439 |
| | | | | | | | 200.1 | neoplasms | Polycythemia vera | 1.41 | 1.2E-05 | 390 / 401145 |
| 5 | 1287194 | TERT | rs2853677 | G | A | -23.89 | 172.2 | neoplasms | Other non-epithelial cancer of skin | 0.90 | 5.4E-15 | 11149 / 395071 |
| | | | | | | | 172 | neoplasms | Skin cancer | 0.92 | 1.9E-11 | 13752 / 395071 |
| | | | | | | | 218 | neoplasms | Benign neoplasm of uterus | 0.92 | 2.5E-08 | 10609 / 204369 |
| | | | | | | | 702.2 | dermatologic | Seborrheic keratosis | 1.15 | 2.9E-08 | 3092 / 403439 |
| | | | | | | | 218.1 | neoplasms | Uterine leiomyoma | 0.92 | 4.7E-08 | 10344 / 204369 |
| | | | | | | | 200 | neoplasms | Myeloproliferative disease | 0.79 | 1.9E-07 | 995 / 404466 |
| | | | | | | | 189.2 | neoplasms | Cancer of bladder | 0.87 | 3.4E-06 | 2427 / 404796 |
| | | | | | | | 189.21 | neoplasms | Malignant neoplasm of bladder | 0.87 | 7.9E-06 | 2146 / 404796 |
| | | | | | | | 702 | dermatologic | Degenerative skin conditions and other dermatoses | 1.09 | 1.8E-05 | 5522 / 398746 |
| 5 | 1292958 | TERT | rs114616103 | C | T | -57.21 | 208 | neoplasms | Benign neoplasm of colon | 0.88 | 2.1E-05 | 20204 / 386011 |
| 6 | 31783208 | HSPA1A | rs1008438 | A | C | -20.33 | 557.1 | digestive | Celiac disease | 2.80 | 7.1E-178 | 1855 / 334783 |
| | | | | | | | 557 | digestive | Intestinal malabsorption (non-celiac) | 2.54 | 1.7E-167 | 2103 / 334783 |
| | | | | | | | 250.1 | endocrine/metabolic | Type 1 diabetes | 1.37 | 2.6E-27 | 2660 / 388756 |
| | | | | | | | 242 | endocrine/metabolic | Thyrotoxicosis with or without goiter | 1.25 | 4.6E-11 | 1860 / 391429 |
| | | | | | | | 695.7 | dermatologic | Prurigo and Lichen | 1.35 | 1.2E-08 | 779 / 402672 |
| | | | | | | | 695 | dermatologic | Erythematous conditions | 1.18 | 1.3E-08 | 2420 / 402672 |
| | | | | | | | 244 | endocrine/metabolic | Hypothyroidism | 1.07 | 3.5E-08 | 14871 / 391429 |
| | | | | | | | 454 | circulatory system | Varicose veins | 0.93 | 6.7E-08 | 12172 / 369592 |
| | | | | | | | 242.1 | endocrine/metabolic | Graves' disease | 1.44 | 9.0E-08 | 457 / 391429 |
| | | | | | | | 454.1 | circulatory system | Varicose veins of lower extremity | 0.93 | 1.1E-07 | 11697 / 369592 |
| | | | | | | | 580.12 | genitourinary | Non-proliferative glomerulonephritis | 1.93 | 2.2E-07 | 136 / 397602 |
| | | | | | | | 250.11 | endocrine/metabolic | Type 1 diabetes with ketoacidosis | 1.64 | 2.3E-07 | 237 / 388756 |
| | | | | | | | 244.4 | endocrine/metabolic | Hypothyroidism NOS | 1.07 | 3.2E-07 | 14171 / 391429 |
| | | | | | | | 250 | endocrine/metabolic | Diabetes mellitus | 1.06 | 4.4E-07 | 20203 / 388756 |
| | | | | | | | 250.7 | endocrine/metabolic | Diabetic retinopathy | 1.22 | 4.6E-07 | 1339 / 396859 |
| | | | | | | | 366 | sense organs | Cataract | 1.06 | 7.6E-07 | 20352 / 388609 |
| | | | | | | | 250.23 | endocrine/metabolic | Type 2 diabetes with ophthalmic manifestations | 1.21 | 1.9E-06 | 1298 / 388756 |
| | | | | | | | 709.2 | dermatologic | Sicca syndrome | 1.35 | 4.3E-06 | 513 / 399404 |
| | | | | | | | 250.12 | endocrine/metabolic | Type 1 diabetes with renal manifestations | 2.16 | 6.7E-06 | 76 / 388756 |
| | | | | | | | 251 | endocrine/metabolic | Other disorders of pancreatic internal secretion | 1.24 | 7.3E-06 | 943 / 405386 |
| | | | | | | | 251.1 | endocrine/metabolic | Hypoglycemia | 1.24 | 7.4E-06 | 939 / 386319 |
| | | | | | | | 285 | hematopoietic | Other anemias | 1.06 | 8.9E-06 | 12256 / 390026 |
| | | | | | | | 709 | dermatologic | Diffuse diseases of connective tissue | 1.11 | 1.4E-05 | 3463 / 399404 |
| | | | | | | | 580.2 | genitourinary | Nephrotic syndrome without mention of glomerulonephritis | 1.34 | 1.6E-05 | 469 / 397602 |
| | | | | | | | 371.3 | sense organs | Inflammation of eyelids | 0.88 | 3.3E-05 | 2396 / 399306 |
| 10 | 101274033 | NKX2-3 | rs10883359 | A | G | -16.52 | 555.2 | digestive | Ulcerative colitis | 0.89 | 1.4E-05 | 3195 / 334783 |
| | | | | | | | 555 | digestive | Inflammatory bowel disease and other gastroenteritis and colitis | 0.90 | 2.3E-05 | 4528 / 334783 |
| 10 | 105676465 | OBFC1 | rs9420907 | C | A | -49.22 | 427.2 | circulatory system | Atrial fibrillation and flutter | 0.89 | 1.1E-09 | 14820 / 380919 |
| | | | | | | | 218 | neoplasms | Benign neoplasm of uterus | 0.89 | 3.7E-08 | 10609 / 204369 |
| | | | | | | | 172.11 | neoplasms | Melanomas of skin | 0.80 | 4.5E-08 | 2691 / 395071 |
| | | | | | | | 172.1 | neoplasms | Melanomas of skin, dx or hx | 0.80 | 4.5E-08 | 2691 / 395071 |
| | | | | | | | 214.1 | neoplasms | Lipoma of skin and subcutaneous tissue | 0.84 | 5.2E-08 | 4611 / 401613 |
| | | | | | | | 218.1 | neoplasms | Uterine leiomyoma | 0.89 | 6.9E-08 | 10344 / 204369 |
| | | | | | | | 214 | neoplasms | Lipoma | 0.87 | 1.2E-07 | 6271 / 401613 |
| | | | | | | | 427 | circulatory system | Cardiac dysrhythmias | 0.93 | 6.1E-07 | 24681 / 380919 |
| | | | | | | | 216 | neoplasms | Benign neoplasm of skin | 0.89 | 4.5E-06 | 7722 / 400618 |
| | | | | | | | 704 | dermatologic | Diseases of hair and hair follicles | 0.88 | 2.2E-05 | 5344 / 402357 |
| | | | | | | | 706 | dermatologic | Diseases of sebaceous glands | 0.91 | 2.5E-05 | 8948 / 399255 |
| | | | | | | | 172 | neoplasms | Skin cancer | 0.92 | 2.6E-05 | 13752 / 395071 |
| | | | | | | | 706.2 | dermatologic | Sebaceous cyst | 0.91 | 3.6E-05 | 8876 / 399255 |
| 14 | 73426290 | DCAF4 | rs2572 | C | T | 27.97 | 401 | circulatory system | Hypertension | 1.05 | 2.1E-05 | 77977 / 330366 |
| | | | | | | | 401.1 | circulatory system | Essential hypertension | 1.05 | 0.0000238 | 77723 / 330366 |
| 20 | 62309554 | RTEL1 | rs41309367 | C | T | -34.10 | 555 | digestive | Inflammatory bowel disease and other gastroenteritis and colitis | 1.11 | 0.00000878 | 4528 / 334783 |
| 20 | 62326874 | RTEL1 | rs181080831 | G | A | 180.51 | 165.1 | neoplasms | Cancer of bronchus; lung | 6.75 | 0.00000512 | 2101 / 406226 |
| | | | | | | | 165 | neoplasms | Cancer within the respiratory system | 4.85 | 0.0000119 | 2700 / 406226 |
| | | | | | | | 214 | neoplasms | Lipoma | 2.50 | 0.0000243 | 6271 / 401613 |

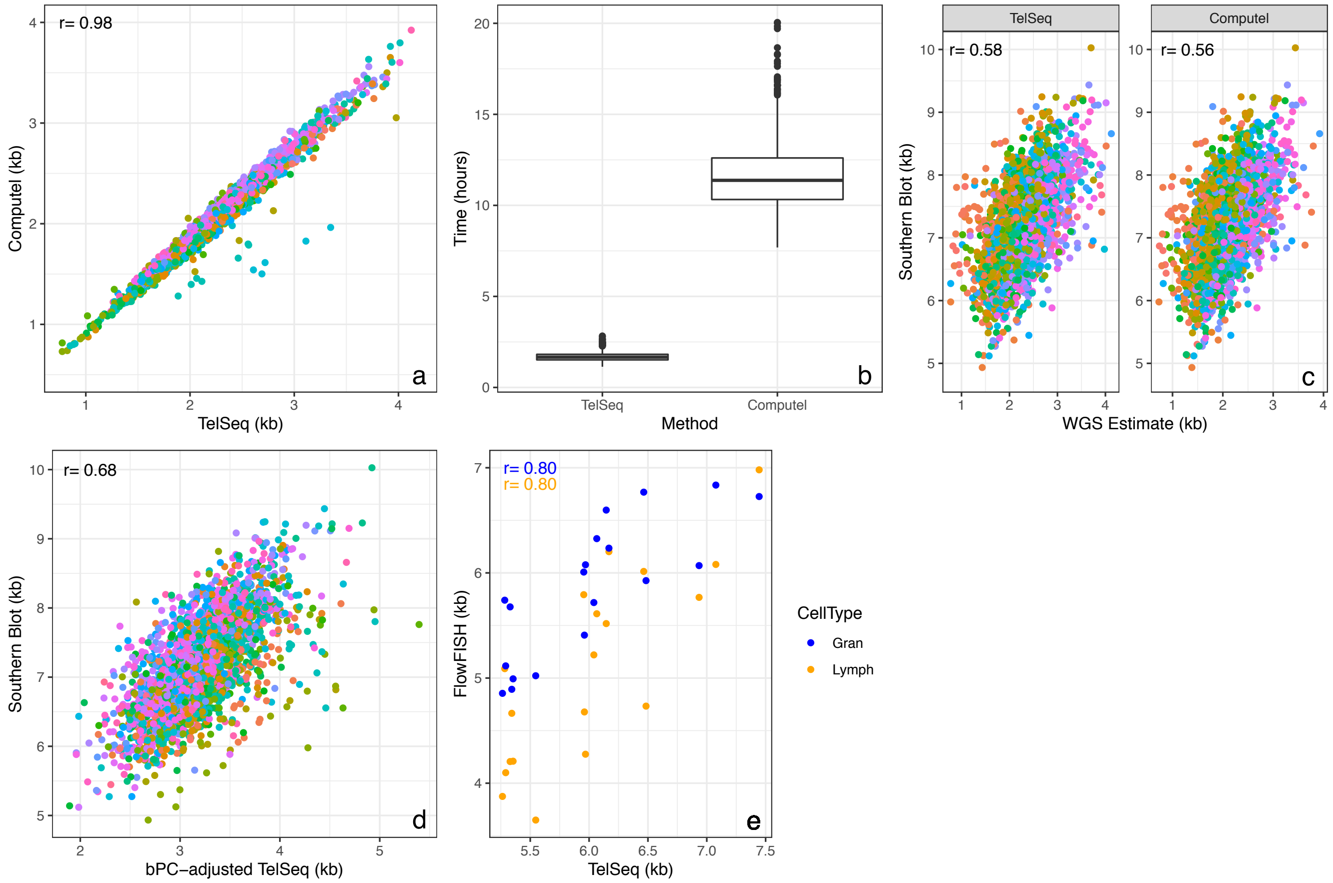


**Figure S1, Related to Estimating telomere length for whole-genome sequencing (WGS) samples and Batch adjustment to correct for technical confounders, Materials and Methods:** For 2,389 samples from the Jackson Heart Study (JHS), **[a]** scatter plot with Pearson correlation between TelSeq and Computel length estimates; **[b]** Comparison of computational times for TelSeq and Computel; **[c]** scatter plots with Pearson correlations between TelSeq (left) and Computel (right) and Southern blot TL estimates; **[d]** scatter plot with Pearson correlation between TelSeq and Southern blot TL estimates after adjustment for the final set of 200 batch principal components (bPCs) used in our full analysis. Colors indicate on which plate samples were shipped to the sequencing center, in Panels A, C and D; and **[e]** scatter plot with Pearson correlation between bPC-adjusted TelSeq and flowFISH data on 19 samples from the GeneSTAR study.

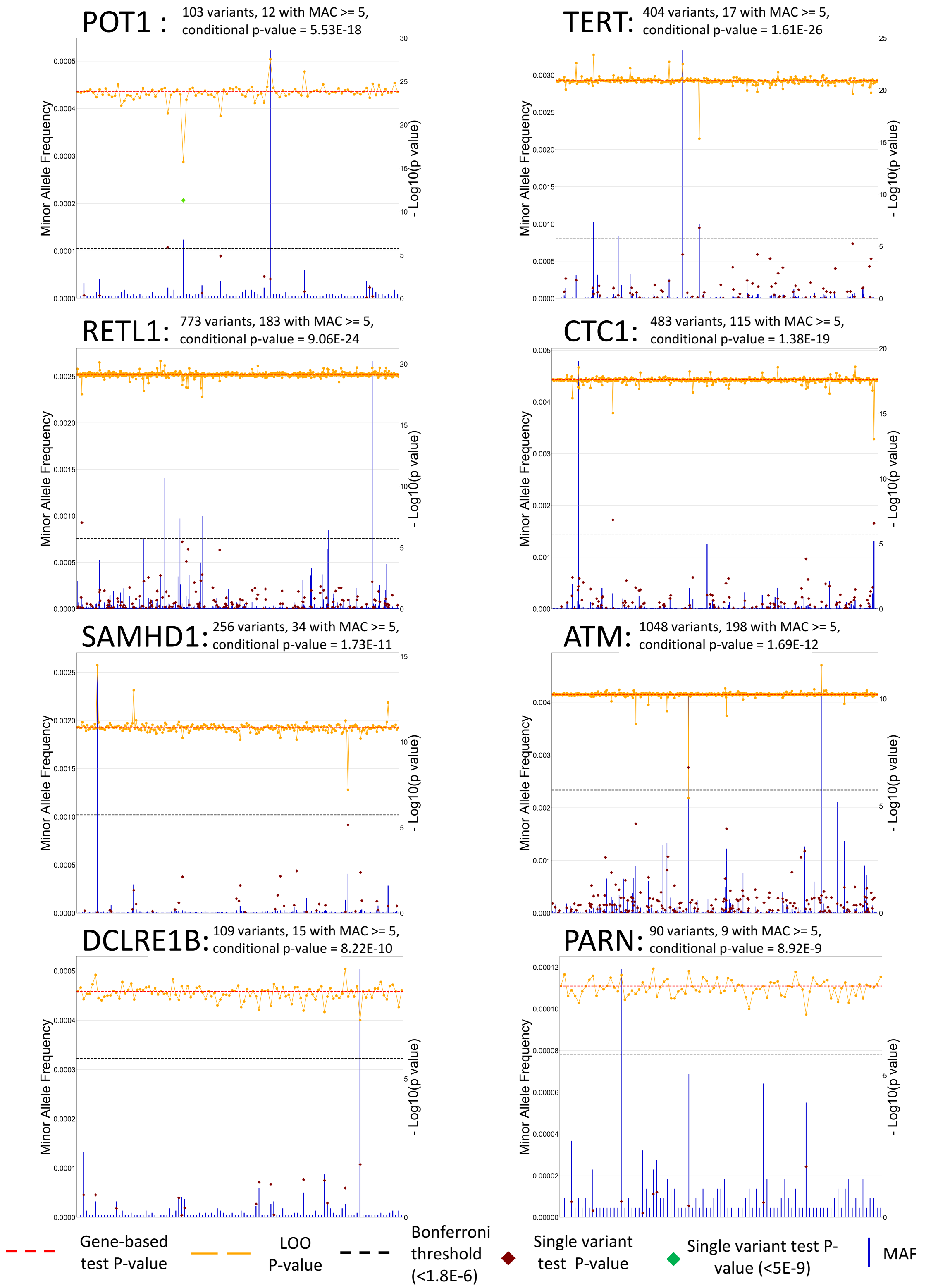


**Figure S2, Related to Gene-based coding variant tests - Tests for association, Materials and Methods**: Eight genes were identified as passing the Bonferroni threshold based on number of genes tested (p-value < 0.05/27,558 = $1.8\times10^{-6}$). For each gene, a leave-one-out analysis was performed iterating the SMMAT test and leaving one variant out at a time. The plots show the change in SMMAT p-value for each variant (orange line with marker) relative to the variant's allele frequency (blue bar), the overall gene-based test including all variants (dotted red line) and the single variant results for all variants with an MAC≥5 that were included in single variant tests for association (brown and green diamonds). For each gene, the number of rare and deleterious variants included in SMMAT is indicated. For any variant with a MAC≥5, a single variant test was also performed as part of the primary analysis. The count of these variants is indicated. In addition the SMMAT p-value for these genes when conditioning on the 59 sentinel variants is also given.

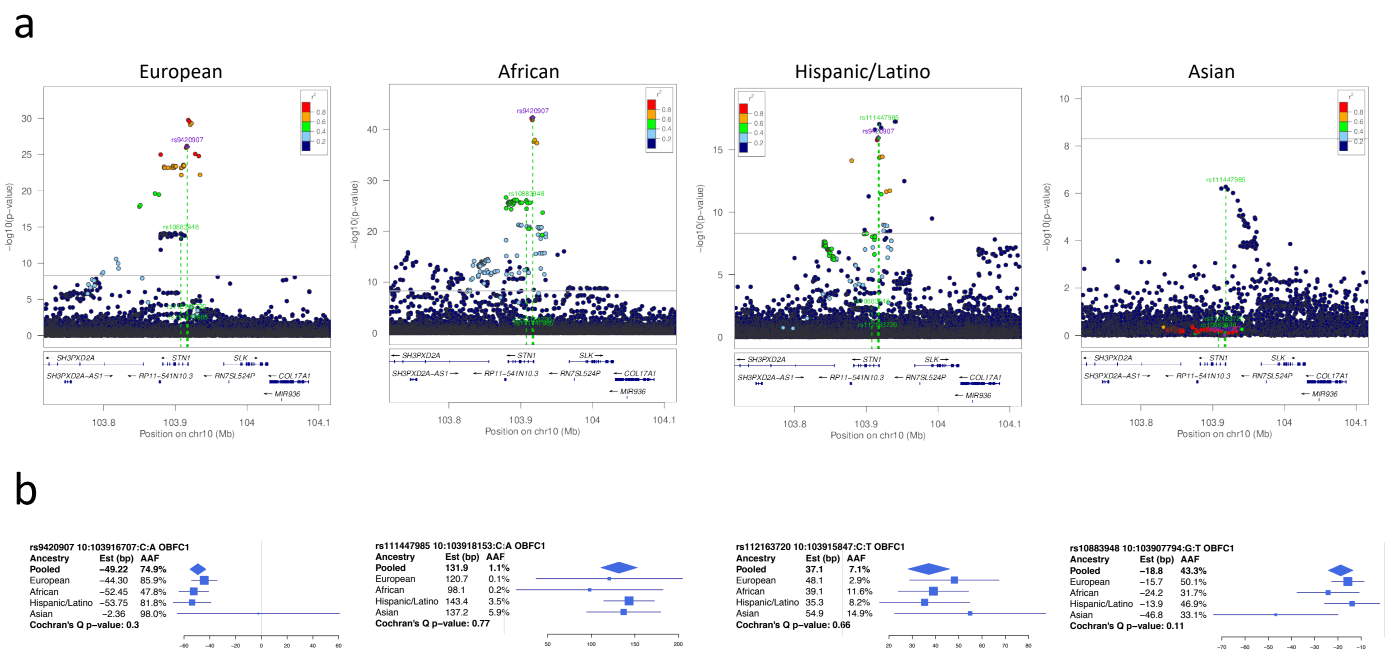


**Figure S3, Related to Figure 4: [a]** LocusZoom plots for four population groups for the *OBFC1* locus. Linkage disequilibrium (LD) was calculated from the set of samples used in the analysis with respect to the peak variant in the pooled trans-ethnic primary analysis, thereby reflecting LD patterns specific to the TOPMed samples. For each figure, the peak sentinel variant from the pooled trans-ethnic analysis is indexed and labeled in purple, and all independent variants identified through the iterative conditional approach are labeled in green and highlighted with green dotted lines. **[b]** Forest plots displaying effect sizes and standard errors, as well as minor allele frequencies, by population group for the four sentinel variants in *OBFC1*.

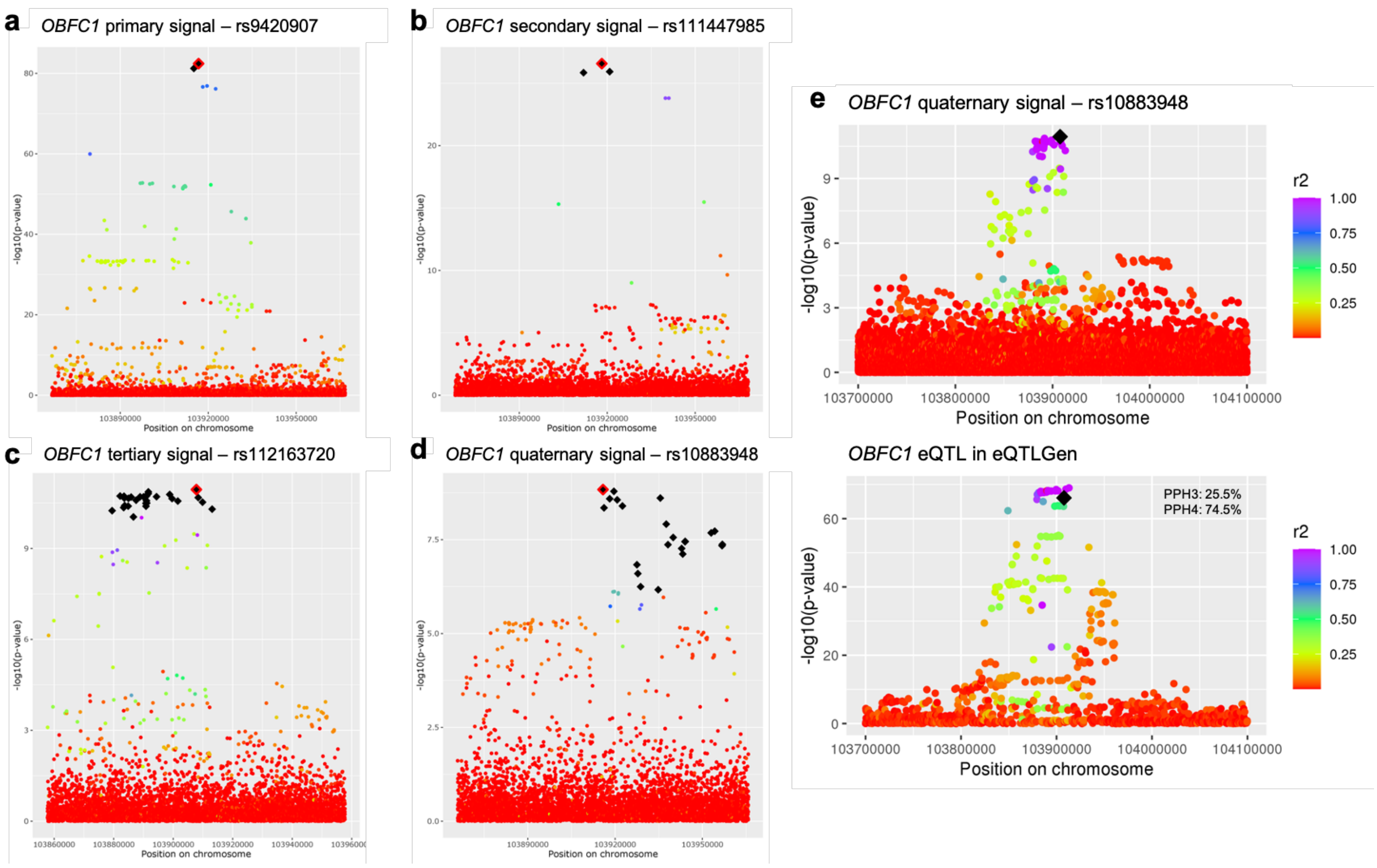


**Figure S4, Related to Figure 4:** Credible set analysis and colocalization analysis in eQTLGen. **[a-d]** Manhattan plots for each *OBFC1* signal are shown where p-values were taken from the appropriate conditional analysis output and LD was calculated with respect to the sentinel variant. Credible set variants are indicated with black diamonds; the sentinel variant is indicated as a black diamond with red outline. **[e]** Manhattan plot for colocalization analysis of the *OBFC1* quaternary signal with an *OBFC1* eQTL in eQTLGen. PPH3 and PPH4 for colocalization are indicated in the top right corner of the eQTL plot. LD was calculated with respect to the sentinel, indicated with a black diamond on each graph, and with pooled trans-ethnic analysis samples.

### Supplementary Information:
### Novel genetic determinants of telomere length from a trans-ethnic analysis of 109,122 whole genome sequences in TOPMed

**Study Acknowledgements**

**Generation of TOPMed whole genome sequencing data by study:**
Whole genome sequencing (WGS) for the Trans-Omics in Precision Medicine (TOPMed) program was supported by the National Heart, Lung and Blood Institute (NHLBI). WGS for NHLBI TOPMed: AFLMU (phs001543) was performed at Broad Genomics (3UM1HG008895-01S2; HHSN268201500014C); WGS for NHLBI TOPMed: Amish (phs000956) was performed at Broad Genomics (3R01HL121007-01S1); WGS for NHLBI TOPMed: ARIC (phs001211) was performed at Baylor (3U54HG003273-12S2 / HHSN268201500015C,3R01HL092577-06S1), Broad Genomics (3U54HG003273-12S2 / HHSN268201500015C,3R01HL092577-06S1); WGS for NHLBI TOPMed: BioMe (phs001644) was performed at MGI (HHSN268201600037I,HHSN268201600033I,3UM1HG008853-01S2), Baylor (HHSN268201600037I,HHSN268201600033I,3UM1HG008853-01S2); WGS for NHLBI TOPMed: CAMP (phs001726) was performed at NWGC (HHSN268201600032I); WGS for NHLBI TOPMed: CARDIA (phs001612) was performed at Baylor (HHSN268201600033I); WGS for NHLBI TOPMed: CARE_BADGER (phs001728) was performed at NWGC (HHSN268201600032I); WGS for NHLBI TOPMed: CARE_CLIC (phs001729) was performed at NWGC (HHSN268201600032I); WGS for NHLBI TOPMed: CARE_PACT (phs001730) was performed at NWGC (HHSN268201600032I); WGS for NHLBI TOPMed: CARE_TREXA (phs001732) was performed at NWGC (HHSN268201600032I); WGS for NHLBI TOPMed: CFS (phs000954) was performed at NWGC (HHSN268201600032I,3R01HL098433-05S1); WGS for NHLBI TOPMed: ChildrensHS_GAP (phs001602) was performed at NWGC (HHSN268201600032I); WGS for NHLBI TOPMed: ChildrensHS_IGERA (phs001603) was performed at NWGC (HHSN268201600032I); WGS for NHLBI TOPMed: ChildrensHS_MetaAir (phs001604) was performed at NWGC (HHSN268201600032I); WGS for NHLBI TOPMed: CHIRAH (phs001605) was performed at NWGC (HHSN268201600032I); WGS for NHLBI TOPMed: CHS (phs001368) was performed at Baylor (HHSN268201600033I,3U54HG003273-12S2 / HHSN268201500015C); WGS for NHLBI TOPMed: COPDGene (phs000951) was performed at NWGC (3R01HL089856-08S1,HHSN268201500014C), Broad Genomics (3R01HL089856-08S1,HHSN268201500014C); WGS for NHLBI TOPMed: CRA (phs000988) was performed at NWGC (3R37HL066289-13S1,HHSN268201600032I); WGS for NHLBI TOPMed: DHS (phs001412) was performed at Broad Genomics (HHSN268201500014C); WGS for NHLBI TOPMed: ECLIPSE (phs001472) was performed at MGI (HHSN268201600037I); WGS for NHLBI TOPMed: EOCOPD (phs000946) was performed at NWGC (3R01HL089856-08S1); WGS for NHLBI TOPMed: FHS (phs000974) was performed at Broad Genomics (3U54HG003067-12S2,3R01HL092577-06S1); WGS for NHLBI TOPMed: GALAI (phs001542) was performed at NWGC (HHSN268201600032I); WGS for NHLBI TOPMed: GALAII (phs000920) was performed at NYGC (3R01HL117004-02S3,HHSN268201600032I), NWGC (3R01HL117004-02S3,HHSN268201600032I), NYGC (UM1 HG008901); WGS for NHLBI TOPMed: GeneSTAR (phs001218) was performed at Psomagen (3R01HL112064-04S1,R01HL112064,HHSN268201500014C), Illumina (3R01HL112064-04S1,R01HL112064,HHSN268201500014C), Broad Genomics (3R01HL112064-04S1,R01HL112064,HHSN268201500014C); WGS for NHLBI TOPMed: GENOA (phs001345) was performed at NWGC (3R01HL055673-18S1,HHSN268201500014C), Broad Genomics (3R01HL055673-18S1,HHSN268201500014C); WGS for NHLBI TOPMed: GenSalt (phs001217) was performed at Baylor (HHSN268201500015C); WGS for NHLBI TOPMed: GOLDN (phs001359) was performed at NWGC (3R01HL104135-04S1); WGS for NHLBI TOPMed: HCHS_SOL (phs001395) was

performed at Baylor (HHSN268201600033I); WGS for NHLBI TOPMed: HVH (phs000993) was performed at Broad Genomics (3R01HL092577-06S1,3U54HG003273-12S2 / HHSN268201500015C), Baylor (3R01HL092577-06S1,3U54HG003273-12S2 / HHSN268201500015C); WGS for NHLBI TOPMed: HyperGEN (phs001293) was performed at NWGC (3R01HL055673-18S1); WGS for NHLBI TOPMed: IPF (phs001607) was performed at MGI (HHSN268201600037I); WGS for NHLBI TOPMed: JHS (phs000964) was performed at NWGC (HHSN268201100037C); WGS for NHLBI TOPMed: LTRC (phs001662) was performed at Broad Genomics (HHSN268201600034I); WGS for NHLBI TOPMed: Mayo_VTE (phs001402) was performed at Baylor (3U54HG003273-12S2 / HHSN268201500015C); WGS for NHLBI TOPMed: MESA (phs001416) was performed at Broad Genomics (3U54HG003067-13S1,HHSN268201500014C); WGS for NHLBI TOPMed: MLOF (phs001515) was performed at Baylor (HHSN268201600033I,HHSN268201500016C), NYGC (HHSN268201600033I,HHSN268201500016C); WGS for NHLBI TOPMed: OMG_SCD (phs001608) was performed at Baylor (HHSN268201500015C); WGS for NHLBI TOPMed: PCGC_CHD (phs001735) was performed at Broad Genomics (HHSN268201600034I); WGS for NHLBI TOPMed: PharmHU (phs001466) was performed at Baylor (HHSN268201500015C); WGS for NHLBI TOPMed: PIMA (phs001727) was performed at NWGC (HHSN268201600032I); WGS for NHLBI TOPMed: PUSH_SCD (phs001682) was performed at Baylor (HHSN268201500015C); WGS for NHLBI TOPMed: REDS-III_Brazil (phs001468) was performed at Baylor (HHSN268201500015C); WGS for NHLBI TOPMed: SAFS (phs001215) was performed at Illumina (R01HL113322,3R01HL113323-03S1); WGS for NHLBI TOPMed: SAGE (phs000921) was performed at NYGC (3R01HL117004-02S3,HHSN268201600032I), NWGC (3R01HL117004-02S3,HHSN268201600032I); WGS for NHLBI TOPMed: SAPPHIRE_asthma (phs001467) was performed at NWGC (HHSN268201600032I); WGS for NHLBI TOPMed: SARP (phs001446) was performed at NYGC (HHSN268201500016C); WGS for NHLBI TOPMed: SAS (phs000972) was performed at NWGC (HHSN268201100037C,HHSN268201500016C), NYGC (HHSN268201100037C,HHSN268201500016C); WGS for NHLBI TOPMed: THRV (phs001387) was performed at Baylor (3R01HL111249-04S1 / HHSN26820150015C); WGS for NHLBI TOPMed: VAFAR (phs000997) was performed at Broad Genomics (3U54HG003067-12S2 / 3U54HG003067-13S1; 3UM1HG008895-01S2; 3UM1HG008895-01S2,3R01HL092577-06S1); WGS for NHLBI TOPMed: VU_AF (phs001032) was performed at Broad Genomics (3R01HL092577-06S1); WGS for NHLBI TOPMed: walk_PHaSST (phs001514) was performed at Baylor (HHSN268201500015C); WGS for NHLBI TOPMed: WGHS (phs001040) was performed at Broad Genomics (3R01HL092577-06S1); WGS for NHLBI TOPMed: WHI (phs001237) was performed at Broad Genomics (HHSN268201500014C). Core support including centralized genomic read mapping and genotype calling, along with variant quality metrics and filtering were provided by the TOPMed Informatics Research Center (3R01HL-117626-02S1; contract HHSN268201800002I). Core support including phenotype harmonization, data management, sample-identity QC, and general program coordination were provided by the TOPMed Data Coordinating Center (R01HL-120393; U01HL-120393; contract HHSN268201800001I). We gratefully acknowledge the studies and participants who provided biological samples and data for TOPMed. NYGC = New York Genome Center; Broad Genomics = Broad Institute Genomics Platform; NWGC = University of Washington Northwest Genomics Center; Illumina = Illumina Genomic Services; Psomagen = Psomagen Corp.; Baylor = Baylor Human Genome Sequencing Center; MGI = McDonnell Genome Institute

**NHLBI TOPMed: Atrial Fibrillation Biobank LMU (AFLMU) in the context of the Arrhythmia-Biobank-LMU**

AFLMU is a repository of AF patients recruited in the context of the German Competence Network for Atrial Fibrillation (AFNET) and at the Department of Medicine I of the University Hospital Munich. In this context, DNA samples were preferentially sampled if the patient developed AF before the age of 60 years. Cases were selected if the diagnosis of atrial fibrillation was made on an electrocardiogram analyzed by a trained physician. Patients with signs of moderate to severe heart failure, moderate to severe valve disease or with hyperthyroidism were excluded from the study. All participants provided written informed consent. AFLMU was approved by the Ethics Committee at the Ludwig-Maximilian's University.

**NHLBI TOPMed: Genetics of Cardiometabolic Health in the Amish (Amish)**
The Amish studies upon which these data are based were supported by NIH grants R01 AG18728, U01 HL072515, R01 HL088119, R01 HL121007, and P30 DK072488. See publication: PMID: 18440328

**NHLBI TOPMed: Atherosclerosis Risk in Communities (ARIC)**
The Atherosclerosis Risk in Communities study has been funded in whole or in part with Federal funds from the National Heart, Lung, and Blood Institute, National Institutes of Health, Department of Health and Human Services (contract numbers HHSN268201700001I, HHSN268201700002I, HHSN268201700003I, HHSN268201700004I and HHSN268201700005I). The authors thank the staff and participants of the ARIC study for their important contributions.

**NHLBI TOPMed: The Genetics and Epidemiology of Asthma in Barbados (BAGS)**
We gratefully acknowledge the contributions of Pissamai and Trevor Maul, Paul Levett, Anselm Hennis, P. Michele Lashley, Raana Naidu, Malcolm Howitt and Timothy Roach, and the numerous health care providers, and community clinics and co-investigators who assisted in the phenotyping and collection of DNA samples, and the families and patients for generously donating DNA samples to the Barbados Asthma Genetics Study (BAGS). The Genetics and Epidemiology of Asthma in Barbados is supported by National Institutes of Health (NIH) National Heart, Lung, Blood Institute TOPMed (R01 HL104608-S1) and: R01 AI20059, K23 HL076322, R01HL087699, and RC2 HL101651. For the specific cohort descriptions and descriptions regarding the collection of phenotype data can be found at: https://www.nhlbiwgs.org/group/bags-asthma. The authors wish to give special recognition to the individual study participants who provided biological samples and or data, without their support in research none of this would be possible.

**NHLBI TOPMed: BioMe Biobank at Mount Sinai (BioMe)**
The Mount Sinai BioMe Biobank has been supported by The Andrea and Charles Bronfman Philanthropies and in part by Federal funds from the NHLBI and NHGRI (U01HG00638001; U01HG007417; X01HL134588). We thank all participants in the Mount Sinai Biobank. We also thank all our recruiters who have assisted and continue to assist in data collection and management and are grateful for the computational resources and staff expertise provided by Scientific Computing at the Icahn School of Medicine at Mount Sinai.

**NHLBI TOPMed: CAMP**
We thank the clinical centers and the Data Coordinating Center of the Childhood Asthma Management Program (CAMP) as well as all of the study participants at the 8 clinical sites. The CAMP study was supported by NHLBI P01 HL132825.

**NHLBI TOPMed: Coronary Artery Risk Development in Young Adults Study (CARDIA)**
The Coronary Artery Risk Development in Young Adults Study (CARDIA) is conducted and supported by the National Heart, Lung, and Blood Institute (NHLBI) in collaboration with the University of Alabama at Birmingham (HHSN268201800005I & HHSN268201800007I), Northwestern University (HHSN268201800003I), University of Minnesota (HHSN268201800006I), and Kaiser Foundation Research Institute (HHSN268201800004I). CARDIA was also partially supported by the Intramural Research Program of the National Institute on Aging (NIA) and an intra-agency agreement between NIA and NHLBI (AG0005).

**NHLBI TOPMed: CARE**_BADGER
This research was supported by grants from the National Heart, Lung, and Blood Institute (NHLBI), ((5U10HL064287, 5U10HL064288, 5U10HL064295, 5U10HL064307, 5U10HL064305, 5U10HL064313, and HL080083)

**NHLBI TOPMed: CARE_CLIC**
This research was supported by grants from the National Heart, Lung, and Blood Institute (NHLBI), ((5U10HL064287, 5U10HL064288, 5U10HL064295, 5U10HL064307, 5U10HL064305, 5U10HL064313, and HL080083)

**NHLBI TOPMed: CARE_PACT**
This research was supported by grants from the National Heart, Lung, and Blood Institute (NHLBI), ((5U10HL064287, 5U10HL064288, 5U10HL064295, 5U10HL064307, 5U10HL064305, 5U10HL064313, and HL080083)

**NHLBI TOPMed: CARE_TREXA**
This research was supported by grants from the National Heart, Lung, and Blood Institute (NHLBI), ((5U10HL064287, 5U10HL064288, 5U10HL064295, 5U10HL064307, 5U10HL064305, 5U10HL064313, and HL080083)

**NHLBI TOPMed: The Cleveland Family Study (CFS)**
The Cleveland Family Study has been supported in part by National Institutes of Health grants [R01-HL046380, KL2-RR024990, R35-HL135818, and R01-HL113338].

**NHLBI TOPMed: Children's Health Study: Integrative Genetic Approaches to Gene-Air Pollution Interactions in Asthma (ChildrensHS_GAP)**
The Integrative Genetic Approaches to Gene-Air Pollution Interactions in Asthma (GAP) study was supported by the National Institute of Environmental Health Sciences (NIEHS) grant # R01ES021801. The Children's Health Study (CHS) was supported by the Southern California Environmental Health Sciences Center (grant P30ES007048); National Institute of Environmental Health Sciences (grants 5P01ES011627, ES021801, ES023262, P01ES009581, P01ES011627, P01ES022845, R01 ES016535, R03ES014046, P50 CA180905, R01HL061768, R01HL076647, R01HL087680 and RC2HL101651), the Environmental Protection Agency (grants RD83544101, R826708, RD831861, and R831845), and the Hastings Foundation.

**NHLBI TOPMed: Children's Health Study: Integrative Genomics and Environmental Research of Asthma (ChildrensHS_IGERA)**
The Integrative Genomics and Environmental Research of Asthma (IGERA) Study was supported by the National Heart, Lung and Blood Institute (grant # RC2HL101543 -The Asthma BioRepository for Integrative Genomics Research, PI Gilliland/Raby). The Children's Health Study (CHS) was supported by the Southern California Environmental Health Sciences Center (grant P30ES007048); National Institute of Environmental Health Sciences (grants 5P01ES011627, ES021801, ES023262, P01ES009581, P01ES011627, P01ES022845, R01 ES016535, R03ES014046, P50 CA180905, R01HL061768, R01HL076647, R01HL087680 and RC2HL101651), the Environmental Protection Agency (grants RD83544101, R826708, RD831861, and R831845), and the Hastings Foundation.

**NHLBI TOPMed: Children's Health Study: Effects of Air Pollution on the Development of Obesity in Children (ChildrensHS_MetaAir)**

The Effects of Air Pollution on the Development of Obesity in Children (Meta-AIR) study was supported by the Southern California Children's Environmental Health Center funded by the National Institute of Environmental Health Sciences (NIEHS) (P01ES022845) and the Environmental Protection Agency (EPA) (RD-83544101–0). The Children's Health Study (CHS) was supported by the Southern California Environmental Health Sciences Center (grant P30ES007048); National Institute of Environmental Health Sciences (grants 5P01ES011627, ES021801, ES023262, P01ES009581, P01ES011627, P01ES022845, R01 ES016535, R03ES014046, P50 CA180905, R01HL061768, R01HL076647, R01HL087680 and RC2HL101651), the Environmental Protection Agency (grants RD83544101, R826708, RD831861, and R831845), and the Hastings Foundation.

**NHLBI TOPMed: Genetics Sub-Study of Chicago Initiative to Raise Asthma Health Equity (CHIRAH)**
Support for the Genetics Sub-Study of Chicago Initiative to Raise Asthma Health Equity was provided by NHLBI grant number UO1 HL072496.

**NHLBI TOPMed: Cardiovascular Health Study (CHS)**
This research was supported by contracts HHSN268201200036C, HHSN268200800007C, HHSN268201800001C, N01HC55222, N01HC85079, N01HC85080, N01HC85081, N01HC85082, N01HC85083, N01HC85086, and grants U01HL080295 and U01HL130114 from the National Heart, Lung, and Blood Institute (NHLBI), with additional contribution from the National Institute of Neurological Disorders and Stroke (NINDS). Additional support was provided by R01AG023629 from the National Institute on Aging (NIA). A full list of principal CHS investigators and institutions can be found at CHS-NHLBI.org. The content is solely the responsibility of the authors and does not necessarily represent the official views of the National Institutes of Health.

**NHLBI TOPMed: Genetic Epidemiology of COPD (COPDGene) in the TOPMed Program**
The COPDGene project described was supported by Award Number U01 HL089897 and Award Number U01 HL089856 from the National Heart, Lung, and Blood Institute. The content is solely the responsibility of the authors and does not necessarily represent the official views of the National Heart, Lung, and Blood Institute or the National Institutes of Health. The COPDGene project is also supported by the COPD Foundation through contributions made to an Industry Advisory Board comprised of AstraZeneca, Boehringer Ingelheim, GlaxoSmithKline, Novartis, Pfizer, Siemens and Sunovion. A full listing of COPDGene investigators can be found at: http://www.copdgene.org/directory

**NHLBI TOPMed: The Genetic Epidemiology of Asthma in Costa Rica (CRA)**
This study was supported by NHLBI grants R37 HL066289 and P01 HL132825. We wish to acknowledge the investigators at the Channing Division of Network Medicine at Brigham and Women's Hospital, the investigators at the Hospital Nacional de Niños in San José, Costa Rica and the study subjects and their extended family members who contributed samples and genotypes to the study, and the NIH/NHLBI for its support in making this project possible.

**NHLBI TOPMed: Diabetes Heart Study (DHS)**
This work was supported by R01 HL92301, R01 HL67348, R01 NS058700, R01 AR48797, R01 DK071891, R01 AG058921, the General Clinical Research Center of the Wake Forest University School of Medicine (M01 RR07122, F32 HL085989), the American Diabetes Association, and a pilot grant from the Claude Pepper Older Americans Independence Center of Wake Forest University Health Sciences (P60 AG10484).

**NHLBI TOPMed: ECLIPSE**
The ECLIPSE study (NCT00292552) was sponsored by GlaxoSmithKline. The ECLIPSE investigators included: ECLIPSE Investigators — Bulgaria: Y. Ivanov, Pleven; K. Kostov, Sofia. Canada: J. Bourbeau,

Montreal; M. Fitzgerald, Vancouver, BC; P. Hernandez, Halifax, NS; K. Killian, Hamilton, ON; R. Levy, Vancouver, BC; F. Maltais, Montreal; D. O'Donnell, Kingston, ON. Czech Republic: J. Krepelka, Prague. Denmark: J. Vestbo, Hvidovre. The Netherlands: E. Wouters, Horn-Maastricht. New Zealand: D. Quinn, Wellington. Norway: P. Bakke, Bergen. Slovenia: M. Kosnik, Golnik. Spain: A. Agusti, J. Sauleda, P. de Mallorca. Ukraine: Y. Feschenko, V. Gavrisyuk, L. Yashina, Kiev; N. Monogarova, Donetsk. United Kingdom: P. Calverley, Liverpool; D. Lomas, Cambridge; W. MacNee, Edinburgh; D. Singh, Manchester; J. Wedzicha, London. United States: A. Anzueto, San Antonio, TX; S. Braman, Providence, RI; R. Casaburi, Torrance CA; B. Celli, Boston; G. Giessel, Richmond, VA; M. Gotfried, Phoenix, AZ; G. Greenwald, Rancho Mirage, CA; N. Hanania, Houston; D. Mahler, Lebanon, NH; B. Make, Denver; S. Rennard, Omaha, NE; C. Rochester, New Haven, CT; P. Scanlon, Rochester, MN; D. Schuller, Omaha, NE; F. Sciurba, Pittsburgh; A. Sharafkhaneh, Houston; T. Siler, St. Charles, MO; E. Silverman, Boston; A. Wanner, Miami; R. Wise, Baltimore; R. ZuWallack, Hartford, CT. ECLIPSE Steering Committee: H. Coxson (Canada), C. Crim (GlaxoSmithKline, USA), L. Edwards (GlaxoSmithKline, USA), D. Lomas (UK), W. MacNee (UK), E. Silverman (USA), R. Tal-Singer (Co-chair, GlaxoSmithKline, USA), J. Vestbo (Co-chair, Denmark), J. Yates (GlaxoSmithKline, USA). ECLIPSE Scientific Committee: A. Agusti (Spain), P. Calverley (UK), B. Celli (USA), C. Crim (GlaxoSmithKline, USA), B. Miller (GlaxoSmithKline, USA), W. MacNee (Chair, UK), S. Rennard (USA), R. Tal-Singer (GlaxoSmithKline, USA), E. Wouters (The Netherlands), J. Yates (GlaxoSmithKline, USA).

**NHLBI TOPMed: Boston Early-Onset COPD Study in the TOPMed Program (EOCOPD)**
The Boston Early-Onset COPD Study was supported by R01 HL113264 and U01 HL089856 from the National Heart, Lung, and Blood Institute.

**NHLBI TOPMed: Whole Genome Sequencing and Related Phenotypes in the Framingham Heart Study (FHS)**
The Framingham Heart Study (FHS) acknowledges the support of contracts NO1-HC-25195, HHSN268201500001I, and 75N92019D00031 from the National Heart, Lung and Blood Institute and grant supplement R01 HL092577-06S1 for this research. We also acknowledge the dedication of the FHS study participants without whom this research would not be possible.

**NHLBI TOPMed: Genes-environments and Admixture in Latino Asthmatics (GALA I) Study**
The Genes-environments and Admixture in Latino Americans (GALA I) Study was supported by the National Heart, Lung, and Blood Institute of the National Institute of Health (NIH) grants R01HL117004 and X01HL134589; study enrollment supported by Sandler Center for Basic Research in Asthma and the Sandler Family Foundation, the American Asthma Foundation, the American Lung Association, the NIH grants K23HL04464 and HL07185, the Resource Centers for Minority Aging Research from the National Institute on Aging, RCMAR P30-AG15272, the National Institute of Nursing Research and the National Center on Minority Health and Health Disparities.

**NHLBI TOPMed: Genes-environments and Admixture in Latino Asthmatics (GALA II) Study**
The Genes-environments and Admixture in Latino Americans (GALA II) Study was supported by the National Heart, Lung, and Blood Institute of the National Institute of Health (NIH) grants R01HL117004 and X01HL134589; study enrollment supported by the Sandler Family Foundation, the American Asthma Foundation, the RWJF Amos Medical Faculty Development Program, Harry Wm. and Diana V. Hind Distinguished Professor in Pharmaceutical Sciences II and the National Institute of Environmental Health Sciences grant R01ES015794 .
WGS of part of GALA II was performed by New York Genome Center under The Centers for Common Disease Genomics of the Genome Sequencing Program (GSP) Grant (UM1 HG008901). The GSP Coordinating Center (U24 HG008956) contributed to cross-program scientific initiatives and provided logistical and general study coordination. GSP is funded by the National Human Genome Research Institute, the National Heart, Lung, and Blood Institute, and the National Eye Institute.

The GALA II study collaborators include Shannon Thyne, UCSF; Harold J. Farber, Texas Children's Hospital; Denise Serebrisky, Jacobi Medical Center; Rajesh Kumar, Lurie Children's Hospital of Chicago; Emerita Brigino-Buenaventura, Kaiser Permanente; Michael A. LeNoir, Bay Area Pediatrics; Kelley Meade, UCSF Benioff Children’s Hospital, Oakland; William Rodriguez-Cintron, VA Hospital, Puerto Rico; Pedro C. Avila, Northwestern University; Jose R. Rodriguez-Santana, Centro de Neumologia Pediatrica; Luisa N. Borrell, City University of New York; Adam Davis, UCSF Benioff Children's Hospital, Oakland; Saunak Sen, University of Tennessee and Fred Lurmann, Sonoma Technologies, Inc. The authors acknowledge the families and patients for their participation and thank the numerous health care providers and community clinics for their support and participation in GALA II. In particular, the authors thank study coordinator Sandra Salazar; the recruiters who obtained the data: Duanny Alva, MD, Gaby Ayala-Rodriguez, Lisa Caine, Elizabeth Castellanos, Jaime Colon, Denise DeJesus, Blanca Lopez, Brenda Lopez, MD, Louis Martos, Vivian Medina, Juana Olivo, Mario Peralta, Esther Pomares, MD, Jihan Quraishi, Johanna Rodriguez, Shahdad Saeedi, Dean Soto, Ana Taveras; and the lab researcher Celeste Eng who processed the biospecimens.

**NHLBI TOPMed: GeneSTAR (Genetic Study of Atherosclerosis Risk)**
The Johns Hopkins Genetic Study of Atherosclerosis Risk (GeneSTAR) was supported by grants from the National Institutes of Health through the National Heart, Lung, and Blood Institute (U01HL72518, HL087698, HL112064) and by a grant from the National Center for Research Resources (M01-RR000052) to the Johns Hopkins General Clinical Research Center. We would like to thank the participants and families of GeneSTAR and our dedicated staff for all their sacrifices.

**NHLBI TOPMed: Genetic Epidemiology Network of Arteriopathy (GENOA)**
Support for GENOA was provided by the National Heart, Lung and Blood Institute (HL054457, HL054464, HL054481, HL119443, and HL087660) of the National Institutes of Health.

**NHLBI TOPMed: Genetic Epidemiology Network of Salt Sensitivity (GenSalt)**
The Genetic Epidemiology Network of Salt-Sensitivity (GenSalt) was supported by research grants (U01HL072507, R01HL087263, and R01HL090682) from the National Heart, Lung and Blood Institute, National Institutes of Health, Bethesda, MD.

**NHLBI TOPMed: Genetics of Lipid Lowering Drugs and Diet Network (GOLDN)**
GOLDN biospecimens, baseline phenotype data, and intervention phenotype data were collected with funding from National Heart, Lung and Blood Institute (NHLBI) grant U01 HL072524. Whole-genome sequencing in GOLDN was funded by NHLBI grant R01 HL104135 and supplement R01 HL104135-04S1.

**NHLBI TOPMed: Hispanic Community Health Study/Study of Latinos (HCHS_SOL)**
The Hispanic Community Health Study/Study of Latinos is a collaborative study supported by contracts from the National Heart, Lung, and Blood Institute (NHLBI) to the University of North Carolina (HHSN268201300001I / N01-HC-65233), University of Miami (HHSN268201300004I / N01-HC-65234), Albert Einstein College of Medicine (HHSN268201300002I / N01-HC-65235), University of Illinois at Chicago – HHSN268201300003I / N01-HC-65236 Northwestern Univ), and San Diego State University (HHSN268201300005I / N01-HC-65237). The following Institutes/Centers/Offices have contributed to the HCHS/SOL through a transfer of funds to the NHLBI: National Institute on Minority Health and Health Disparities, National Institute on Deafness and Other Communication Disorders, National Institute of Dental and Craniofacial Research, National Institute of Diabetes and Digestive and Kidney Diseases, National Institute of Neurological Disorders and Stroke, NIH Institution-Office of Dietary Supplements.

**NHLBI TOPMed: Heart and Vascular Health Study (HVH)**

The Heart and Vascular Health Study was supported by grants HL068986, HL085251, HL095080, and HL073410 from the National Heart, Lung, and Blood Institute.

**NHLBI TOPMed: Hypertension Genetic Epidemiology Network (HyperGEN)**
The HyperGEN Study is part of the National Heart, Lung, and Blood Institute (NHLBI) Family Blood Pressure Program; collection of the data represented here was supported by grants U01 HL054472 (MN Lab), U01 HL054473 (DCC), U01 HL054495 (AL FC), and U01 HL054509 (NC FC). The HyperGEN: Genetics of Left Ventricular Hypertrophy Study was supported by NHLBI grant R01 HL055673 with whole-genome sequencing made possible by supplement -18S1.

**NHLBI TOPMed: IPF**
This research was supported by the National Heart, Lung and Blood Institute (R01-HL097163, P01-HL092870, and UH3-HL123442) and the Department of Defense (W81XWH-17-1-0597).

**NHLBI TOPMed: The Jackson Heart Study (JHS)**
The Jackson Heart Study (JHS) is supported and conducted in collaboration with Jackson State University (HHSN268201800013I), Tougaloo College (HHSN268201800014I), the Mississippi State Department of Health (HHSN268201800015I) and the University of Mississippi Medical Center (HHSN268201800010I, HHSN268201800011I and HHSN268201800012I) contracts from the National Heart, Lung, and Blood Institute (NHLBI) and the National Institute on Minority Health and Health Disparities (NIMHD). The authors also wish to thank the staffs and participants of the JHS.

**NHLBI TOPMed: LTRC**
This study utilized biological specimens and data provided by the Lung Tissue Research Consortium (LTRC) supported by the National Heart, Lung, and Blood Institute (NHLBI). The LTRC was sponsored by a contract from the NHLBI: HHSN2682016000021

**NHLBI TOPMed: Mayo Clinic Venous Thromboembolism Study (Mayo_VTE)**
Funded, in part, by grants from the National Institutes of Health, National Heart, Lung and Blood Institute (HL66216 and HL83141). the National Human Genome Research Institute (HG04735, HG06379), and research support provided by Mayo Foundation.

**NHLBI TOPMed: Multi-Ethnic Study of Atherosclerosis (MESA)**
Support for the Multi-Ethnic Study of Atherosclerosis (MESA) projects are conducted and supported by the National Heart, Lung, and Blood Institute (NHLBI) in collaboration with MESA investigators. Support for MESA is provided by contracts 75N92020D00001, HHSN268201500003I, N01-HC-95159, 75N92020D00005, N01-HC-95160, 75N92020D00002, N01-HC-95161, 75N92020D00003, N01-HC-95162, 75N92020D00006, N01-HC-95163, 75N92020D00004, N01-HC-95164, 75N92020D00007, N01-HC-95165, N01-HC-95166, N01-HC-95167, N01-HC-95168, N01-HC-95169, UL1-TR-000040, UL1-TR-001079, and UL1-TR-001420. MESA Family is conducted and supported by the National Heart, Lung, and Blood Institute (NHLBI) in collaboration with MESA investigators. Support is provided by grants and contracts R01HL071051, R01HL071205, R01HL071250, R01HL071251, R01HL071258, R01HL071259, by the National Center for Research Resources, Grant UL1RR033176. The provision of genotyping data was supported in part by the National Center for Advancing Translational Sciences, CTSI grant UL1TR001881, and the National Institute of Diabetes and Digestive and Kidney Disease Diabetes Research Center (DRC) grant DK063491 to the Southern California Diabetes Endocrinology Research Center.

**NHLBI TOPMed: My Life, Our Future (MLOF)**
The My Life, Our Future samples and data are made possible through the partnership of Bloodworks Northwest, the American Thrombosis and Hemostasis Network, the National Hemophilia Foundation,

and Bioverativ. We gratefully acknowledge the hemophilia treatment centers and their patients who provided biological samples and phenotypic data.

**NHLBI TOPMed: Outcome Modifying Genes in Sickle Cell Disease (OMG-SCD)**
The OMG-SCD study was administered by Marilyn J. Telen, M.D. and Allison E. Ashley-Koch, Ph.D. from Duke University Medical Center, and collection of the data set was supported by grants HL068959 and HL079915 from the National Heart, Lung, and Blood Institute (NHLBI) of the National Institute of Health (NIH).

**NHLBI TOPMed: Pediatric Cardiac Genomics Consortium's Congenital Heart Disease Biobank (PCGC-CHD)**
The Pediatric Cardiac Genomics Consortium (PCGC) program is funded by the National Heart, Lung, and Blood Institute, National Institutes of Health, U.S. Department of Health and Human Services through grants UM1HL128711, UM1HL098162, UM1HL098147, UM1HL098123, UM1HL128761, and U01HL131003.

**NHLBI TOPMed: The Pharmacogenomics of Hydroxyurea in Sickle Cell Disease (PharmHU)**
Collection of the PharmHU samples and data were supported in part by the Department of Pediatrics, Baylor College of Medicine funds, National Institutes of Health (NIH) National Institute of Diabetes and Digestive and Kidney Diseases (NIDDK) grant 1K08 DK110448-01, NIH NHLBI R01 HL069234, and U01-HL117721 funded by NHLBI. We are very grateful to the patients with sickle cell disease for their participation in PharmHU.

**NHLBI TOPMed: PIMA**
This research was supported by grants from the National Heart, Lung, and Blood Institute (NHLBI), ((5U10HL064287, 5U10HL064288, 5U10HL064295, 5U10HL064307, 5U10HL064305, 5U10HL064313, and HL080083)

**NHLBI TOPMed: The Pulmonary Hypertension and the Hypoxic Response in Sickle Cell Disease (PUSH_SCD)**
Children and adolescents of 3 to 20 years-old with SCD at three field centres, namely Howard University, Children's National Medical Center and University of Michigan were enrolled.
We thank Dr. Victor R Gordeuk and the investigators of the PUSH study and the patients who participated in the study. We also thank the PUSH clinical site team: Howard University: Victor R Gordeuk, Sergei Nekhai, Oswaldo Castro, Sohail Rana, Mehdi Nouraie, James G Taylor 6th, Juan Salomon-Andonie, Fayuan Wen, Angela Rock and Xiaomei Niu. Children National Medical Center: Caterina Minniti, Deepika Darbari, Lori Lutchman-Jones, Nitti Dham, Craig Sable, NHLBI: Mark Gladwin, Greg Kato. University of Michigan: Andrew Campbell, Gregory Ensing, Manuel Arteta, Special thanks to the volunteers who participated in the PUSH study. This project was funded with federal funds from the NHLBI, NIH. Detail description of the study was published in Haematologica. 2009 Mar;94(3):340-7, MInniti C, et al. "Elevated tricuspid regurgitant jet velocity in children and adolescents with sickle cell disease: association with hemolysis and hemoglobin oxygen desaturation."

**NHLBI TOPMed: Recipient Epidemiology and Donor Evaluation Study-III (REDS-III_Brazil)**
The Recipient Epidemiology and Donor Evaluation Study (REDS)-III was funded by NIH NHLBI contract HHSN268201100007I and conducted under the leadership of Simone Glynn (NHLBI), and principle investigators Brian Custer and Ester Sabino. We are grateful to the Brazilian sickle cell disease patients who participated in the REDS-III study and provided blood samples for whole genome sequencing as well as the REDS-III staff: Vitalant Research Institute (Shannon Kelly), University of Sao Paulo (Miriam V Flor Park, Ligia Capuani), Hemominas Belo Horizonte (Anna Barbara Proietti),

Hemominas Montes Claros (Rosimere Alfonso), Hemominas Juiz de Fora (Daniela de O. Werneck Rodrigues), Hemope (Paula Loureiro), Hemorio (Claudia Maximo).

**NHLBI TOPMed: San Antonio Family Heart Study (SAFS)**
Collection of the San Antonio Family Study data was supported in part by National Institutes of Health (NIH) grants P01 HL045522, R01 MH078143, R01 MH078111 and R01 MH083824; and whole genome sequencing of SAFS subjects was supported by U01 DK085524 and R01 HL113323. We are very grateful to the participants of the San Antonio Family Study for their continued involvement in our research programs.

**NHLBI TOPMed: Study of African Americans, Asthma, Genes and Environment (SAGE)**
The Study of African Americans, Asthma, Genes and Environments (SAGE) was supported by by the National Heart, Lung, and Blood Institute of the National Institute of Health (NIH) grants R01HL117004 and X01HL134589; study enrollment supported by the Sandler Family Foundation, the American Asthma Foundation, the RWJF Amos Medical Faculty Development Program, Harry Wm. and Diana V. Hind Distinguished Professor in Pharmaceutical Sciences II. The SAGE study collaborators include Harold J. Farber, Texas Children's Hospital; Emerita Brigino-Buenaventura, Kaiser Permanente; Michael A. LeNoir, Bay Area Pediatrics; Kelley Meade, UCSF Benioff Children’s Hospital, Oakland; Luisa N. Borrell, City University of New York; Adam Davis, UCSF Benioff Children’s Hospital, Oakland and Fred Lurmann, Sonoma Technologies, Inc. The authors acknowledge the families and patients for their participation and thank the numerous health care providers and community clinics for their support and participation in SAGE. In particular, the authors thank study coordinator Sandra Salazar; the recruiters who obtained the data: Lisa Caine, Elizabeth Castellanos, Brenda Lopez, MD, Shahdad Saeedi; and the lab researcher Celeste Eng who processed the biospecimens.

**NHLBI TOPMed: Study of Asthma Phenotypes & Pharmacogenomic Interactions by Race-Ethnicity (SAPPHIRE_asthma)**
The SAPPHIRE cohort was supported by grant funding from the Fund for Henry Ford Hospital, the American Asthma Foundation, and the following institutes of the National Institutes of Health: the National Heart Lung and Blood Institute (R01HL141845, R01HL118267, X01HL134589, R01HL079055), the National Institute of Allergy and Infectious Diseases (R01AI079139, R01AI061774), and the National Institute of Diabetes and Digestive and Kidney Diseases (R01DK113003, R01DK064695).

**NHLBI TOPMed: Genetics of Sarcoidosis in African Americans (Sarcoidosis)**
Supported by the National Institutes of Health under Grant R01HL113326-05, P30 GM110766-01, and U54GM104938-06.

**NHLBI TOPMed: Severe Asthma Research Program (SARP)**
The authors acknowledge the contributions of the study coordinators and staff at each of the clinical centers and the Data Coordinating Center as well as all the study participants that have been integral to the success of the NHLBI Severe Asthma Research Program (funded by U10 HL109164, U10 HL109257, U10 HL109146, U10 HL109172, U10 HL109250, U10 HL109250, U10 HL109250, U10 HL109168, U10 HL109152, U10 HL109086).

**NHLBI TOPMed: Genome-wide Association Study of Adiposity in Samoans (SAS)**
Financial support from the U.S. National Institutes of Health Grants R01-HL093093 and R01HL133040. We acknowledge the assistance of the Samoa Ministry of Health and the Samoa Bureau of Statistics for their guidance and support in the conduct of this study. We thank the local village officials for their help and the participants for their generosity. The following publication describes the origin of the dataset: Hawley NL, Minster RL, Weeks DE, Viali S, Reupena MS, Sun G, Cheng H, Deka R, McGarvey ST.

Prevalence of Adiposity and Associated Cardiometabolic Risk Factors in the Samoan Genome-Wide Association Study. Am J Human Biol 2014. 26: 491-501. DOI: 10.1002/jhb.22553. PMID: 24799123.

**NHLBI TOPMed: Rare Variants for Hypertension in Taiwan Chinese (THRV)**
The Rare Variants for Hypertension in Taiwan Chinese (THRV) is supported by the National Heart, Lung, and Blood Institute (NHLBI) grant (R01HL111249) and its participation in TOPMed is supported by an NHLBI supplement (R01HL111249-04S1). THRV is a collaborative study between Washington University in St. Louis, LA BioMed at Harbor UCLA, University of Texas in Houston, Taichung Veterans General Hospital, Taipei Veterans General Hospital, Tri-Service General Hospital, National Health Research Institutes, National Taiwan University, and Baylor University. THRV is based (substantially) on the parent SAPPHIRe study, along with additional population-based and hospital-based cohorts. SAPPHIRe was supported by NHLBI grants (U01HL54527, U01HL54498) and Taiwan funds, and the other cohorts were supported by Taiwan funds.

**NHLBI TOPMed: The Vanderbilt AF Ablation Registry (VAFAR)**
The research reported in this article was supported by grants from the American Heart Association to Dr. Shoemaker (11CRP742009), Dr. Darbar (EIA 0940116N), and grants from the National Institutes of Health (NIH) to Dr. Darbar (R01 HL092217), and Dr. Roden (U19 HL65962, and UL1 RR024975). The project was also supported by a CTSA award (UL1 TR00045) from the National Center for Advancing Translational Sciences. Its contents are solely the responsibility of the authors and do not necessarily represent the official views of the National Center for Advancing Translational Sciences or the NIH.

**NHLBI TOPMed: The Vanderbilt Atrial Fibrillation Registry (VU_AF)**
The research reported in this article was supported by grants from the American Heart Association to Dr. Darbar (EIA 0940116N), and grants from the National Institutes of Health (NIH) to Dr. Darbar (HL092217), and Dr. Roden (U19 HL65962, and UL1 RR024975). This project was also supported by CTSA award (UL1TR000445) from the National Center for Advancing Translational Sciences. Its contents are solely the responsibility of the authors and do not necessarily represent the official views of the National Center for Advancing Translational Sciences of the NIH.

**NHLBI TOPMed: Treatment of Pulmonary Hypertension and Sickle Cell Disease With Sildenafil Therapy (Walk-PHaSST)**
We thank Dr. Mark Gladwin and the investigators of the Walk-PHasst study and the patients who participated in the study. We also thanks the walk-PHaSST clinical site team: Albert Einstein College of Medicine: Jane Little and Verlene Davis; Columbia University: Robyn Barst, Erika Rosenzweig, Margaret Lee and Daniela Brady; UCSF Benioff Children's Hospital Oakland: Claudia Morris, Ward Hagar, Lisa Lavrisha, Howard Rosenfeld, and Elliott Vichinsky; Children’s Hospital of Pittsburgh of UPMC: Regina McCollum; Hammersmith Hospital, London: Sally Davies, Gaia Mahalingam, Sharon Meehan, Ofelia Lebanto, and Ines Cabrita; Howard University: Victor Gordeuk, Oswaldo Castro, Onyinye Onyekwere,, Alvin Thomas, Gladys Onojobi, Sharmin Diaz, Margaret Fadojutimi-Akinsiku, and Randa Aladdin; Johns Hopkins University: Reda Girgis, Sophie Lanzkron and Durrant Barasa; NHLBI: Mark Gladwin, Greg Kato, James Taylor, Vandana Sachdev, Wynona Coles, Catherine Seamon, Mary Hall, Amy Chi, Cynthia Brenneman, Wen Li, and Erin Smith; University of Colorado: Kathryn Hassell, David Badesch, Deb McCollister and Julie McAfee; University of Illinois at Chicago: Dean Schraufnagel, Robert Molokie, George Kondos, Patricia Cole-Saffold, and Lani Krauz; National Heart & Lung Institute, Imperial College London: Simon Gibbs. Thanks also to the data coordination center team from Rho, Inc.: Nancy Yovetich, Rob Woolson, Jamie Spencer, Christopher Woods, Karen Kesler, Vickie Coble, and Ronald W. Helms. We also thank Dr. Yingze Zhang for directing the Walk-PHasst repository and Dr. Mehdi Nouraie for maintaining the Walk-PHasst database and Dr. Jonathan Goldsmith as a NIH program director for this study. Special thanks to the volunteers who participated in the Walk-PHaSST study. This project was funded with federal funds from the NHLBI, NIH, Department of Health and Human Services,

under contract HHSN268200617182C. This study is registered at www.clinicaltrials.gov as NCT00492531. Detail description of the study was published in Blood, 2011 118:855-864, Machado et al "Hospitalization for pain in patients with sickle cell disease treated with sildenafil for elevated TRV and low exercise capacity".

**NHLBI TOPMed: Novel Risk Factors for the Development of Atrial Fibrillation in Women (WGHS)**
The WGHS is supported by the National Heart, Lung, and Blood Institute (HL043851 and HL080467) and the National Cancer Institute (CA047988 and UM1CA182913). The most recent cardiovascular endpoints were supported by ARRA funding HL099355.

**NHLBI TOPMed: Women's Health Initiative (WHI)**
The WHI program is funded by the National Heart, Lung, and Blood Institute, National Institutes of Health, U.S. Department of Health and Human Services through contracts HHSN268201600018C, HHSN268201600001C, HHSN268201600002C, HHSN268201600003C, and HHSN268201600004C. This manuscript was prepared in collaboration with investigators of the WHI, and has been reviewed and/or approved by the Women's Health Initiative (WHI). The short list of WHI investigators can be found at https://www.whi.org/researchers/Documents%20%20Write%20a%20Paper/WHI%20Investigator%20Short%20List.pdf.

**Other Acknowledgements**

Rebecca NA Keener was supported in part by NIH/NIGMS grant 5K12GM123914.
Tyne W Miller-Fleming was supported in part by grant(s) T32 HG008341.
Marios NA Arvanitis was supported in part by grant(s) T32HL007227.
Lucas NA Barwick was supported in part by NHLBI Contract HHSN2682016000021.
John NA Blangero was supported in part by grant(s) R01 Hl113323, U01 DK085524, P01 HL045522, R01 MH078143, R01 MH078111; R01 MH083824.
Esteban G Burchard was supported in part by the Sandler Family Foundation, American Asthma Foundation, RWJF Amos Medical Faculty Development Program, Harry Wm. and Diana V. Hind Distinguished Professor in Pharmaceutical Sciences II, and grants U01HL138626, R01HL117004, R01HL128439, R01HL135156, X01HL134589, R01HL141992, R01HL141845, U01HG009080, R01ES015794, R21ES24844, P60MD006902, R01MD010443, RL5GM118984, R56MD013312, R01HD085993, 24RT-0025 and 27IR-0030.
Juan C Celedon was supported in part by grant(s) R01 HL117191, R01 MD011764, R01 HL119952.
Dawn L DeMeo was supported in part by grant(s) PO1 HL114501 and PO1 HL132825.
Stefan Kääb is supported by the DZHK (German Centre for Cardiovascular Research), partner site: Munich Heart Alliance, Munich, Germany, and by the Munich Center of Health Sciences (MC Health) as part of LMUinnovativ.
Laura M Raffield was supported in part by grant(s) T32 HL129982.
Daniel E Weeks was supported in part by grant(s) R01HL093093, R01HL1333040.
L. Keoki NA Williams was supported in part by grant(s) R01AI061774, R01AI079139, R01HL079055, R01HL118267, R01HL141845, R01DK064695, R01DK113003, the American Asthma Foundation, and the Fund for Henry Ford Hospital.
Brian E Cade was supported in part by grant(s) K01-HL135405.
Zhanghua NA Chen was supported in part by grant(s) R00ES027870.
Michael H Cho was supported in part by NHLBI grants R01HL113264, R01HL137927, and R01HL135142.
Joanne E Curran was supported in part by grant(s) R01 Hl113323, U01 DK085524, P01 HL045522, R01 MH078143, R01 MH078111; R01 MH083824.
Eimear E Kenny was supported in part by grant(s) R01HL104608; U01HG009610; R01DK110113; U01HG009080; UM1HG0089001.
Ryan L Minster was supported in part by grant(s) R01HL093093, R01HL1333040.
Marquitta J. White was supported in part by grant(s) K01HL140218.

Scott T Weiss was supported in part by grant(s) R37 HL066289 and P01 HL132825.
Ramachandran S Vasan was supported in part by Contracts NO1-HC-25195, HHSN268201500001I and 75N92019D00031 from the National Heart, Lung and Blood Institute and grant supplement R01 HL092577-06S1; Dr. Vasan is supported in part by the Evans Medical Foundation and the Jay and Louis Coffman Endowment from the Department of Medicine, Boston University School of Medicine.
Moritz F Sinner is supported by the DZHK (German Centre Cardiovascular Research), partner site: Munich Heart Alliance, Munich, Germany
Edwin K Silverman was supported in part by grant(s) U01HL089856, P01HL114501, R01HL147148, R01HL137927.
Wayne H-H Sheu was supported in part by grant(s) MOST 107-2314B-075A-001-MY3.
Susan NA Redline was supported in part by grant(s) R35-HL135818.
Juan M Peralta was supported in part by grant(s) R01 Hl113323, U01 DK085524, P01 HL045522, R01 MH078143, R01 MH078111; R01 MH083824.
Stephen T McGarvey was supported in part by grant(s) R01HL093093, R01HL1333040.
Angel C.Y. Mak was supported in part by grant(s) R01MD010443, U01HG009080, R01HL128439, R01HL135156, R01HL141992, R01HL141845, R01HL117004.
Ruth JF Loos was supported in part by grant(s) R01DK110113; R01 DK107786; R01 HL142302.
Rajesh NA Kumar was supported in part by grant(s) UG1 HL139125, U01HL138626-01A1, HHSN2752011300013C, U19AR06952, R01AI127695.
Frank D Gilliland was supported in part by the Hastings Foundation.
Bruce Gelb was supported in part by grants(s) R01 HL098123.
Goncalo NA Abecasis was supported in part by grant(s) HHSN268201800002I, OT1HL14248, U24HG008956.
James G Wilson was supported in part by grant(s) U54GM115428.
Alexander P Reiner was supported in part by grant(s) R01HL130733, R01HL116446.

The Analysis Commons was funded by R01HL131136.

**TOPMed Consortium Members:**
Namiko Abe, Gonçalo Abecasis, Francois Aguet, Christine Albert, Laura Almasy, Alvaro Alonso, Seth Ament, Peter Anderson, Pramod Anugu, Deborah Applebaum-Bowden, Kristin Ardlie, Dan Arking, Donna K Arnett, Allison Ashley-Koch, Stella Aslibekyan, Tim Assimes, Paul Auer, Dimitrios Avramopoulos, John Barnard, Kathleen Barnes, R. Graham Barr, Emily Barron-Casella, Lucas Barwick, Terri Beaty, Gerald Beck, Diane Becker, Lewis Becker, Rebecca Beer, Amber Beitelshees, Emelia Benjamin, Takis Benos, Marcos Bezerra, Larry Bielak, Joshua Bis, Thomas Blackwell, John Blangero, Eric Boerwinkle, Donald W. Bowden, Russell Bowler, Jennifer Brody, Ulrich Broeckel, Jai Broome, Karen Bunting, Esteban Burchard, Carlos Bustamante, Erin Buth, Brian Cade, Jonathan Cardwell, Vincent Carey, Cara Carty, Richard Casaburi, James Casella, Peter Castaldi, Mark Chaffin, Christy Chang, Yi-Cheng Chang, Daniel Chasman, Sameer Chavan, Bo-Juen Chen, Wei-Min Chen, Yii-Der Ida Chen, Michael Cho, Seung Hoan Choi, Lee-Ming Chuang, Mina Chung, Ren-Hua Chung, Clary Clish, Suzy Comhair, Matthew Conomos, Elaine Cornell, Adolfo Correa, Carolyn Crandall, James Crapo, L. Adrienne Cupples, Joanne Curran, Jeffrey Curtis, Brian Custer, Coleen Damcott, Dawood Darbar, Sayantan Das, Sean David, Colleen Davis, Michelle Daya, Mariza de Andrade, Lisa de las Fuentes, Michael DeBaun, Ranjan Deka, Dawn DeMeo, Scott Devine, Qing Duan, Ravi Duggirala, Jon Peter Durda, Susan Dutcher, Charles Eaton, Lynette Ekunwe, Adel El Boueiz, Patrick Ellinor, Leslie Emery, Serpil Erzurum, Charles Farber, Tasha Fingerlin, Matthew Flickinger, Myriam Fornage, Nora Franceschini, Chris Frazar, Mao Fu, Stephanie M. Fullerton, Lucinda Fulton, Stacey Gabriel, Weiniu Gan, Shanshan Gao, Yan Gao, Margery Gass, Bruce Gelb, Xiaoqi (Priscilla) Geng, Mark Geraci, Soren Germer, Robert Gerszten, Auyon Ghosh, Richard Gibbs, Chris Gignoux, Mark Gladwin, David Glahn, Stephanie Gogarten, Da-Wei Gong, Harald Goring, Sharon Graw, Daniel Grine, C. Charles Gu, Yue Guan, Xiuqing Guo, Namrata Gupta, Jeff Haessler, Michael Hall, Daniel Harris, Nicola L. Hawley, Jiang He, Ben Heavner, Susan Heckbert, Ryan Hernandez, David Herrington, Craig Hersh, Bertha Hidalgo, James Hixson, Brian Hobbs, John Hokanson, Elliott Hong, Karin Hoth, Chao (Agnes) Hsiung, Yi-Jen Hung, Haley Huston, Chii Min Hwu, Marguerite Ryan Irvin, Rebecca Jackson, Deepti Jain, Cashell Jaquish, Min A Jhun, Jill Johnsen, Andrew Johnson, Craig Johnson, Rich Johnston, Kimberly Jones, Hyun Min Kang, Robert Kaplan, Sharon Kardia, Sekar Kathiresan, Shannon Kelly, Eimear Kenny, Michael Kessler, Alyna Khan, Wonji Kim, Greg Kinney, Barbara Konkle, Charles Kooperberg, Holly Kramer, Christoph Lange, Ethan Lange, Leslie Lange, Cathy Laurie, Cecelia Laurie, Meryl LeBoff, Jiwon Lee, Seunggeun Shawn Lee, Wen-Jane Lee, Jonathon LeFaive, David Levine, Dan Levy, Joshua Lewis, Xiaohui Li, Yun Li, Henry Lin, Honghuang Lin, Keng Han Lin, Xihong Lin, Simin Liu, Yongmei Liu, Yu Liu, Ruth J.F. Loos, Steven Lubitz, Kathryn Lunetta, James Luo, Michael Mahaney, Barry Make, Ani Manichaikul, JoAnn Manson, Lauren Margolin, Lisa Martin, Susan Mathai, Rasika Mathias, Susanne May, Patrick McArdle, Merry-Lynn McDonald, Sean McFarland, Stephen McGarvey, Daniel McGoldrick, Caitlin McHugh, Hao Mei, Luisa Mestroni, Deborah A Meyers, Julie Mikulla, Nancy Min, Mollie Minear, Ryan L Minster, Braxton D. Mitchell, Matt Moll, May E. Montasser, Courtney Montgomery, Arden Moscati, Solomon Musani, Stanford Mwasongwe, Josyf C Mychaleckyj, Girish Nadkarni, Rakhi Naik, Take Naseri, Pradeep Natarajan, Sergei Nekhai, Sarah C. Nelson, Bonnie Neltner, Deborah Nickerson, Kari North, Jeff O'Connell, Tim O'Connor, Heather Ochs-Balcom, David Paik, Nicholette Palmer, James Pankow, George Papanicolaou, Afshin Parsa, Juan Manuel Peralta, Marco Perez, James Perry, Ulrike Peters, Patricia Peyser, Lawrence S Phillips, Toni Pollin, Wendy Post, Julia Powers Becker, Meher Preethi Boorgula, Michael Preuss, Bruce Psaty, Pankaj Qasba, Dandi Qiao, Zhaohui Qin, Nicholas Rafaels, Laura Raffield, Vasan S. Ramachandran , D.C. Rao, Laura Rasmussen-Torvik, Aakrosh Ratan, Susan Redline, Robert Reed, Elizabeth Regan, Alex Reiner, Muagututi‘a Sefuiva Reupena, Ken Rice, Stephen Rich, Dan Roden, Carolina Roselli, Jerome Rotter, Ingo Ruczinski, Pamela Russell, Sarah Ruuska, Kathleen Ryan, Ester Cerdeira Sabino, Danish Saleheen, Shabnam Salimi, Steven Salzberg, Kevin Sandow, Vijay G. Sankaran, Christopher Scheller, Ellen Schmidt, Karen Schwander, David Schwartz, Frank Sciurba, Christine Seidman, Jonathan Seidman, Vivien Sheehan, Stephanie L. Sherman, Amol Shetty, Aniket Shetty, Wayne Hui-Heng Sheu, M. Benjamin Shoemaker, Brian Silver, Edwin Silverman, Jennifer Smith, Josh Smith, Nicholas Smith, Tanja Smith, Sylvia Smoller, Beverly

Snively, Michael Snyder, Tamar Sofer, Nona Sotoodehnia, Adrienne M. Stilp, Garrett Storm, Elizabeth Streeten, Jessica Lasky Su, Yun Ju Sung, Jody Sylvia, Adam Szpiro, Carole Sztalryd, Daniel Taliun, Hua Tang, Margaret Taub, Kent D. Taylor, Matthew Taylor, Simeon Taylor, Marilyn Telen, Timothy A. Thornton, Machiko Threlkeld, Lesley Tinker, David Tirschwell, Sarah Tishkoff, Hemant Tiwari, Catherine Tong, Russell Tracy, Michael Tsai, Dhananjay Vaidya, David Van Den Berg, Peter VandeHaar, Scott Vrieze, Tarik Walker, Robert Wallace, Avram Walts, Fei Fei Wang, Heming Wang, Karol Watson, Daniel E. Weeks, Bruce Weir, Scott Weiss, Lu-Chen Weng, Jennifer Wessel, Cristen Willer, Kayleen Williams, L. Keoki Williams, Carla Wilson, James Wilson, Quenna Wong, Joseph Wu, Huichun Xu, Lisa Yanek, Ivana Yang, Rongze Yang, Norann Zaghloul, Maryam Zekavat, Yingze Zhang, Snow Xueyan Zhao, Wei Zhao, Degui Zhi, Xiang Zhou, Xiaofeng Zhu, Michael Zody, Sebastian Zoellner

**TOPMed Hematology and Hemostasis Working Group:**
Laura Almasy, Allison Ashley-Koch, Paul Auer, Abraham Aviv, Melissa Bailey, Emily Barron-Casella, David Beame, Lewis Becker, Alexander Bick, Larry Bielak, Thomas Blackwell, John Blangero, Michael Bowers, Jennifer Brody, Pamela Burton, James Casella, Christy Chang, Han Chen, Ming-Huei Chen, Michael Cho, Jason Collins, Matthew Conomos, Adolfo Correa, Rhea Cosentino, Paul de Vries, Jennifer Dean, Pinkal Desai, Qing Duan, Connor Emdin, Nauder Faraday, Jean Feng, Regina Fisher, Annette Fitzpatrick, James Floyd, Santhi Ganesh, Brady Gaynor, Manjit Hanspal, Ross Hardison, Ben Heavner, Craig Hersh, Chani Hodonsky, Michael Honigberg, Steve Horvath, Yao Hu, Jennifer Huffman, Kruthika Raman Iyer, Deepti Jain, Sidd Jaiswal, Jill Johnsen, Andrew Johnson, Brian Joyce, Joel Kaufman, Addison Keely, Rebecca Keener, Shannon Kelly, Derek Klarin, Malgorzata Klauzinska, Barbara Konkle, Charles Kooperberg, Ethan Lange, Leslie Lange, Cathy Laurie, Cecelia Laurie, Brandon Lê, Grace Lee, Claire Leiser, Guillaume Lettre, David Levine, Dan Levy, Joshua Lewis, Bingshan Li, Yun Li, Amarise Little, Shelly-Ann Love, Rasika Mathias, Ravi Mathur, Caitlin McHugh, Karen Miga, Anna Mikhaylova, Julie Mikulla, Braxton D. Mitchell, Alanna C Morrison, Rakhi Naik, Drew Nannini, Pradeep Natarajan, Deborah Nickerson, Jeff O'Connell, Christopher O'Donnell, Nels Olson, Nathan Pankratz, Benedict Paten, James Perry, Steve Pipe, James Pirruccello, Linda Polfus, Bruce Psaty, Jennifer Anne Purnell, Laura Raffield, Alex Reiner, Stephen Rich, Shabnam Salimi, Vijay G. Sankaran, Noah Simon, Nicholas Smith, Adrienne M. Stilp, Hua Tang, Margaret Taub, Marilyn Telen, Timothy A. Thornton, Russell Tracy, Md Mesbah Uddin, Kate Wehr, Joshua Weinstock, Ellen Werner, Marsha Wheeler, Ann Whitney, Eric Whitsel, Kerri Wiggins, Lisa Yanek, Yu-Chung Yang, Maryam Zekavat, Wei Zhao, Xiuwen Zheng, Yinan Zheng

**TOPMed Structural Variation Working Group:**
Paul Auer, Melissa Bailey, Kathleen Barnes, David Beame, Michael Bowers, Harrison Brand, Ulrich Broeckel, Mark Chaisson, Lavanya Challagundla, Kei Hang Katie Chan, Seung Hoan Choi, Zechen Chong, Bradley Coe, John Cole, Ryan Collins, Matthew Conomos, Michelle Daya, Jennifer Dean, Scott Devine, Evan Eichler, Annette Fitzpatrick, Stephanie Gogarten, C. Charles Gu, Amelia Weber Hall, Ira Hall, Bob Handsaker, Ben Heavner, James Hixson, Deepti Jain, Jicai Jiang, Jill Johnsen, Brian Joyce, Goo Jun, Hyun Min Kang, Addison Keely, Spencer Kelley, Charles Kooperberg, John Lane, Cathy Laurie, Seung-been Steven Lee, David Levine, Dan Levy, Honghuang Lin, Simin Liu, Angel CY Mak, Alisa Manning, Rasika Mathias, Steve McCarroll, Julie Mikulla, Drew Nannini, Giuseppe Narzisi, Deborah Nickerson, Jeff O'Connell, Wanda O'Neal, Grier Page, Nathan Pankratz, Alexandre Pereira, Patricia Peyser, Nathan Pezant, Jennifer Anne Purnell, Aakrosh Ratan, Alex Reiner, Stephen Rich, Ingo Ruczinski, Aniko Sabo, Steven Salzberg, Jenny Schoenberg, Jonathan Seidman, Minseok Seo, Nasa Sinnott Armstrong, Albert Vernon Smith, Vinodh Srinivasasainagendra, Arvis Sulovari, Margaret Taub, Olivia Tintea, Kate Wehr, Joshua Weinstock, Marsha Wheeler, James Wilson, Huichun Xu, Sandy Zellner, Wei Zhao, Yinan Zheng, Degui Zhi, Sebastian Zoellner
